# A taxonomic revision of the genus *Hydnora* (Hydnoraceae)

**DOI:** 10.1101/2022.10.13.512068

**Authors:** Sebastian A. Hatt, Chris J. Thorogood, Jay F. Bolin, Lytton J. Musselman, Duncan D. Cameron, Olwen M. Grace

## Abstract

A systematic monograph is presented for Hydnora (Hydnoraceae), a poorly known genus of holoparasitic plants distributed across Africa, Madagascar and southern Arabia. Species of *Hydnora* are characterised by their underground habit, unusual fleshy flowers and complete absence of leaves or photosynthetic tissue. This is the first detailed monograph of the genus Hydnora since 1935 and is informed by a comprehensive survey of herbarium specimens and literature. Detailed descriptions, full synonymy, distribution maps and discussion concerning confusable taxa are provided for each species, along notes on ethnobotany, ecology and conservation. We place particular emphasis on the taxonomic value of osmophore geometry and positioning in living and dried material, which are highly consistent within species. We also provide the first detailed assessment of host range across the genus. *Hydnora hanningtonii* Rendle and *H. solmsiana* Dinter are reinstated from synonymy, and *H. bolinii* Hatt is newly described here. The infrageneric classification is reviewed and a key is provided for both living and dried material. Species are accompanied by both illustrations and photographs of living and dried material where possible.

## INTRODUCTION

*Hydnora* Thunb. is a genus of parasitic plants distributed across much of southern and eastern Africa, Madagascar, and southern Arabia (Map 1). Together with the American sister genus *Prosopanche* de Bary, *Hydnora* is holoparasitic, meaning that it is entirely dependent on a host plant for all of its carbon and nutrient intake. The two genera are placed in the family Hydnoraceae, which is nestled within the Piperales (Jost et al., 2021). Following *Cassytha* L. (Lauraceae), Hydnoraceae are the second earliest diverging lineage of extant parasitic plants (Naumann et al., 2013; Nickrent et al., 2020).

The genus is unusual in that all plants are completely devoid of photosynthetic vegetative parts, and because of their extreme morphological divergence, the plants are sometimes mistaken for a fungus at first glance (Bolin et al., 2011). Primarily a large subterranean rhizome (Tennakoon et al., 2007), *Hydnora* only breaches the surface of the soil to flower, in some cases for just a few weeks following the rains (Bolin et al., 2018). Their bizarre, fleshy flowers attract pollinating beetles with a potent, foetid odour, trapping them within their hollow reproductive chamber (Seymour et al., 2009). The beetles can escape when the flowers wither a few days later, and pollination follows (Bolin et al., 2009a). The ovary then develops into a large swollen fruit containing thousands of tiny black seeds. The fruit is sought after by jackals and other smaller mammals, hence its common name in South Africa: *jakkalskos* (Bolin et al., 2011).

Currently represented by eight species, the genus has suffered prolonged neglect and taxonomic confusion since it was described in 1775 by Carl Thunberg (Musselman & Visser, 1989; Bolin et al., 2018). A combination of a largely hypogeous habit and remote distribution, often in politically unstable regions, has meant that *Hydnora* has remained elusive to plant collectors and taxonomists (Bolin, 2009b). Furthermore, a considerable number of the specimens stored in herbaria are in relatively poor condition; many are shattered, deformed fragments of little or no taxonomic value. These factors partly explain the vague and incomplete descriptions and consequent synonymy that is characteristic of much of the research into *Hydnora* in the 19^th^ and early 20^th^ centuries. The work of Lytton Musselman and Jay Bolin over the last few decades has significantly helped to clarify some of this confusion, although considerable uncertainty remains (Musselman & Visser, 1989; Bolin et al., 2018). The lack of a published widely sampled phylogeny has also greatly hindered our understanding of the evolutionary relationships among species.

Host preference varies considerably across the genus (Musselman & Visser, 1989). Subgenus *Euhydnora* Decaisne (now subgenus *Hydnora*) in southern Africa exclusively parasitise succulent *Euphorbia* L.. Within this group, *H. visseri* Bolin, E.Maass & Musselman, *H. longicollis* (Welw.) Bolin and *H. triceps* Drege & E. Mey are restricted to one, two or a few mutually exclusive host species, while *H. africana* Thunb. has at least 12 recorded hosts (see Host Specificity and Speciation). The remaining species of *Hydnora* parasitise Fabaceae (primarily *Vachellia* Wight & Arn. and *Senegalia* Raf. species although other genera have been recorded), with the exception of *H. sinandevu* Beentje & Q. Luke which parasitises *Commiphora* Jacq. (Burseraceae) hosts. What little information exists about host preference is scattered across herbarium specimen label data, observational notes, and floristic accounts. However, these data are often unreliable as collectors rarely excavate *Hydnora* to confirm host-parasite contact. The complex pattern of host preference exhibited across the genus indicates potential co-evolution between host and parasite; like many aspects of the ecology and evolution of *Hydnora*, this remains unexamined (Bolin et al., 2011).

Distributional data are limited to the relatively small number of herbarium specimens, a few scattered observations in floristic accounts and more recently the small, but steadily growing, collection of observations recorded by digital means such as iNaturalist (https://www.inaturalist.org). Consequently, conservation assessments are scarce and often unreliable given the paucity of data to inform them (Dougherty et al., 2015). Sparse collecting, a lack of records, and poor preservation of specimens together point to significant gaps in our knowledge of the distribution of *Hydnora*. For example, a population of *H. abyssinica* was described from Nigeria just five years ago, more than 1,500 miles from the nearest other collection of the same species (Agyeno et al., 2018).

This monograph is the first comprehensive revision of the genus *Hydnora* since 1935 (Vaccaneo, 1934; Harms, 1935). In 1989, Musselman & Visser estimated that there were fewer than 100 meaningful specimens of *Hydnora* in the world’s herbaria. Now, at least 443 specimens are known to the authors, most of which were examined for this monograph. This study includes a detailed list of recorded host species, together with detailed descriptions, accompanying illustrations, distribution maps and all available ecological and ethnobotanical data for each species. These are complemented by a key for living and dried specimens. *H. solmsiana* Dinter and *H. hanningtonii* Rendle are reinstated from synonymy, and *H. bolinii* Hatt is newly described here.

## TAXONOMIC HISTORY

The taxonomic history of *Hydnora* begins with the collection of *H. africana* Thunb. in South Africa by the Swedish botanist Carl Thunberg (1775), who published it as an extraordinary new species of fungus (see illustration in Fig. 1). A year later, Erik Acharius, student of Carl Linnaeus, asserted the plant genus, *Aphyteia*, with single species *Aphyteia hydnora* [*Hydnora africana*] (Acharius & von Linné, 1776). In the decades that followed, *Aphyteia* and *Hydnora* were used somewhat interchangeably, until the early 1800s, by which time *Hydnora* had become the dominant name in use. According to International Code of Nomenclature for algae, fungi, and plants, the initial name, *Hydnora*, is valid even though it was described as a fungus, as the Code encompasses both plants and fungal nomenclature (Preamble 8, Turland et al., 2018) Indeed, it appears Thunberg was unsure whether it should be described as a plant or a fungus, as evidenced in his report to the Royal Academy of Science in Stockholm in 1775: ‘but of all that I have so far had the opportunity to see and discover, nothing has seemed to me more strange than the fungus whose description I have the honour of submitting. So strange is its composition that many would certainly doubt the existence of such a plant on the face of the earth’ (Svedelius, 1944). In the early 1800s, a small number of collections were made and deposited in European herbaria by ‘entrepreneurial collectors’ Christian Ecklon, Karl Zeyher and Johann Franz Drège (Arnold, 2021). One of these collections by Drège precipitated the description of a second species, *H. triceps* Drège & E.Mey, by Ernesto Meyer (1833), who published a small but relatively detailed report on the two accepted species.

**Fig. 1.**
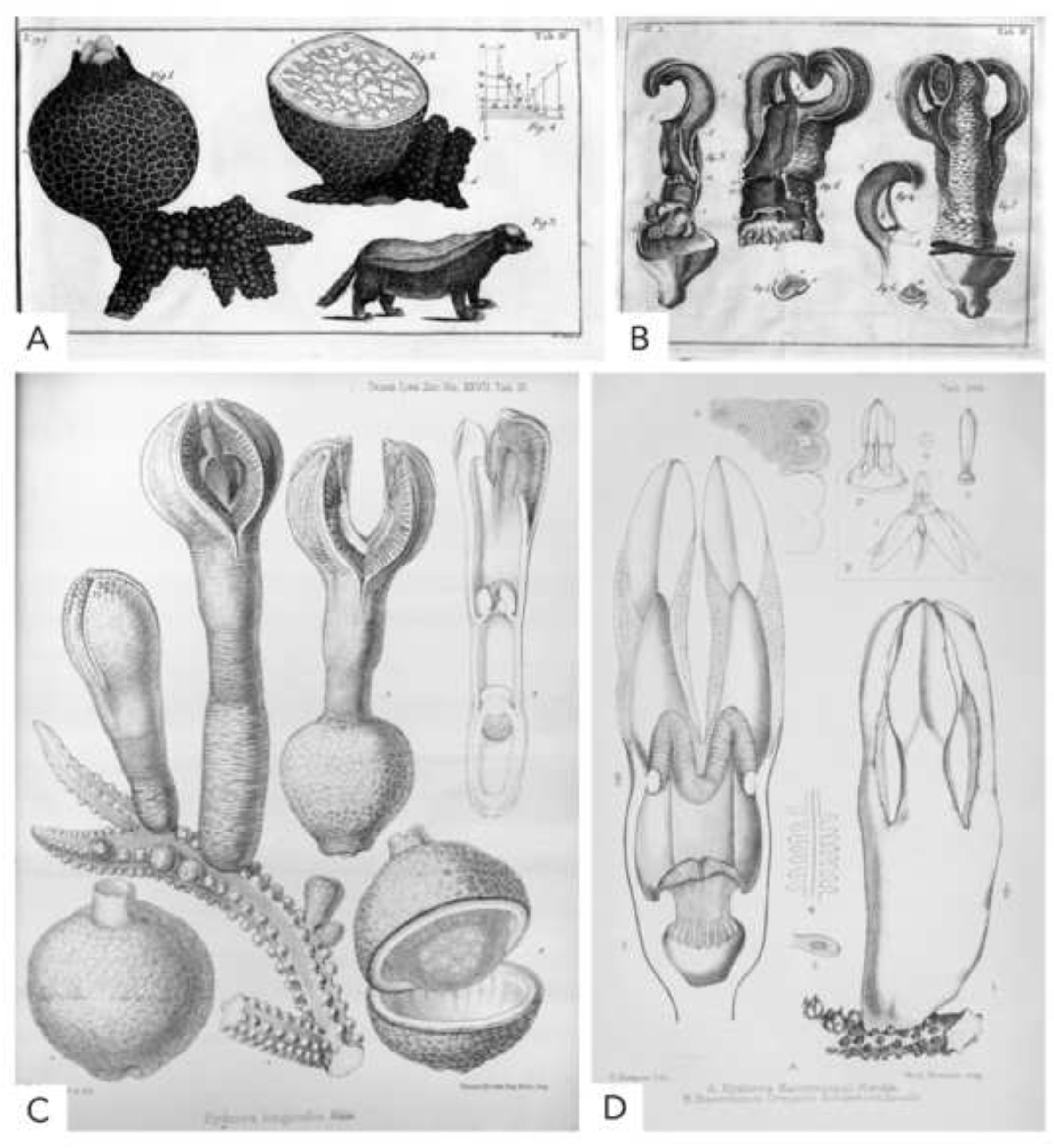
Early illustrations of *Hydnora* species. **A, B** *Hydnora africana* from Thunberg (1775); **C** *H. longicollis* from Welwitsch (1869); **D** *H. hanningtonii* from Rendle (1896).

Debate ensued over the affinities of *Hydnora*, with British botanist Robert Brown believing it to be closely related to *Rafflesia* R.Br. ex Gray (Rafflesiaceae) due to its shared trait of parasitism, while others disagreed, such as Brown’s student William Griffith (Brown, 1844; Arnold, 2021). Over the remainder of the 19^th^ century, several new species of *Hydnora* were published by European botanists including Welwitsch (1869; Fig. 1), Beccari (1871), Decaisne (1873) and Rendle (1896, Fig. 1). Decaisne made the first attempt to divide *Hydnora* into subgenera (Decaisne, 1873). He grouped those with four tepals into subgenus *Dorhyna* Decaisne and those with three into subgenus *Euhydnora* (now subgenus *Hydnora*). Many of the species during this period were described in isolation and authors rarely observed other specimens in person (Arnold, 2021). As a result, many of the descriptions share insignificant differences and the species have since been synonymised (Musselman & Visser, 1989).

The turn of the century saw an increase in floristic work, with floras by Dinter (1909), Thistleton-Dyer (1912) and Marloth (1913) consolidating *Hydnora* species in southern Africa. New species continued to be described until Vaccaneo’s exhaustive monograph (1934). In addition to describing a new species, *H. cornii* Vaccaneo, he presented detailed descriptions and ecological data. He also suggested that many species described thus far did not hold up to scrutiny. He retained Decaisne’s two subgenera, although altered the criteria such that they were based on rhizome characters rather than the variable and unreliable trait of floral merosity. Hermann Harms published another monograph a year later (1935), largely based on the foundations laid by Vaccaneo. Most notably, he extended the subgeneric classification to include subgenus *Neohydnora* Harms (*H. esculenta* Jum. & H.Perrier), and subgenus *Tricephalohydnum* Harms (*H. triceps*).

Only a few publications on *Hydnora* emerged in the decades that followed. This hiatus was lifted by a series of publications by Lytton Musselman and Johann Visser, partly triggered by the instigation of various floristic projects across Africa in the 1980s (Visser, 1981; Musselman & Visser, 1987). This culminated in a report in 1989 on the natural history and updated taxonomy of the genus (Musselman & Visser, 1989). Accounts that followed this paper, mostly published by Musselman, synonymised 11 species described from East Africa under *H. johannis* Becc. (Musselman, 1993; 1997; 2001). Beentje and Luke (2002) determined that *H. abyssinica* A.Br. was in fact the valid name for this species, although many herbarium specimens are still labelled as *H. johannis*. In the years since, Musselman’s student Jay Bolin has described two new species, *H. visseri* and *H. arabica* Bolin & Musselman (Bolin et al., 2011; 2018). In his PhD thesis, he produced a revision of subgenus *Euhydnora* (now subgenus *Hydnora*) in southern Africa, and the first molecular phylogeny for the genus, including seven species of *Hydnora* using *rpoB* and ITS regions (Bolin, 2009b). As it stands, there are eight accepted species of *Hydnora*, grouped into four subgenera, and a comprehensive monograph of the entire genus has not been updated for almost a century (Harms, 1935; Bolin et al., 2018).

## MATERIALS & METHODS

This monograph has been compiled through the examination of herbarium specimens, literature, recorded observations and fieldwork. Specimens of *Hydnora* were observed by the authors at BM, BOL, EA, GHPG, GRA, K, NBG, OBG, ODU, SAM, TAN and WIND. All specimens from other herbaria (AAU, B, BR, E, FT, G, HARG, HBG, HUH, JUHN, L, LD, LISU, M, MO, P, PRE, PSUB, RO, S, SRGH, UPS, US) were examined digitally. Species descriptions were updated with new and relevant data from herbarium specimens and the literature.

### Species Concepts

The species concept used follows the ‘morphological cluster’ concept outlined in Vorontsova & Knapp (2016): “assemblages of individuals with morphological features in common and separate from other such assemblages by correlated morphological discontinuities in a number of features” (Davis & Heywood, 1963). Although the taxonomic decisions made here are based on clear and consistent morphological characters, DNA sequence data would provide a higher level of resolution. As such, the taxonomic decisions made here will be corroborated with molecular data in the future.

### Host Specificity

A list of potential host species for each *Hydnora* species was compiled using herbarium specimen label data and literature (Table 1). Each entry is accompanied by a reference and a confidence rating, with a brief justification. These are ranked from ‘Low’, ‘Medium’ or ‘High’ based number and quality of the records in the context of all known records for the species. Host species are marked ‘High’ if they have been reported numerous times, or observed in situ by the authors; ‘Medium’ if there is a single or two reports, with the species in the expected family/genus for that *Hydnora* species (e.g., *Euphorbia* for subgenus *Hydnora*), or ‘Low’ if the host is not from the expected family/genus for that *Hydnora* species.

**Table 1.**
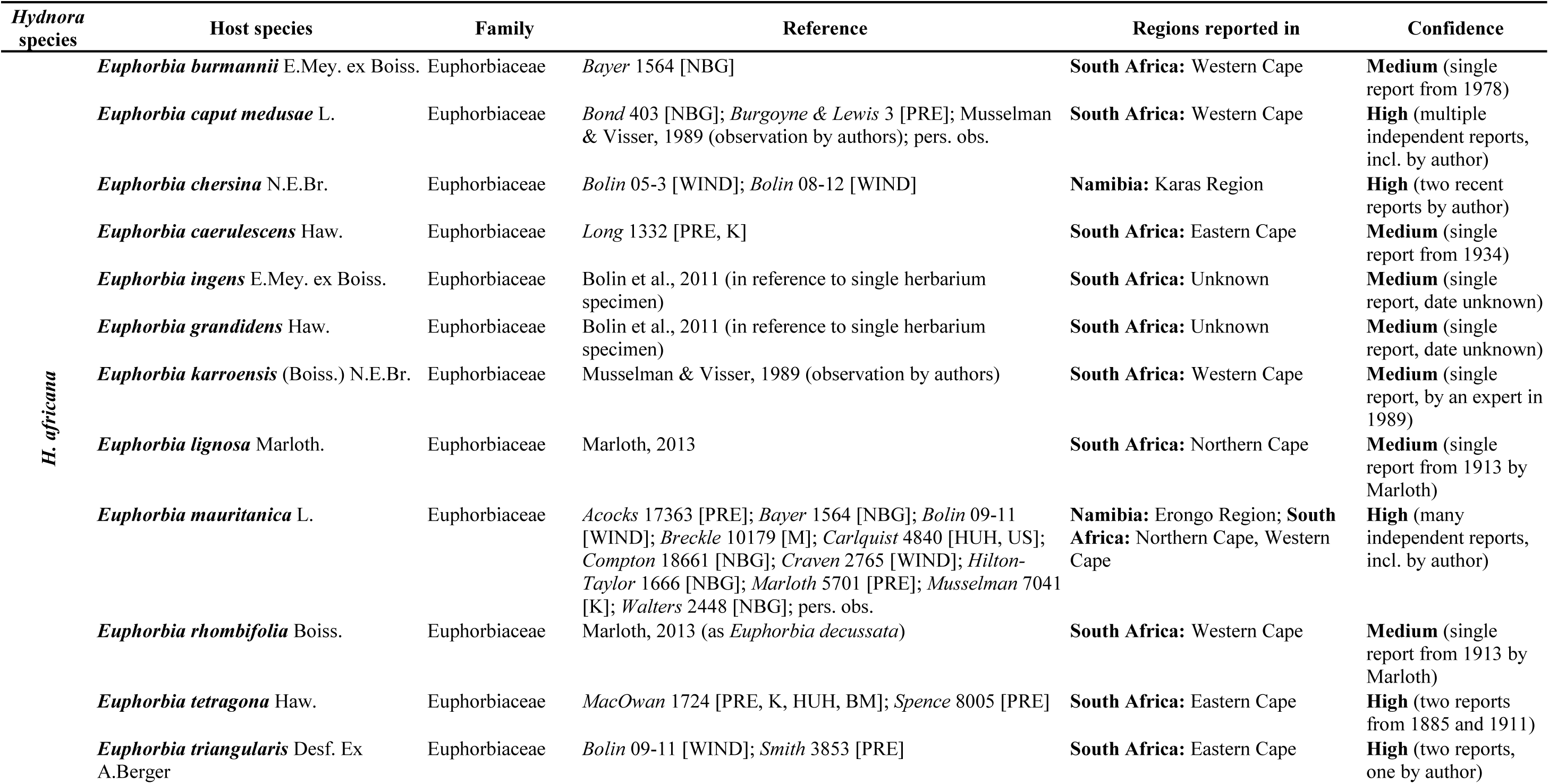
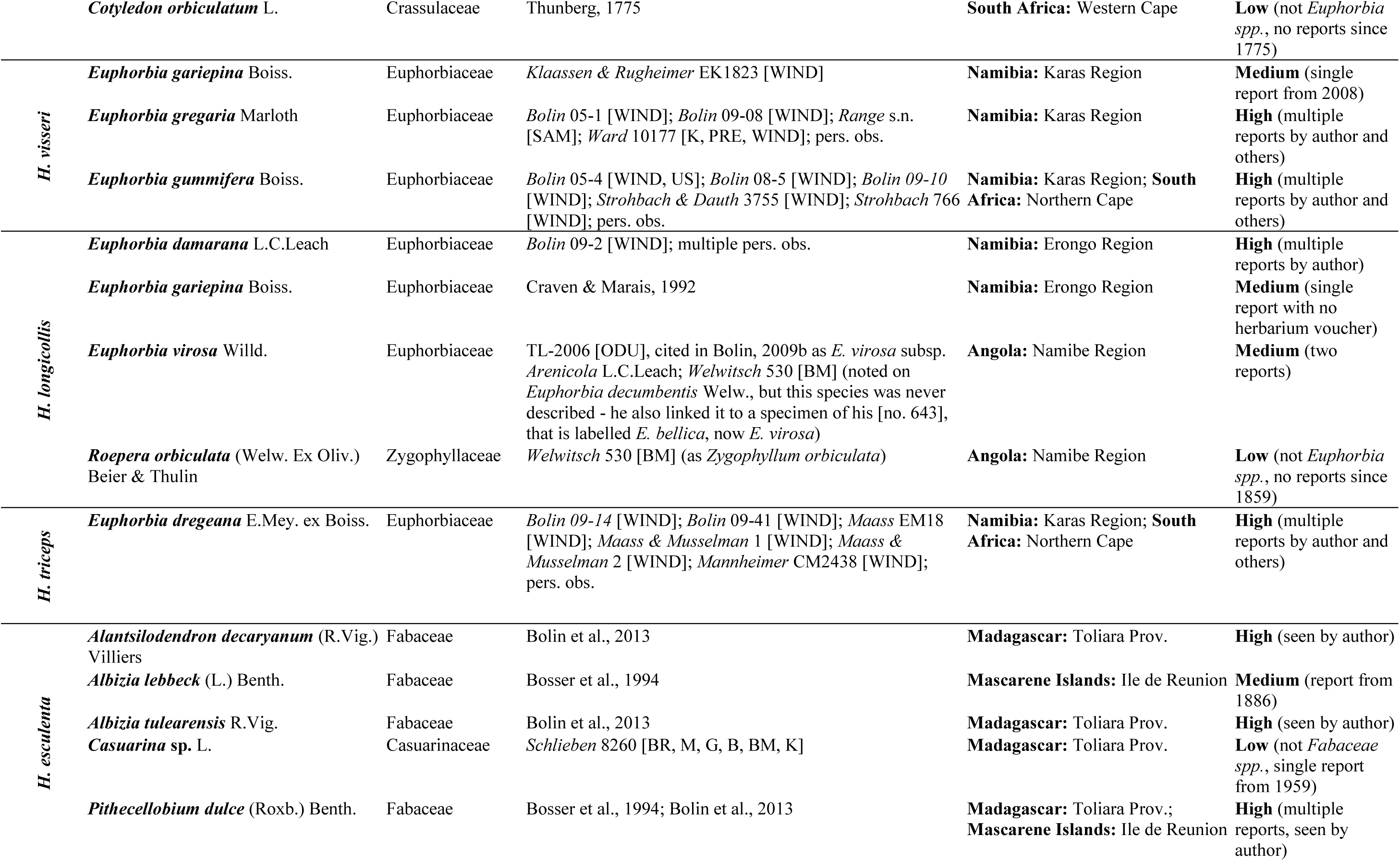
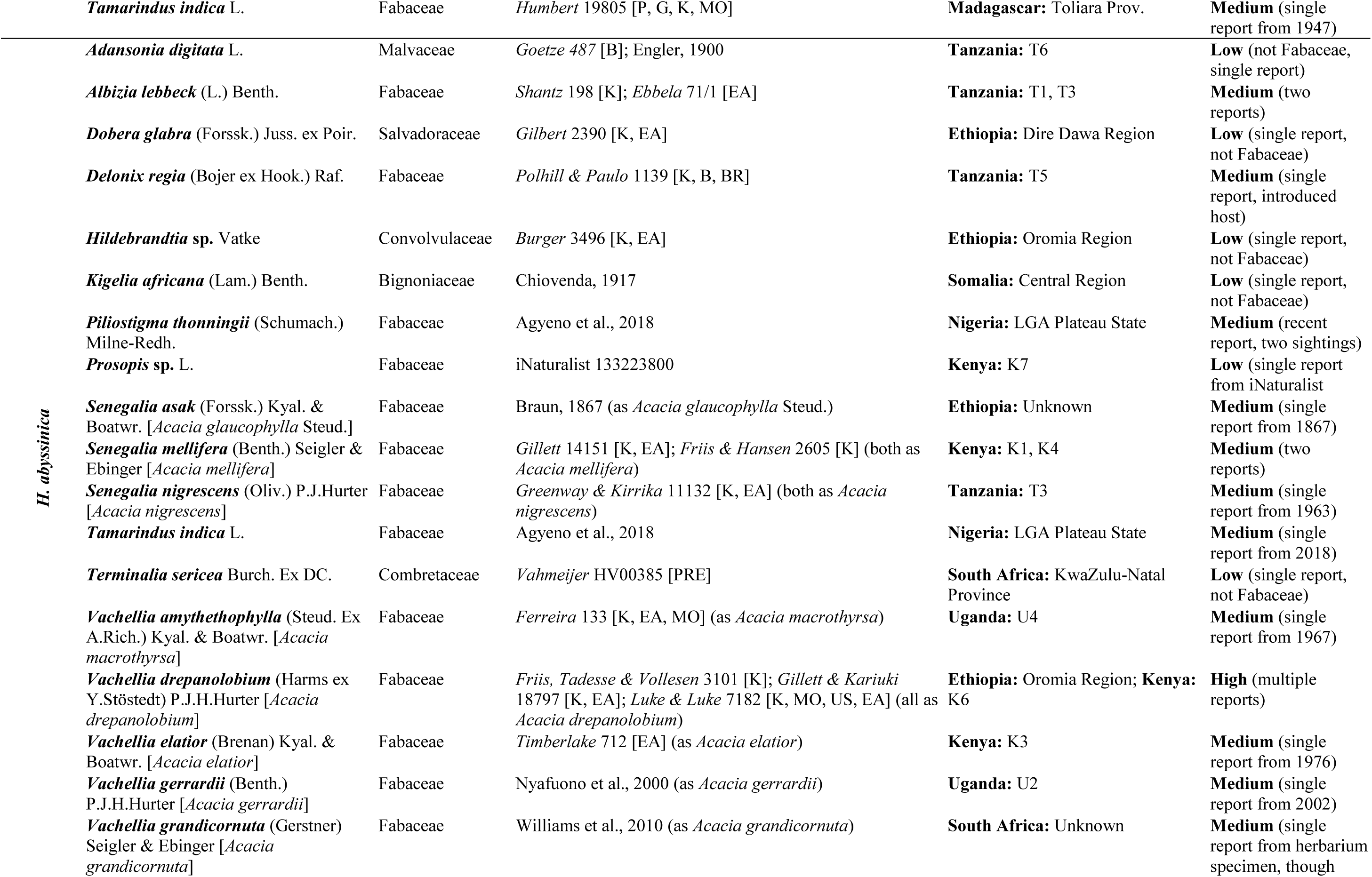
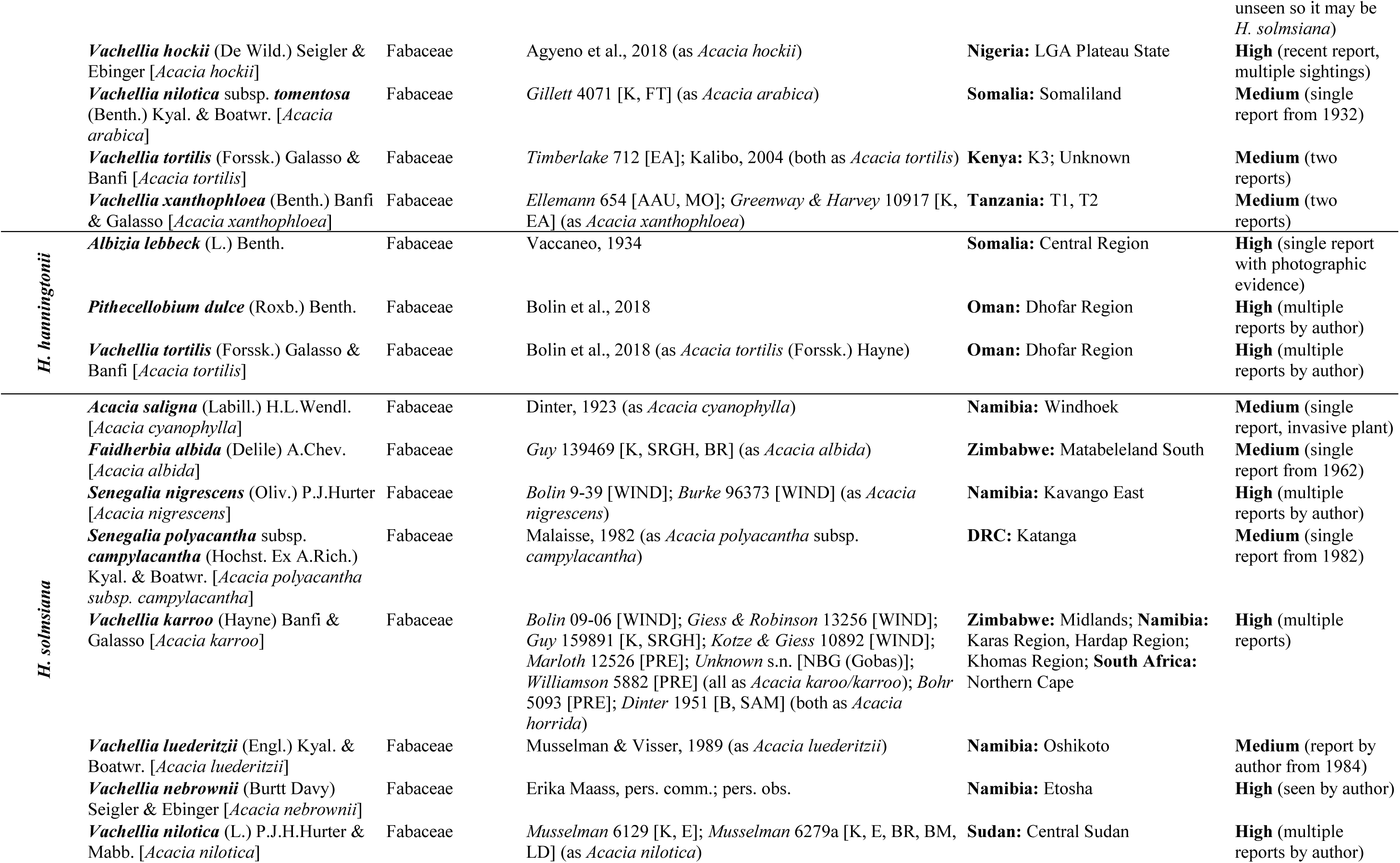
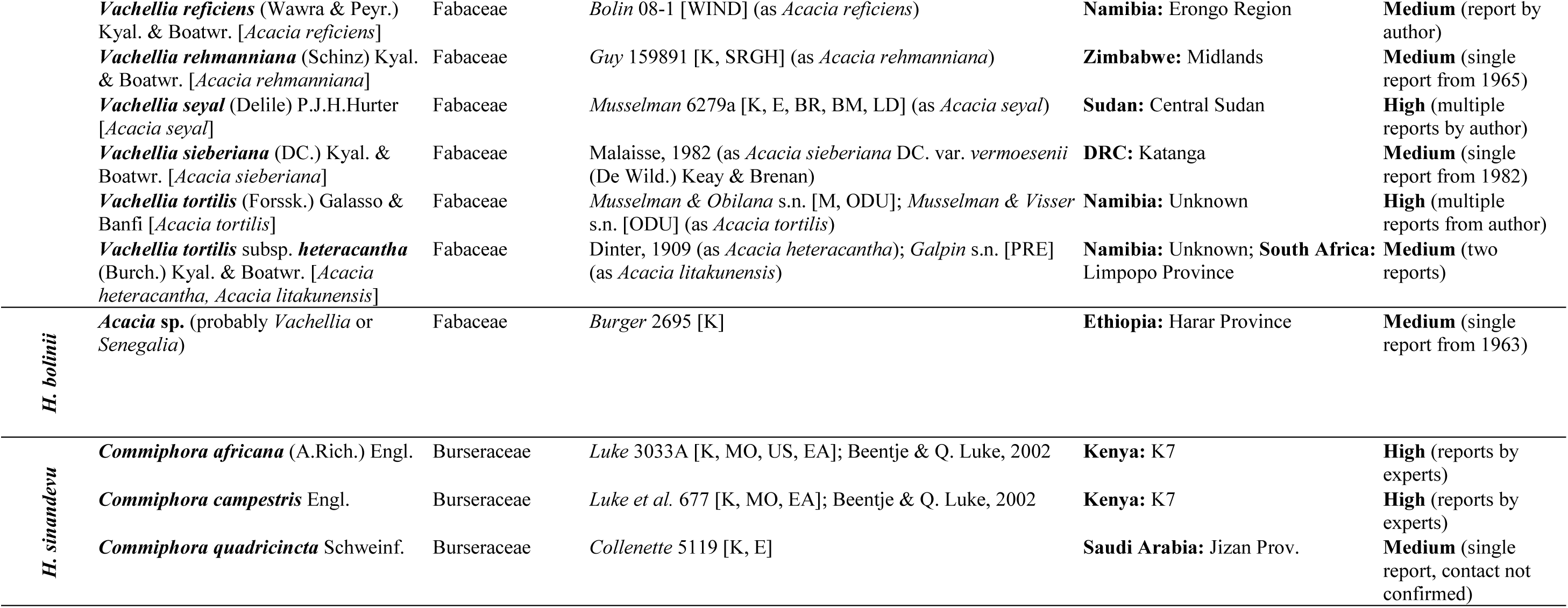
List of recorded host species for Hydnora. Each entry is accompanied by a reference and a confidence measure with a brief justification. These are ranked either ‘Low’, ‘Medium’ or ‘High’ (see Materials and Methods). They are subjective measures and should be taken as such, but will hopefully provide some indication on how valid the host record is likely to be.

### Distribution Maps

Distribution maps were assembled by combining all available data from herbarium specimens, literature and iNaturalist observations (iNaturalist, 2020). Most herbarium specimens lacked co-ordinates, and were therefore georeferenced where possible using Google Earth and collecting locality indices such as (Polhill, 1988). Point data were inputted into ArcGIS Pro 2.8, from which distributional maps were produced (ESRI, 2021). Collecting localities written in the ‘specimens examined’ sections appear as written on herbarium labels. Any spelling corrections of historic localities are put in square brackets. For specimens from Kenya, Tanzania and Uganda, the geographical system used by Flora of Tropical East Africa (Polhill, 1988) is used (e.g., K1, T1, U1).

#### Figures

Illustrations and line drawings were prepared by the author for all species, with the exception of an illustration for *Hydnora bolinii*, which lacked sufficient photographic material to produce a reliable illustration in this way and therefore specimen photographs and line drawings are supplied instead. The illustrations were prepared from collated reference material of photographs of living material and herbarium specimens.

## MORPHOLOGY

*Hydnora* possess unique morphology, such that certain characters are impossible to describe succinctly using conventional botanical terminology. This confusion is reflected in former descriptions of *Hydnora*, where the same words have been employed to describe different features. For example, the term ‘bait body’ is sometimes used to refer to the apical osmophore, and at other times the long setae that line the tepal margin in *H. africana* (Musselman & Visser, 1989; Bolin, 2009b). To overcome this obstacle, existing terms (Beentje, 2020) have been adapted here and outlined in an illustrated glossary (Fig. 2), with efforts stay as coherent with previous descriptions of *Hydnora* as possible. To ensure clarity, the terms defined here are deployed consistently throughout the monograph.

**Fig. 2.**
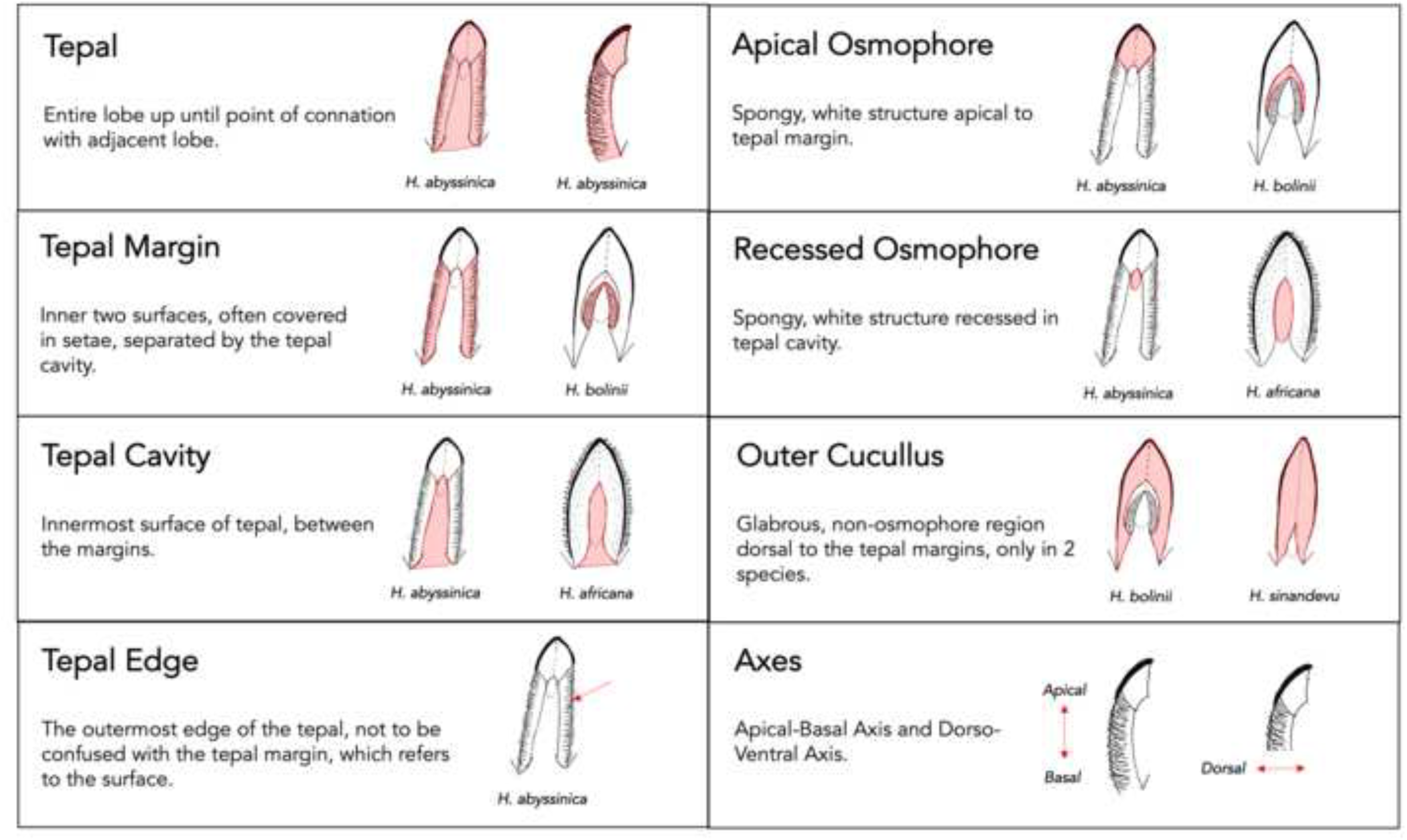
Illustrated glossary of terms used in this monograph to apply to characters on *Hydnora* tepals. Each word is accompanied by an illustrated example of how the term is applied. The bottom-right box illustrates the two floral axes referred to in the text.

*Hydnora* have fewer taxonomically valuable morphological characters than most autotrophic plants, thus putting added value on ecological traits and molecular data for identification. This is further complicated by the loss of certain key characters only available in the field, such as colour, smell, and gross morphology. Even within a single population, flowers can vary in morphology and colour; partly due to differing growing conditions or the differing stages of transformation that the flower undergoes during anthesis. However, there are a set of key characters outlined below that can be reliably used to distinguish dried specimens of almost any age, assuming both the flower and rhizome have been collected.

### Rhizome

Gross morphology and surface texture of the rhizome is inherently variable depending on environmental conditions (Fig. 3A–E). However, the positioning of the tubercles along the rhizome surface has significant taxonomic value (Fig. 3H, J). Either the tubercles are randomly distributed across the surface, with the rhizome therefore generally terete or subterete in cross-section, or they are arranged in distinct parallel lines and thus the rhizome is angular in cross-section (Fig. 3F, G). This trait reliably separates East African subgenera *Dorhyna* and *Sineseta* from the southern African subgenus *Hydnora* and the Madagascan subgenus *Neohydnora*. Beyond this trait, the rhizome has not been found to be of taxonomic value. The colour of the internal rhizome flesh can vary from pink to red to black depending on age at the time of cross-section.

**Fig. 3.**
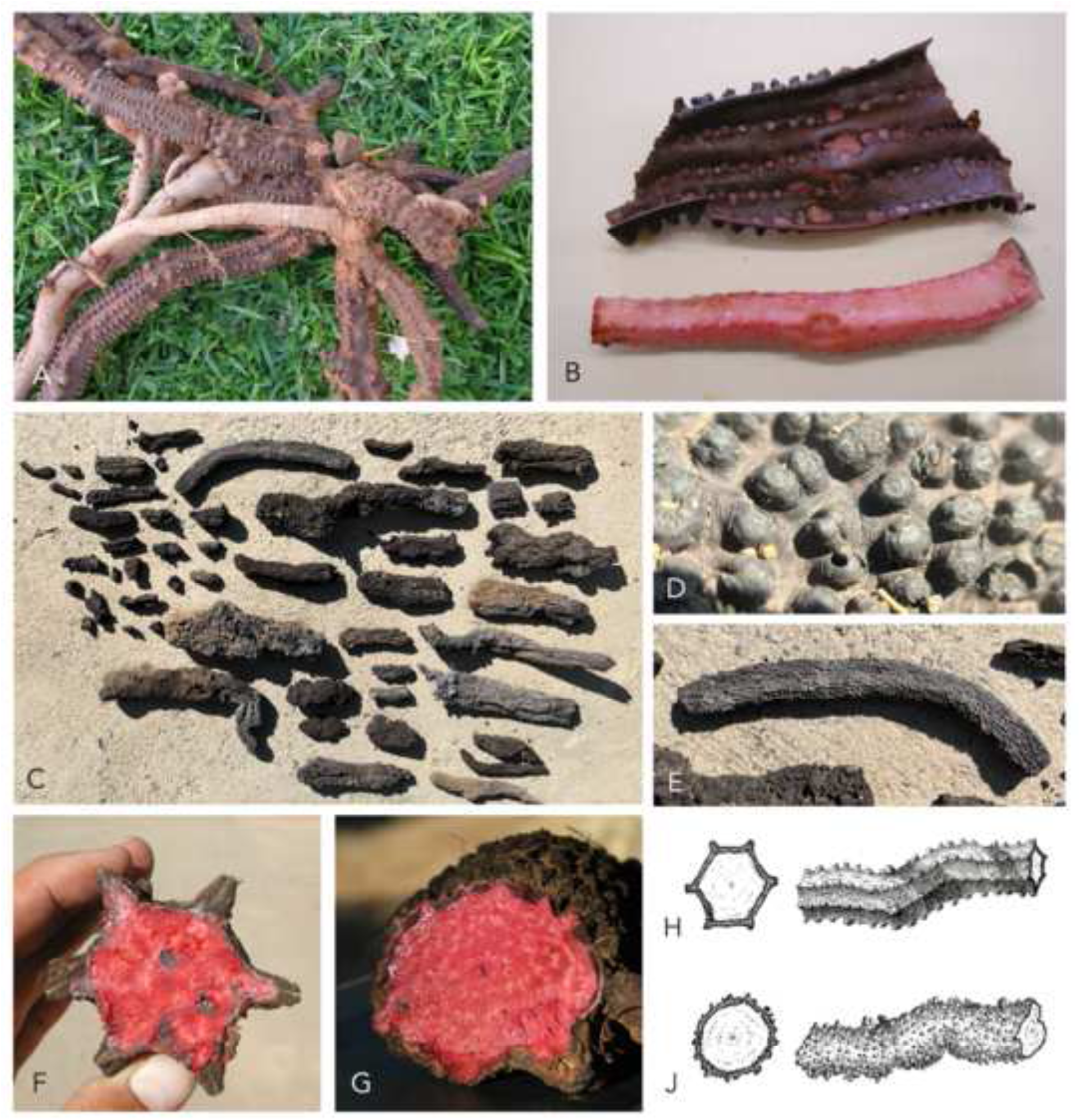
Rhizome morphology of *Hydnora*. **A** *Hydnora africana*: rhizome and associated host root; **B** *H. africana*: flesh of rhizome detached from outer skin; **C, E** *H. solmsiana*: fragments of dried rhizomes; **D** *H. solmsiana*: surface of dried rhizome; **F** *H. africana*: rhizome cross-section; **G** *H. africana*: rhizome side view; **H** angular rhizome type (subgenus *Hydnora* and *Neohydnora*); **J** terete rhizome type (subgenus *Dorhyna* and *Sineseta*). **H – J** DRAWN BY SEBASTIAN HATT.

### Flower

Gross floral anatomy is conserved across the genus (Fig. 4A–G). Floral merosity has been used in the past as a subgenus-defining character. Although it is no longer considered a reliable character due to natural phenotypic fluctuation in this trait, most individuals in subgenus *Hydnora* have three tepals, most in subgenera *Dorhyna* and *Sineseta* have four and most in subgenus *Neohydnora* have three to five. While it can be a valuable character for identification, it cannot be used in isolation. Fruit and seed morphology have not been found to vary considerably between species (Fig. 4J–N).

**Fig. 4.**
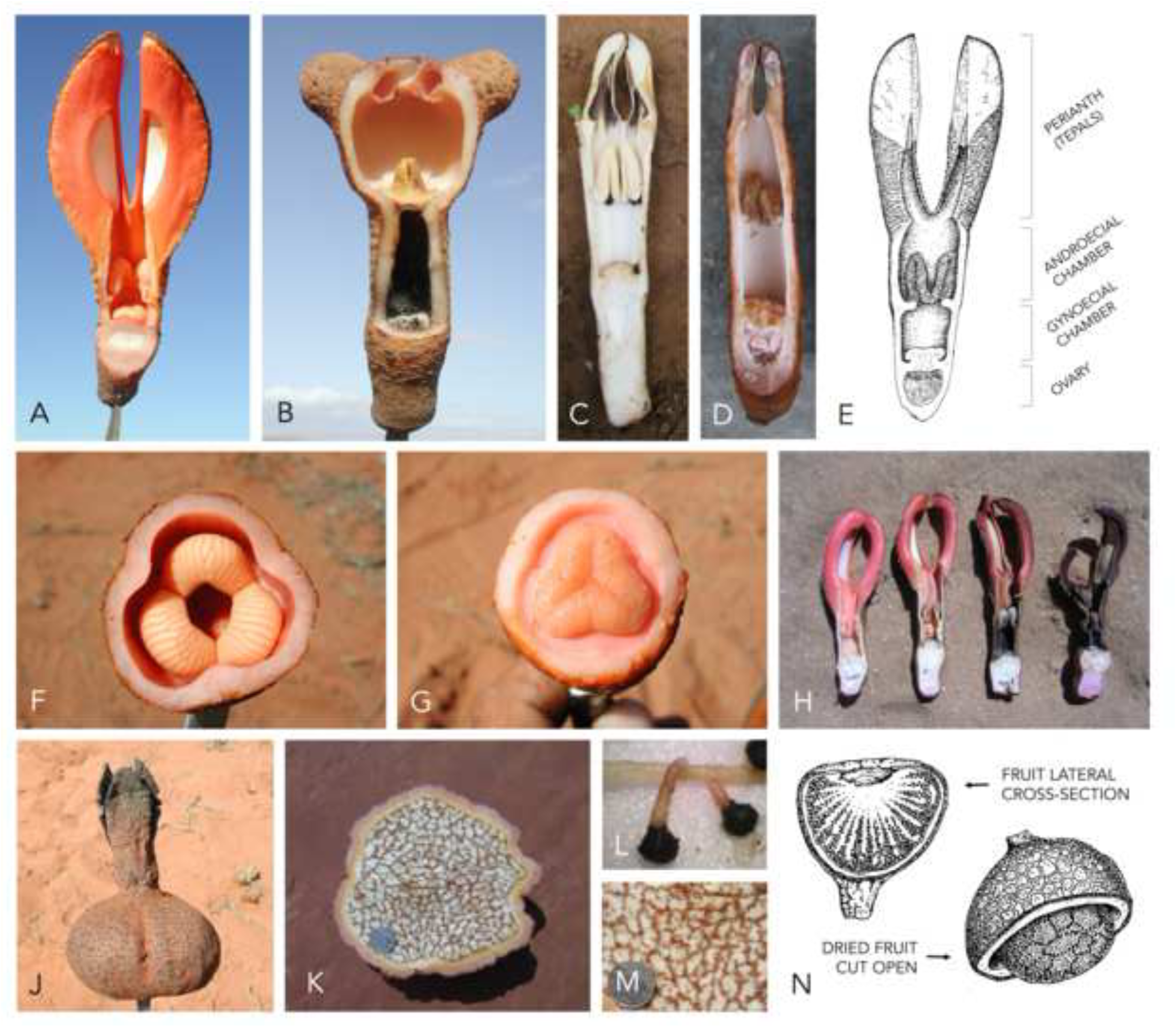
Flower morphology of *Hydnora*. **A** *Hydnora visseri* flower cross-section; **B** *H. triceps* flower cross-section; **C** *H. esculenta* flower cross-section; **D** *H. solmsiana* flower cross-section; **E** diagram of general floral structure (*H. hanningtonii* in this example); **F** androecial chamber cross-section (*H. visseri*); **G** gynoecial chamber cross-section (*H. visseri*); **H** four stages of decay of *H. visseri* flowers, fresh (left) to dry (right); **J** fruit (*H. triceps*); **K, M** fruit cross-section (*H. triceps*); **L** germinating seeds (*H. triceps*); **N** diagram of general fruit structure. **E & N** DRAWN BY SEBASTIAN HATT.

### Osmophore

‘Osmophore’ is the term used to describe the spongy, pale coloured tissue that generates the foetid odour emitted by the flowers (Fig. 5A–F). In dried specimens, they usually appear as a distinctly paler region (e.g., Fig. 5D), though are sometimes distinctly blackened by poor drying conditions. There are two types of osmophore: the apical osmophore, positioned apical to the tepal margin (Fig. 5B), and the recessed osmophore, positioned within the tepal cavity (Fig. 5A). We found that the shape and positioning of these osmophores on the tepals is highly consistent within species and therefore has taxonomic value (Fig. 5G–P). Species in subgenus *Hydnora* possess just one, large recessed osmophore (Fig. 5G), while those in subgenus *Dorhyna* have both an apical and recessed osmophore (Fig. 5K–N), though the former is most conspicuous. Subgenus *Neohydnora* (*H. esculenta*) only has an apical osmophore (Fig. 5J), while subgenus *Sineseta* (*H. sinandevu*) only has a recessed osmophore (Fig. 5P).

**Fig. 5.**
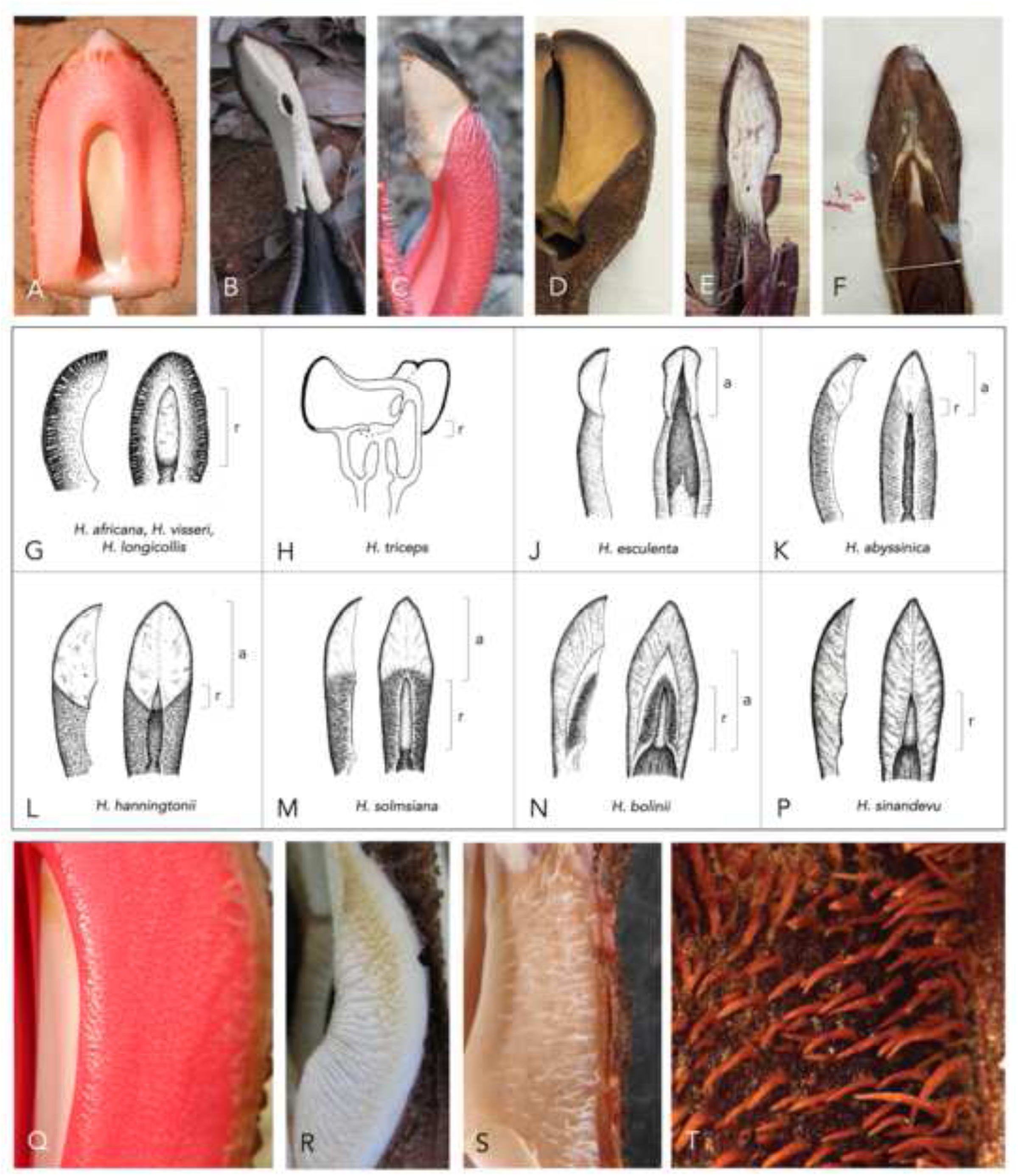
Tepal morphology of *Hydnora*. **A** *Hydnora africana* tepal showing recessed osmophore; **B** *H. esculenta* tepal showing apical osmophore; **C** *H. abyssinica* tepal showing apical osmophore; **D** *H. hanningtonii* dried tepal showing apical osmophore (*Thesiger* s.n. [BM]); **E** *H. solmsiana* dried tepal showing apical osmophore (*Bolin* 09-06 [WIND]); **F** *H. bolinii* dried tepal showing apical and recessed osmophore (*Gillett* 12781 [K]); **G – P** osmophore positioning for all species; **Q** *H. visseri* tepal setae; **R** *H. esculenta* tepal setae; **S** *H. solmsiana* tepal setae; **T** *H. hanningtonii* tepal setae. **G – P** DRAWN BY SEBASTIAN HATT.

### Setae

Setae coverage on the tepal margin does have some taxonomic value despite also being a highly variable trait with size, density and colour changing considerably over the lifespan of the flower (Fig. 5Q–T). *H. sinandevu* is unusual in being entirely glabrous. *H. hanningtonii* has dense, strigose setae that cover the entire tepal margin (Fig. 5T) while *H. abyssinica* and *H. solmsiana* have setae of variable length and density that rarely reach the ventral edge (Fig. 5S). Similarly, *H. visseri* has distinctly shorter and more sparsely distributed setae (Fig. 5Q) than *H. africana*. Setae can be dimorphic in some species, *H. africana*, *H. visseri* and *H. longicollis*, with an additional type of thicker, longer seta found only on the tepal edge. *H. esculenta* is unique in having fleshy ribs in place of setae (Fig. 5R).

### Live material

Tepal colour is not preserved in dried specimens, and is quite variable in living material (Fig. 4H). However, *H. abyssinica* is consistently white inside the floral tube, while *H. hanningtonii* is consistently bright red. Different species are reported to vary in the smell emitted by the flower and the taste of the fruit. While these characters may be useful for field identification and should be recorded in label data, they are variable and subjective and cannot be used in isolation.

Beyond these characters, little else has been found to be of taxonomic value in dried or living specimens. The dimensions of the flower appear to be highly variable, as well as the relative distance between the components of the reproductive chamber and the tepals.

## DISTRIBUTION

*Hydnora* is widespread across southern and eastern Africa, southern Madagascar and the southern Arabian Peninsula (Map 1). The genus has a preference for arid to semi-arid climates, and has been reported from Sudan, Eritrea, Somalia, Ethiopia, Rwanda, Uganda, Kenya, Tanzania, the Democratic Republic of the Congo, Mozambique, Zimbabwe, Angola, Namibia, Botswana, Eswatini and South Africa. Recently, a population resembling *H. abyssinica* was reported from the semi-arid Jos plateau in Nigeria, ∼1,500 km from the nearest known population recorded hitherto (Agyeno et al., 2018). The broad distribution of *Hydnora*, particularly that of the most widespread species *H. abyssinica* and *H. solmsiana*, indicate the tolerance of a range of climatic conditions. For example, *H. esculenta* has been recorded growing in both sub-arid scrubland and transitional rainforest areas (Bolin et al., 2013). Distribution may be more restricted by host range. For example, *H. triceps* has a relatively narrow range in South Africa that follows the narrow range of its host, *Euphorbia dregeana* E.Mey. ex Boiss. In contrast, *H. abyssinica* has a vast range and it can parasitise a number of different Fabaceae hosts with broad ranges across Africa. In addition, it has also been shown to be able to parasitise introduced species such as *Pithecellobium dulce* (Roxb.) Benth. and *Tamarindus indica* L. that may, in theory, have triggered a range expansion. However, very little is known about the climatic requirements of *Hydnora* and, certainly in the case of *H. abyssinica*, many of its hosts can grow in regions that *Hydnora* clearly cannot. The recalcitrance of *Hydnora* to cultivation ex-situ may also suggest that it does have specific requirements, perhaps edaphic as well as climatic, that are not yet appreciated (Thorogood et al., 2022).

The range of *Hydnora* may be underestimated. This is plausible given the dearth in collection in many of these regions, hindered also by a hypogeous habit and infrequent flowering (Sosef et al., 2017). For example, the distribution of *H. sinandevu* was previously thought to be restricted to Kenya and Tanzania. However, our research has revealed collections of *H. sinandevu* in Saudi Arabia, Somalia and Ethiopia.

## HOST SPECIFICITY

Our understanding of host specificity is poor for holoparasitic plants, because collectors rarely excavate to confirm the point of host attachment. For this reason, many host records are unreliable. For example, *H. solmsiana* was observed by the author growing many metres from its verified host; despite flowering next to a different species of potential host.

Host range across the genus is summarised in Table 1. Species in the southern African subgenus *Hydnora* appear to be restricted to succulent *Euphorbia* (Euphorbiaceae) hosts. Interestingly, the four *Hydnora* species in this subgenus have mutually exclusive hosts, even in sympatry (Bolin et al., 2011). While *H. visseri*, *H. longicollis* and *H. triceps* appear very restricted to either one, two or three host species, *H. africana* has been recorded to parasitise at least 12 species. These 12 known hosts of *H. africana* are found in different subgenera and sections of *Euphorbia*, rather than being closely related as might be expected given the level of specificity seen in *H. visseri*, *H. longicollis* and *H. triceps.* This raises the question of why these specific hosts are preferred, but not other more closely related *Euphorbia* species native to the same locality. Understanding this pattern of host preference in subgenus *Hydnora* requires knowledge of the mechanisms of host preference and rejection in these species that we currently lack. Notably, in the only germination study of *Hydnora*, *H. triceps* only germinated in response to cut root exudates of its host and not for other *Euphorbia* species (Bolin et al., 2009c).

Species of subgenus *Dorhyna* parasitise trees in the Fabaceae. The widespread *H. abyssinica* has 16 recorded hosts, *H. solmsiana* has 14 recorded hosts and *H. hanningtonii* has three recorded hosts, with most of these belonging to *Vachellia* and *Senegalia* (both ex *Acacia*). In subgenus *Neohydnora*, *H. esculenta* parasites at least five species of Fabaceae from four different genera. *H. sinandevu* is unusual in parasitising exclusively species of *Commiphora* (Burseraceae).

According to the most recent phylogeny for the genus (Jost et al., 2022), a shift has occurred from the parasitism of CAM-photosynthesising *Euphorbia* (Malpighiales) to C3-photosynthesising Fabaceae (Fabales). Whether there is any significance in the difference in photosynthetic metabolism between these two groups is unknown. The identity of the ancestral host family remains uncertain, although the phylogeny suggests that the ancestral *Hydnora* parasitised Fabaceae (Bolin, 2009b; Jost et al., 2022). Therefore, a transition must presumably also have been made from Fabaceae to *Commiphora* (Burseraceae) in *H. sinandevu*. The fact that these family-level transitions appear to correlate with genetically defined lineages of *Hydnora* may suggest that host specificity has contributed to genetic divergence in the genus.

Interestingly, some *Hydnora* appear to be able to parasitise certain non-native hosts that have only been introduced to the native region in the last few centuries. There are at least two incidences of *H. abyssinica* parasitising introduced Fabaceae hosts, namely *Albizia lebbeck* (L.) Benth., *Delonix Regia* (Bojer ex Hook.) Raf. and *Tamarindus indica* L. (Bosser, 1994; Agyeno et al., 2018). Furthermore, both *H. hanningtonii* and *H. esculenta* have been recorded on the non-native *Pithecellobium dulce* (Roxb.) Benth. Indeed, Bolin et al. (2013) report that this introduced species was the most common host encountered for *H. esculenta* in Madagascar.

## INFRAGENERIC CLASSIFICATION

We follow the infrageneric sections of Decaisne (1873), added to by Harms (1935), but we make modifications to accommodate new taxa, in line with the existing phylogenetic data for the genus (Bolin et al., 2009b; Jost et al., 2022). The revised subgeneric classification is presented summarised in Table 2. Subgenus *Euhydnora* is renamed here to subgenus *Hydnora*; an autonym generated in line with nomenclatural code requirements that the subdivision of a genus that includes the type of the generic name must repeat the generic name unaltered as its subdivisional epithet (Art. 22.1, 22.2, Turland et al., 2018). *H. triceps* was previously positioned in its own subgenus, subg. *Tricephalohydnum* by Harms (1935) due to its uniquely fused tepals. Subgenus *Tricephalohydnum* is included here in subgenus *Hydnora* due to its shared characters of recessed osmophores and angular rhizomes with tubercles in distinct parallel lines, as previously suggested by Musselman & Visser (1989).

**Table 2.**
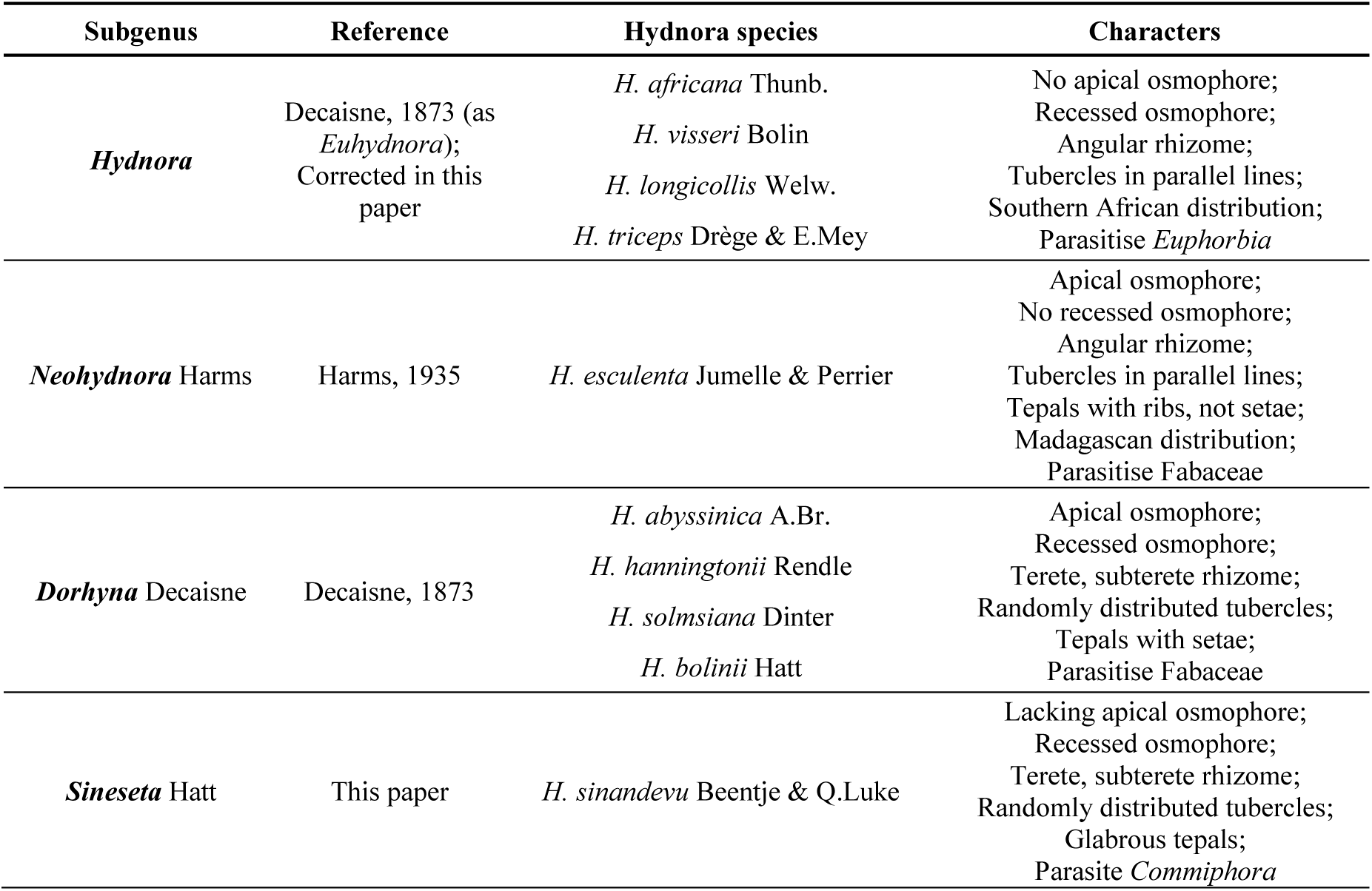
Revised subgeneric classification for Hydnora. A summary of characters is provided for each subgenus

Subgenus *Dorhyna* previously contained *H. abyssinica* and *H. hanningtonii* (as *H. arabica*). Here *H. solmsiana* and *H. bolinii* are assigned to subgenus *Dorhyna* based on their shared rhizome morphology, Fabaceae hosts and positioning of the osmophores. Subgenus *Neohydnora* remains in place and contains only *H. esculenta*. On the basis that *H. sinandevu* lacks both setae and an apical osmophore, and parasitises *Commiphora* (Burseraceae) rather than Fabaceae, thus conflicting considerably with the criteria for subgenus *Dorhyna*, it is moved from subgenus *Dorhyna* to its own new section, subgenus *Sineseta* (translating to ‘without setae’ in Latin). A widely sampled molecular phylogeny will be required to confirm these taxonomic decisions, although they already comply with existing phylogenetic trees for the genus (Bolin et al., 2009b; Jost et al., 2022).

## SYSTEMATIC TREATMENT

### Hydnora

*Thun.* (Thunberg 1775: 69). Type: *Hydnora africana* Thunberg (1775: 69).

*Aphyteia* Acharius & von Linné (1776: 7). Type: *Aphyteia hydnora* Acharius & von Linné (1776: 10).

Herbaceous subterranean perennial root holoparasite, without leaves or stem. *Rhizome* terete, subterete or angular; rhizome surface coriaceous, dark brown, lighter coloured (when fresh) near growth tip; rhizome spreads laterally and often bi- or trifurcating or branching irregularly to form smaller terminal branches; rhizome ornamented with numerous lateral tubercles of variable size and shape, randomly distributed or arranged in parallel lines on the surface; tubercles remain same or develop into flower or develop into haustorium; numerous flowers and flower buds on single rhizomes; rhizome fleshy, brick red to reddish pink internal tissue when broken. *Flower* 3 – 5 merous; flower emerges only partially from soil or not at all; two floral chambers, androecial chamber subtended by gynoecial chamber, inner surfaces of chambers glabrous, pedicel absent or present up to 2.5 cm long. *Perianth* external tissues coriaceous, dark brown to reddish brown; perianth tissues fleshy; perianth tube red, pink or white inside, darkening to red-brown or black when dried; tepal margins present with setae present or absent; tepal margins red to white to orange, lanceolate to elliptic, fused or not fused. *Osmophore* positioned either on tepal apices or recessed in tepal cavity or both; spongy, cream to white (in life), darkening to tan or black when dried, generating foetid odour. *Androecium* antheral ring formed by connate inverted V-shaped anther lobes, forming a central orifice, transversely striate and divided into numerous horizontal pollen sacs. *Gynoecium* ovary inferior, unilocular with numerous ovules produced from apical placenta; lobed and cushion-like stigma on the floor of the gynoecial chamber. *Fruit* subterranean, globose, leathery brown pericarp. *Seed* spherical, black-brown, thousands of seeds embedded within mealy white pulp.

### Distribution

Ten species distributed across the arid and semi-arid regions of southern and eastern Africa, southern Madagascar, Nigeria and the southern Arabian Peninsula. Map 1.

## KEY TO SPECIES OF *HYDNORA*

1. Rhizome angular, usually 4–7 angled, with tubercles in distinct parallel lines 2
2. Rhizome terete, subterete or laterally compressed, with tubercles randomly distributed 6
3. Osmophores only apical, not recessed; tepal margins white, ribbed, without setae; flowers often 5–merous, (Madagascar and Mascarene Islands) 5. **H. esculenta**
4. Osmophores not apical, only recessed; tepal margins red, orange or pink, not ribbed, with setae; flowers usually 3–merous, (mainland Southern Africa) 3
5. Tepals entirely fused for apical half of length thus forming 3 isolated funnel-shaped apertures; flower entirely hypogeous; parasitising *Euphorbia dregeana* 4. **H. triceps**
6. Tepals not fused at apex or only transiently; flower not or only partially hypogeous, parasitising *Euphorbia* spp. other than *E. dregeana* 4
7. Tepals 0.9 – 2.4 cm long; tepal margins often orange; less than 1/5 flower above ground; parasitising *Euphorbia damarana* (Angola, northern Namibia) 3. **H. longicollis**
8. Tepals 2.5 – 9 cm long; tepal margins often pink to red; more than 1/5 flower above ground; parasitising *Euphorbia* spp. other than *E. damarana* (Namibia, South Africa) 5
9. Tepals 2.5 – 6 cm long; tepal margins red to orange; tepals usually lightly fused at the apex at anthesis; parasitising *Euphorbia* spp. other than *E. gregaria* or *E. gummifera* 1. **H. africana**
10. Tepals 5.5 – 9 cm long; tepal margins red to pink; tepals usually free at the apex at anthesis; parasitising *Euphorbia gregaria* or *E. gummifera* **H. visseri**
11. Tepal margins entirely glabrous; apical osmophore absent, recessed osmophore inconspicuous; parasitising *Commiphora* spp. 10. **H. sinandevu**
12. Tepal margins with at least some setae; apical osmophore present, recessed osmophore present but may be inconspicuous; parasitising Fabaceae spp 7
13. Tepal margins and perianth tube entirely red; apical osmophore occupying ½ or more of the total tepal length; setae strigose **H. hanningtonii**
14. Tepal margins red to pink and perianth tube white internally; apical osmophore occupying less than ½ of the total tepal length; setae villosa 8
15. Apical osmophore reduced to narrow inverted V-shaped strip above setae margins, outside this a glabrous outer cucullus; reduced tepal margins with setae not reaching dorsal edge of tepal; recessed osmophore prominent, extending down almost to basal end of tepal margin **H. bolinii**
16. Apical osmophore clearly present, not as above; tepal margins with setae reaching dorsal edge of tepal; recessed osmophore present or inconspicuous 9
17. Recessed osmophore small or inconspicuous; tepal margins usually reddish-pink; border between apical osmophore and tepal margins very distinct, descending sharply from dorsal to ventral edge 6. **H. abyssinica**
18. Recessed osmophore prominent, extending down almost to basal end of tepal margin; tepal margins usually pink to whitish-cream; border between apical osmophore and tepal margins very indistinct, not descending sharply from dorsal to ventral edge, often horizontal or ascending from dorsal to ventral edge 8. **H. solmsiana**

**I) Hydnora** subgenus **Hydnora**. Type: *Hydnora africana* Thunberg

*Hydnora* subgenus *Euhydnora* Decaisne (1873: 75)

*Rhizome* angular (4 – 8 sided) in cross-section, though growth tip and terminal several cm of rhizome may be terete with thinner periderm; rhizome ornamented with numerous lateral tubercles arranged in distinct parallel lines, with these lines defining the number of angles of the rhizome in cross section, on older specimens many of the tubercles may be swollen and distorted such that it is less evident that they are arranged in clear parallel lines. *Flower* usually 3-merous (rarely 4). *Perianth* tube red to pink to orange (in life), then darkening to dark red to brown or black over several days; tepals apically fused or not fused, if not fused then elliptic-lanceolate and curving inwards towards apex; tepal margins with setae though often conspicuous, tepal edges with or without large setae. *Osmophore* recessed in tepal cavity, spongy and white (at anthesis), darkening to grey/brown over several days, generating strong odours of rotting meat. *Gynoecium* with fleshy pedicel absent or rarely present.

**1. Hydnora africana** *Thunberg* (1775: 69); Meyer (1833: 98); Pappe (1862: 51); Thunberg (1823: 499); Solms-Laubach (1901: 6); Dinter (1909: 57); Thistleton-Dyer (1912: 486); Marloth (1913: 177); Dinter (1923: 424); Vaccaneo (1934: 436); Harms in Engler (1935: 288); Schreiber in Merxmüller (1968: 41); Musselman & Visser (1989: 323); Bolin et al. (2009b: 259); Bolin et al. (2018: 107). Type: South Africa, Caput Bonae Spei [Cape of Good Hope] [Western Cape], 1772–1775, *Thunberg* s.n. (lectotype LD! [LD1239903], selected here; isolectotype UPS [UPSV126487).

*Aphyteia hydnora* Acharius & von Linné (1776:10). Type: as for *Hydnora africana*.

*Aphyteia africana* (Thunb.) Oken (1841: 801); Merrill (1950: 270). Type: as for *Hydnora africana*.

Herbaceous subterranean perennial root holoparasite, without leaves or stem. *Rhizome* angular (4 – 8 sided) in cross-section, 1 – 3 cm in diameter, growth tip and first several cm of rhizome terete with thinner periderm; rhizome surface coriaceous, brown to dark brown, lighter coloured near growth tip (in life); rhizome spreads laterally, may occasionally bi- or tri-furcate; rhizome ornamented with numerous lateral tubercles arranged in distinct parallel lines, with these lines defining the number of angles of the rhizome in cross section, tubercles remain same or develop into flower or develop into haustorium, on older specimens many of the tubercles may be swollen and distorted such that it is less evident that they are arranged in clear parallel lines; internally fleshy, coloured bright red to pink. *Flower* usually 3-merous (rarely 4); flower emerges only partially from the soil; two floral chambers, androecial chamber subtended by gynoecial chamber, inner surfaces of chambers glabrous. *Perianth* external tissues coriaceous, brown, total length 8.2 – 19.5 cm; perianth tissues fleshy; perianth tube pink (in life), then darkening to orange and red over several days, 2 – 3.5 cm wide; tepal margins red to orange, tepals (measured from apex to point of connation with adjacent tepal) 2.5 – 6 cm long × 2 – 6.5 cm wide (measured at midpoint); tepals elliptic-lanceolate, curving inwards towards apex, not fused at apex; tepal margin with short setae up to 2 mm, with almost complete coverage across surface, largest on dorsal side but still visible even on ventral side; dorsal tepal edge lined with single row of considerably larger, conspicuous, broad, pale setae up to 6 mm. *Osmophore* recessed in tepal cavity, spongy and white (at anthesis), darkening to grey/brown over several days, generating strong odours of rotting meat. *Androecium* antheral ring formed by connate w-shaped anther lobes, 1.0 – 2.1 cm wide, forming a central orifice, transversely striate and divided into numerous horizontal pollen sacs; pollen bisulcate. *Gynoecium* ovary inferior, unilocular with numerous ovules, ovary 2.2 – 4.1 cm wide; sessile, cushion-like stigma on the floor of the gynoecial chamber, stigma 1.9 – 2.4 cm wide, fleshy pedicel sometimes present, 0 – 5 cm. *Fruit* subterranean, turbinate berry, leathery pericarp, 7 – 18 cm in diameter. *Seed* spherical, black-brown, 0.7 – 1.2 mm, thousands of seeds within fruit, embedded within white pulp. Fig. 6.

**Fig. 6.**
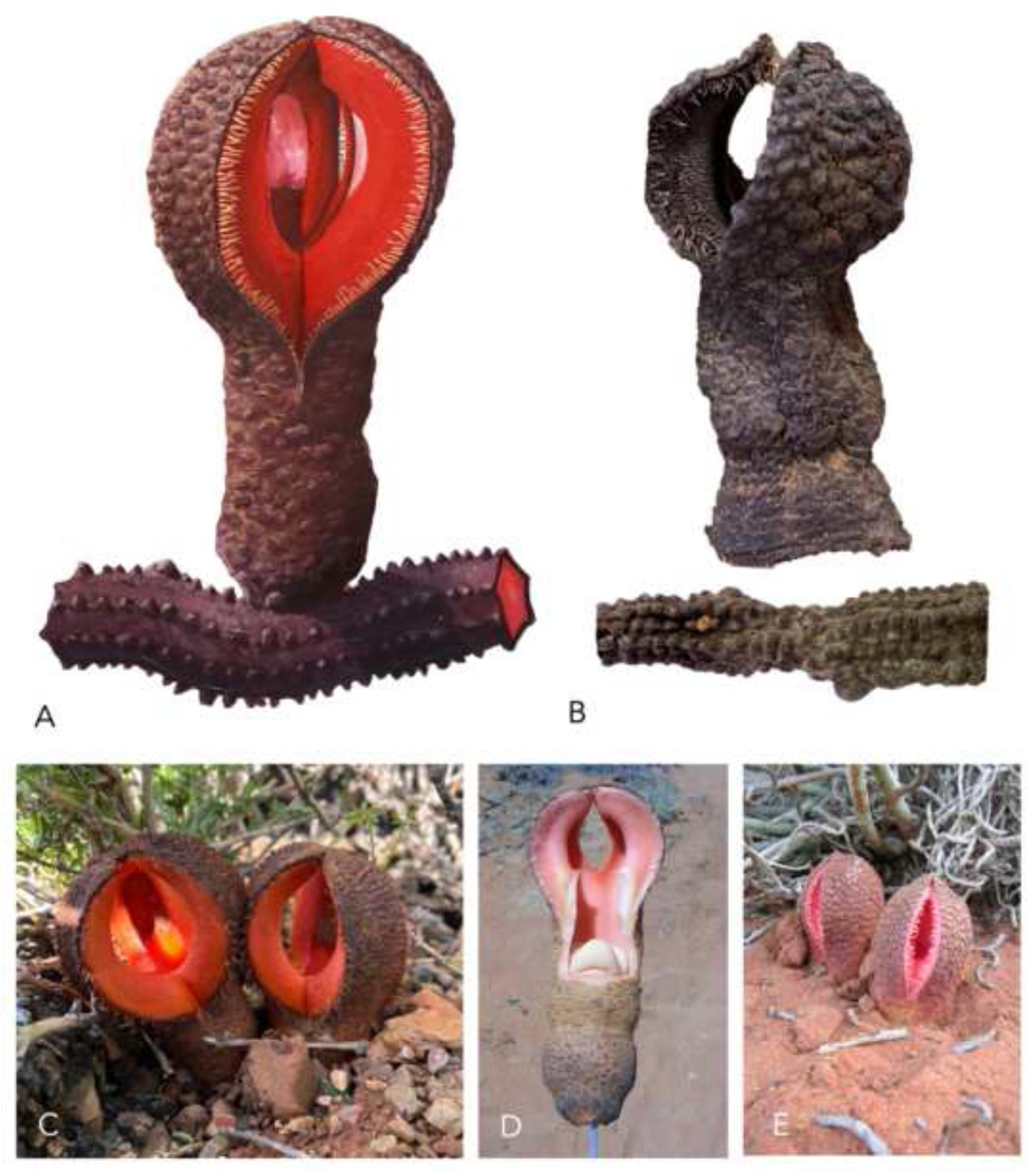
*Hydnora africana*. **A** illustration of flower and rhizome; **B** example of herbarium specimen flower and rhizome (both *Bolin* 09-11 [WIND]); **C–E** live flowers. **A** ILLUSTRATED BY SEBASTIAN HATT.

### DISTRIBUTION

*H. africana* follows the distribution of its succulent *Euphorbia* hosts across southern Africa. It occurs from the Karas Region of southern Namibia (with a single collection in the Erongo Region), down into South Africa through the Northern, Western and Eastern Cape provinces. There are a few records in KwaZulu-Natal, and single records from the North West province and Free State province. It has generally been collected within 200 miles of the coast, with almost no specimens having been found in the interior provinces of South Africa. Within Namibia, it tends to be restricted to inselbergs or areas with winter rainfall (Bolin, 2009). Map 2.

### SPECIMENS EXAMINED

**NAMIBIA** Erongo Region: Omaruru, Brandberg, 15 April 1987, *Craven* 2765 (WIND!); Karas Region: Luderitz Dist., Namib-Naukluft Park [26°40’12.3”S, 15°22’48.8”E], 10 Nov. 2008, *Bolin* 09-24 (WIND!); Luderitz Dist., Spitzkoppe Farm [27°51’46”S, 16°43’04”E], 4 Aug. 2016, *Chase & Rugheimer* FC1071 (WIND!); Tschaunaup [2617DA], 25 Feb. 1947, *Gerstner* 6309 (PRE!); Warmbad, Farm Sperlingspütz [2181CA], 27 May 1972, *Giess & Muller* 12241 (WIND!); Rosh Pinah, Farm Namuskluft [S27 52.127, E16 51.727], 18 Oct. 2005, *Bolin* 08-12 (WIND!); Rosh Pinah, Farm Namuskluft [S27 52.127, E16 51.727], 3 Dec. 2008, *Bolin* 05-3 (WIND!). **SOUTH AFRICA.** Northern Cape: Port Nolloth, Nick Kotze Farm [S29 16.225 E17 6.477], 17 Feb. 2009, *Bolin* 09-11 (WIND!); Port Nolloth [2916BD], 6 June 1951, *Hughes* s.n. (NBG!); Richtersveld, E foothills of Numeesberg [2816BD], 3 Oct. 1972, *Wisura* 2523 (NBG!); Richtersveld, Cornel’s Kop [2816BD], 1 July 1978, *Williamson* 2818 (NBG!); Namaqualand [c2917], May 1916, *Marloth* 7425 (PRE!); Richtersveld, Khubusi District, SW side below Kornelskop on Bloeddrif Road [28°25.237’S, 16°35.047’E], 11 Sept. 2002, *Venter & Venter* 9804 (MO!, NY!, S); Richtersveld 11 [28°22’29”S, 16°54’52”E], 10 Sept. 2010, *Bester* 10105 (PRE!); Lekkersing, Karuchabpoort [2917AA], 26 Aug. 1977, *Oliver et al.* 144 (PRE!); Pofadder Dist., Great Bushmanland, 15 Nov. 1961, *Horn* 5 (NBG!); Pofadder Dist., Naroep [2818DC], 10 July 2008, *Bruyns* 11226 (BOL!, NBG!); Little Namaqualand, Anenous, 1931, *Orpen* s.n. (BOL!); Namaqualand, Veldevreden?, Oct. 1939, *Hanekam?* s.n. (SAM!); Kleinsee, 1931, *Orpen* s.n. (BOL!); Holgat, 1926, *Pillans* s.n. (BOL!); Between Zilverfontein, Kooperberg and Kaus, no date, *Drege* s.n. (K!, HBG!, L); Western Cape: Klaver, Varyhnsdorp [3118DC], 28 Sep. 1959, *Drumsden* s.n. (NBG!); Belville Distr., N of Cape Town [3318CB], 3 Jan. 2001, *Burgoyne & Lewis* 3 (PRE!); Malmesbury, Langebaan, 10 May 1944, *Chaplin* 4 (NBG!); Calvinia [3119BD], July 1937, *Schmidt* 602 (PRE!, K!); Malmesbury, Hopefield [3318AB], no date, *Marloth* 10559 (PRE!); Saldana, no date, *Osbeck* s.n. (S); Rooifontein Farm, 6 km N of Karoopoort, [3319BA], 4 Oct. 1986, *Hilton-Taylor* 1666 (NBG!); Near Ceres Karoo and Witzinberg, 20 Oct. 1973, *Carlquist* 4840 (HUH!, US!); Calvinia, Touws-Rivier, 1 Dec. 1986, *Breckle* 10179 (M!); Calvinia, Lokenburg [3119CA], 12 Dec. 1953, *Acocks* 17363 (PRE!); Worcester, Brewelskloof [3319CD], 15 Oct. 1978, *Bayer* 1564 (NBG!); Worcester, in veld opposite entrance to “Kanetvlei” [3319DA], 16 Sep. 1980, *Walters* 2448 (NBG!); Worcester [3319CB], 19 Nov. 1925, *Pole-Evans* 3023 (PRE!); Worcester [3319CB], Oct. no year, *Marloth* 5701 (PRE!); Worcester, Karoo Botanic Gardens [3319CB], 4 Oct. 1991, *Steiner* 2418 (NBG!); Worcester, Karoo Botanic Gardens, 1946, *Leighton* 2835 (BOL!); Worcester, Karoo Botanic Gardens, 6 April 1985, *Musselman* 7041 (K!); Worcester, Karoo Botanic Gardens, 21 Dec. 1946, *Compton* 18661 (NBG!); Swellendam, 20 miles from Heidelberg Cape, 7 Oct. 1949, *Kramer* s.n. (NBG!); Swellendam, 1930, *Hurling* s.n. (BOL!); Prince Albert, Swartzkraal, 17 July 1939, *Oosthuyzen* s.n. (NBG!); Ladismith, on farm of Mrs RW von Moltke, 16 Oct. 1959, *Von Moltke* s.n. (NBG!); Riversdale, near Riversdale, 22 Feb. 1926, *Muir* s.n. (K!); Riversdale, near Riversdale, 16 Jan. 1928, *Muir* s.n. (K!); Cape Peninsula, Llandudno, 26 June 1940, *Bond* 403 (NBG!); Cape Peninsula, Llandudno, 9 Jan. 1938, *Hafström & Acocks* 409 (S); Cape Peninsula, Houdeklip Bay, between Llandudno and Hout Bay [3418AB], 4 Oct. 1925, *Dumsday* 6653 (PRE!); Cape Town, Llandudno Beach, 1941, *Esterhuysen* 7621 (BOL!); Llandudno, no date, *Acocks* s.n. (S). Eastern Cape: East London [3327BB], Oct. 1964, *Buys* 34 (PRE!); East London, Buffalo River, 2^nd^ Creek [3327BB], 2 March 1898, *Heaton & Galpin* 3189 (PRE!); East London, 2^nd^ Creek [3327BB], Jan. 1927, *Smith* 3852 (PRE!); East London, bank of Nahoon River, 6 Oct. 1900, *Medley-Wood* 53471 (PRE!); East London, 1933, *Everitt* s.n. (BOL!); East London [3327BB], Nov. 1917, *Doidge* s.n. (PRE!); East London, Nov. 1962, *Bokelmann* 3 (NBG!); Kidd’s Beach, 30 miles W of East London [3327BA], 9 Feb. 1967, *Venter* 3273 (NBG!, PRE!); Queenstown Distr., banks of White Kei River, Gwatyu [c3127CC], Jan 1911, *Spence* 8005 (PRE!); King William’s Town, Convent of the Sacred Heart [3227CD], no date, *de Victoria* s.n. (PRE!); Port Alfred, Kowie River Preserve [S33 34.055, E26 51.751], 25 Nov. 2008, *Bolin* 0-82 (WIND!); Kowie River West [3326DB], Aug. 1917, *Tyson* TRV19227 (PRE!); Bathurst Distr., Port Alfred, 23 Aug. 1948, *Symons* s.n. (NBG!); Bathurst Distr., Slankpe Farm, 1959, *collector unknown* s.n. (GRA!); Alexandria, 1946, *Holland* s.n. (BOL!); 15km E of Alexandria, 1978, *Kopke* s.n. (GRA!); Jansenville [3224DC], 4 Jan. 1934, *Long* 1322 (PRE!, K!); Jansenville, Wolverfontein [3224DC], Nov. 1885, *MacOwan* 1724 (PRE!, K!, HUH!, BM!); Steyterville, 1915, *Hepburn* s.n. (GRA!). KwaZulu-Natal: Tugela Ferry and Keats Drift, Aug. 1937, *Smith* 64032 (PRE!); Tugela Valley [2830D], no date, *Smith* s.n. (PRE!). Free State: Close to Rietrivier, before 1887, *MacOwan* 1488 (BM!). North West Prov.: Zwartruggens Distr., Somerset, ad Loot’s Kloof, no date, *Leonard* s.n. (BM!). Unlocalised: no date, *collector unknown* s.n. (MO [MO2740130]); no date, *collector unknown* s.n. (MO! [MO2861368]); early 1800s, *Zeyher* 1511 (K!, G!); 1877, *Barkley* s.n. (K!); no date, *Brown* s.n. (BM!); Kap, early 1800s, *Ecklon & Zeyher* s.n. (B!); Cape of Good Hope, no date, *Masson* s.n. (BM!); no date, *McGibbon* s.n. (K!); Cap. B. Spei., 1772–1775, *Thunberg* s.n. (lectotype LD!, isolectotype UPS); ex Africae aristidis desertis, 1772–1775, *Thunberg* s.n. (UPS); Habitat ad Cap. B. Spei, no date, *collector unknown* s.n. (B!); Kaap Prov, Sep. 1887, *collector unknown* s.n. (L!); no date, *collector unknown* s.n. (BM! [BM000939563]); no date, *collector unknown* s.n. (BM! [BM000939564]); no date, *Atherstone* s.n. (K!); Karroo Desert, before 1867, *Kochmann* s.n. (K!); Cap. Ban. Op.?, no date, *collector unknown* s.n. (M!); no date, *Thunberg* s.n. (S); Goda Hoppsudden, no date, *Thunberg* s.n. (S); Habitat ad Cap. B. Spei, no date, *Thunberg* s.n. (S); no date, *collector unknown* s.n. (S [S11-11398]); no date, *collector unknown* s.n. (S [S-PLE-E10324]); before 1867, *Baron von Ludwig* s.n. (K!); no date, *collector unknown* s.n. (BM! [BM000939567]).

### HABITAT

Rocky or sandy soils amongst succulent Karoo vegetation and coastal forest grasslands, in close proximity to its *Euphorbia* hosts.

### CONSERVATION STATUS

*Hydnora africana* is relatively widespread across South Africa and grows on a number of different *Euphorbia* hosts. In 2008 it was listed as ‘Least Concern, Not Threatened’ in the Red List of South African Plants (Williams et al., 2008). The same ranking is true for all of the recorded hosts, suggesting that *H. africana* is not currently threatened. However, populations may be locally threatened where harvesting for food/medicinal utility is common.

### PHENOLOGY

Flowering time is very dependent of rainfall, which varies considerably across its distribution in South Africa. For example, flowering occurs more around November to January in the summer rainfall areas of the Eastern Cape, while around August to October in the winter rainfall areas of the Northern and Western Cape and around February to April in southern Namibia (Bolin, 2009b). Fruit maturation time is thought to be at least several months to a year.

### ETYMOLOGY

The specific epithet refers to its distribution in South Africa.

### VERNACULAR NAMES

In Afrikaans, Hydnora is commonly known as *jakkalskos*, meaning ‘jackal food’, referring to the tendency of jackals to dig up and eat the fruits. It is also less commonly known as *bobbejaankos*, meaning ‘baboon food’, *Baviaanskos*, *Baviaanskost*, *Jakhalskost*, *Kannip*, *Kannikan* and *Kaw-imp* (Williams et al., 2008; De Beer & Van Wyk, 2011). It is known as *uMavumbaka* in Zulu, meaning ‘the one that pops up’, a name that is shared with *Sarcophyte sanguinea* (Williams et al., 2010). It is also known as *Idololenkonyane*, *Umafumbuka* and *Ubuklunga* in Xhosa (Williams et al., 2008; Bisi-Johnson et al., 2010).

### ETHNOBOTANY

Xhosa men and women are known to use the roots and fruiting body in skin care (Mwinga et al., 2019). They are crushed and macerated with a coarse stone into thin reddish paste, which is applied to the skin for preventing acne and treating pimples, with daily application until symptoms disappear (Dold & Cocks, 2005). The paste is also used for sun protection and is recorded to be used on a daily basis by many outdoor workers. The apparent effectiveness has been suggested to be linked to the high tannin content in the fruit. It has been recorded being sold at traditional medicine market in South Africa, such as Grahamstown Market, Eastern Cape. The fruit is reported to be delectable (Visser, 1981) and apparently eaten by Nama people (*Gerstner* 6309 [PRE0369198]). In 1934, Vaccaneo recorded that the Khoekhoe people eat the fruit raw or roasted. The rhizomes have been used in the treatment of diarrhoea, swollen glands or inflamed throat (Bisi-Johnson et al., 2010), oral thrush (De Beer & Van Wyk, 2011), dysentery (Olajuyigbe & Afolayan, 2012), cardiovascular diseases (Omoruyi et al., 2012) and fever, asthma, constipation, hypertension and oesophageal cancer (Otang et al., 2012). Baboons and jackals have been recorded eating the fruits (Williams et al., 2008; De Beer & Van Wyk, 2011).

### HOST SPECIFICITY

*H. africana* appears to be restricted to succulent *Euphorbia* hosts (Table 1). The following hosts have been recorded, grouped here into subgenera following Bruyns et al., 2006: ***Euphorbia* subgenus *Chamaesyce***: *E. burmannii* (Klotzsch & Garcke) E. Mey. ex Boiss., *E. chersina* N.E.Br., *E. karroensis* (Boiss.) N.E.Br., *E. rhombifolia* Boiss.; ***E.* subgenus *Rhizanthium***: *E. caput-medusae* L.; *E. lignosa* Marloth.; ***E.* subgenus *Euphorbia***: *E. caerulescens* Haw, *E. ingens* E. Mey. ex Boiss., *E. grandidens* Haw., *Euphorbia tetragona* Haw., *Euphorbia triangularis* Desf. ex A.Berger.; ***E.* subgenus *Esula***: *E. mauritanica* L. Within these species, there are representatives from at least four subgenera of *Euphorbia*. Outside Euphorbiaceae, *Cotyledon orbiulatum* L. was reported as a host by Thunberg in the protologue for *H. africana* (1775). However, the validity of this plant as a host is highly unlikely given no evidence has been found of non-*Euphorbia* hosts since then, despite a considerable number of collections and observations having been made.

### NOTES

*H. africana* is the type for the genus and perhaps the most well-known *Hydnora* species. Although *H. visseri*, *H. solmsiana* and *H. abyssinica* have been recorded from South Africa, *H. africana* is by far the dominant *Hydnora* species in South Africa. For much of its range, particularly in the Western and Eastern Cape provinces, it is the only recorded *Hydnora* species. In a few 19^th^ century journals, there are scattered mentions of a species called *Hydnora capensis* from South Africa (McGibbon, 1868) or *Hydnora capensi* Thunb. by Welwitsch (1869). These names are not validly published and are almost certainly synonymous with *H. africana*, although the origin of this confusion is unclear.

*H. africana* is only confusable with *H. visseri* and *H. longicollis* and, morphologically, it falls in between the two. *H. longicollis* predominantly grows in Angola and northern Namibia rather than South Africa, and is distinctly smaller, with tepals < 2.4 cm long and more sharply curved. *H. visseri* also has a more northerly range, although there is some overlap with *H. africana* around the South African-Namibian border region. They are distinguishable from herbarium specimens, with *H. visseri* being distinctly larger with flowers up to 24 cm long and tepals 5.5 – 9 cm long that are more erect than *H. africana*. *H. africana* parasitises a range of species but has not been recorded on the hosts of *H. visseri* or *H. longicollis*, although this should not be treated as a reliable character on its own.

**Hydnora visseri** *Bolin, Maass & Musselman* (2011: 255); Bolin et al. (2018: 107). Type: Namibia, Karas Region: Rosh Pinah, Farm Namuskluft, parasitising *Euphorbia gummifera*, 554 m, 27°54.283 S, 16°50.308 E, 12 Dec 2005, *Bolin* 05-4 (holotype WIND!; isotype US!).

Herbaceous subterranean perennial root holoparasite, without leaves or stem. *Rhizome* angular (4–6 sided) in cross-section, 1–7 cm in diameter, growth tip and first several cm of rhizome terete and with thinner periderm; rhizome surface coriaceous, brown to brown-orange, lighter coloured near growth tip (in life); rhizome spreads laterally, usually at shallow depths less than 30 cm below the soil surface, may occasionally bi- or trifurcate; rhizome ornamented with numerous lateral tubercles arranged in distinct parallel lines, with these lines defining the number of angles of the rhizome in cross-section, tubercles remain same or develop into flower or develop into haustorium; rhizome fleshy, internally coloured dark red (thicker portions) to pale pink (near growth tip). *Flower* usually 3-merous (rarely 2, 4 or 5); flower emerges only partially from the soil; two floral chambers, androecial chamber subtended by gynoecial chamber, inner surfaces of chambers glabrous. *Perianth* external tissues coriaceous, brown, often resembling the soil, total length 10.5–24 cm; perianth tissues fleshy; perianth tube pink (in life), then darkening to orange and red over several days, 4.8–11.2 cm wide; tepals reddish pink to red, tepal (measured from apex to point of connation with adjacent tepal) 5.5–9 cm long × 2.0–5.7 cm wide (measured at midpoint); tepals elongate-lanceolate, curving inwards towards apex, not fused at apex; tepal margin with small setae up to 4 mm, but generally smaller than this and inconspicuous; dorsal tepal edge lined with single row of considerably larger, conspicuous, broad, pale setae up to 6 mm, but generally smaller than this. *Osmophore* recessed in tepal cavity, spongy and white (at anthesis), darkening to grey/brown over several days, generating odours of rotting meat. *Androecium* antheral ring formed by connate w-shaped anther lobes, forming a central orifice, transversely striate and divided into numerous horizontal pollen sacs; pollen bisulcate, 22–27 × 15–20 µm. *Gynoecium* ovary inferior, unilocular with numerous ovules, ovary 2.2–4.5 cm wide; sessile, cushion-like stigma on the floor of the gynoecial chamber, stigma 1–3.1 cm wide. *Fruit* subterranean, turbinate to globose berry, leathery pericarp, 5–20 cm in diameter. *Seed* spherical, black-brown, 0.7–1.2 mm, thousands of seeds within fruit, embedded within white pulp, seed with hard testa and undifferentiated spherical embryo embedded in endosperm. Fig. 7.

**Fig. 7.**
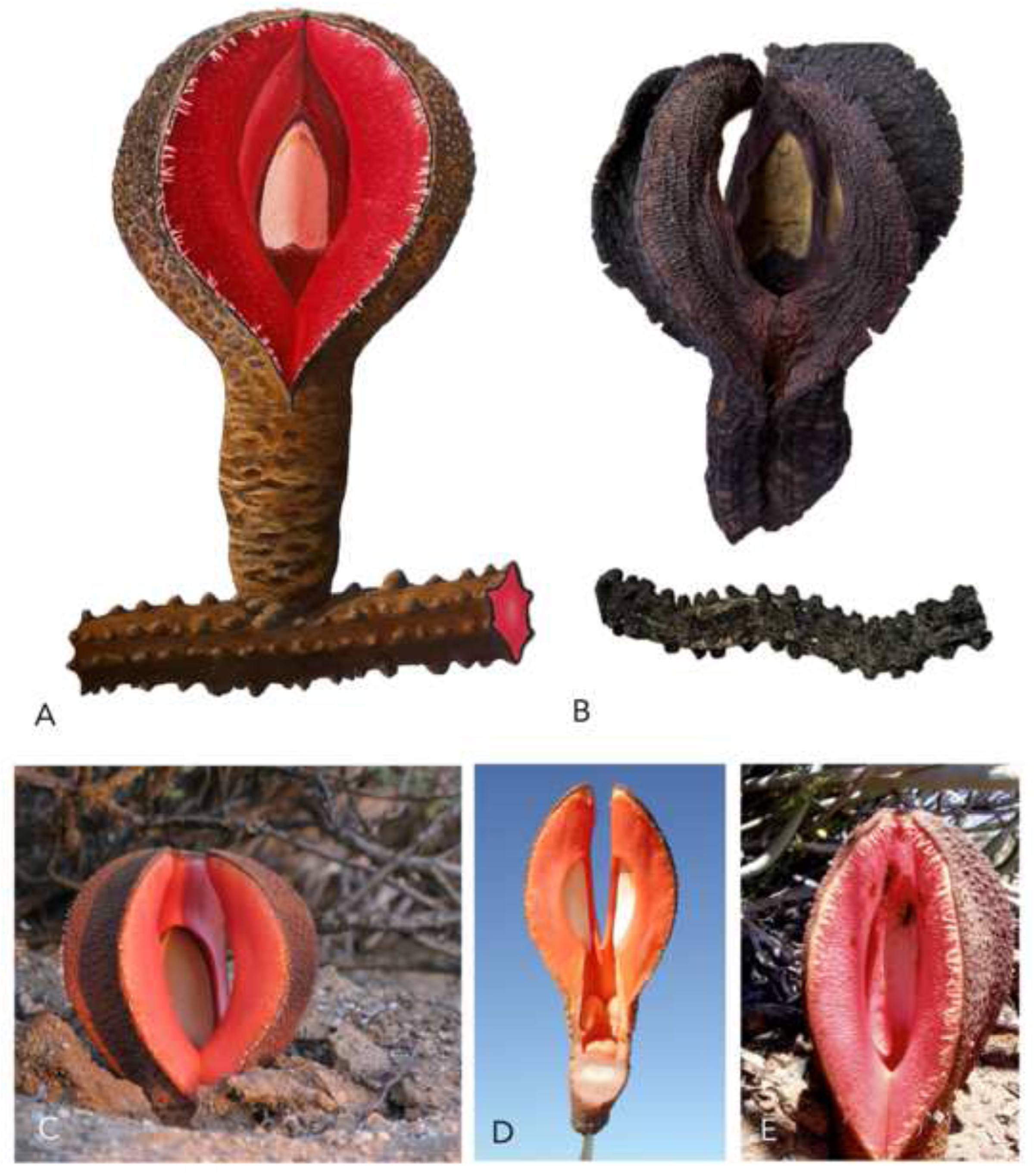
*Hydnora visseri*. **A** illustration of flower and rhizome; **B** example of herbarium specimen flower (*Bolin* 09-24 [WIND]) and rhizome (*Mannheimer* 250 [WIND]); **C–E** live flowers. **A** ILLUSTRATED BY SEBASTIAN HATT.

### DISTRIBUTION

Known primarily from the Hardap and Karas regions of southern Namibia, but also areas of the Northern Cape of South Africa close to the border with Namibia, in proximity to the Orange River. These are all areas of mostly summer rainfall. The distribution appears to match closely with those of its two hosts, *Euphorbia gregaria* and *E. gummifera*. Although not recorded from the eastern portion of the Northern Cape, *H. visseri* is thought likely to grow there given the abundance of *E. gregaria* present (Bolin et al., 2011). Collected from an elevation of 0–1600 m. Map 3.

### SPECIMENS EXAMINED

**NAMIBIA.** Hardap Region: 50 km S of Maltahöhe, 1981, *Lavranos & Pehlemann* 20205 (WIND!); Karas Region: Bethanie Distr., near Kuibis, May 1913, *Range* s.n. (SAM!); South Luderitz, Aug. 1971, *Logan & Jensen* 4966 (WIND!, only rhizomes but likely *H. visseri* due to locality); Farm Altdorn 3 [27°54’S, 17°41’E], 19 Jan. 1987, *Ward* 10117 (WIND!, PRE!, K!); Sandveld, on koppie next to road [27°1’1”S, 18°14’9”E], 27 March 1998, *Strohbach & Dauth* 3755 (WIND!); Sperrgebiet [26°59’47”S, 15°21’58”E], 11 July 1995, *Strohbach* 766 (WIND!); Sperrgebiet, Roter Kamm. [27°37’30”S, 16°22’30”E], 29 Sep. 1996, *Mannheimer & Mannheimer* 425 (WIND!); Eastern Sperrgebiet [27°0’27”S, 15°47’23”E], 25 Sep. 1996, *Mannheimer & Mannheimer* 303 (WIND!); Eastern Sperrgebiet [27°1’13”S, 15°42’58”E], 25 Sep. 1996, *Mannheimer & Mannheimer* 299 (WIND!); Sperrgebiet, approach to Klinghardt Mountains [27°21’2”S, 15°41’11”E], 21 Sep. 1996, *Mannheimer* 250 (WIND!); Karasburg, Farm Karios 8 [27°37’30”S, 17°52’30”E], 15 April 2000, *Strassen* 114 (WIND!); Seeheim, 1927, *Pellans* 5850 (BOL!); Seeheim, Farm Kanas, 20 Oct. 2005, *Bolin* 0-51 (WIND!); Karas, Gondwana Canyon Park [S27 32.850, E17 52.972], 14 Feb. 2009, *Bolin* 908 (WIND!); Farm Jerusalem 73 [28°23’46”S, 19°36’51”E], 7 March 2008, *Klaassen & Rugheimer* EK1823 (WIND!, US); Farm Sandfontein 131 [28°45’36”S, 18°25’18”E], 11 March 2008, *Klaassen & Rugheimer* EK1891 (WIND!); Rosh Pinah, Farm Namuskluft [S27 54.283, E16 50.308], 3 Dec. 2008, *Bolin* 0-85 (WIND!); Rosh Pinah, Farm Namuskluft [S27 54.283, E16 50.308], 12 Dec. 2005, *Bolin* 0-54 (Holotype WIND!, Isotype US!); Unlocalised: 28 Jan. 1929, *Dinter* 6011 (B!). **SOUTH AFRICA.** Northern Cape: Sendlingsdrift [S28 20.699, E16 55.522], 16 Feb. 2009, *Bolin* 9-10 (WIND!); Aggeneys Black Mountain Mine [2918BA], 2 Feb. 2000, *Burgoyne* 7981 (PRE!).

### HABITAT

Flat, sandy soils amongst Nama-Karoo and succulent Karoo vegetation types, in areas where *Euphorbia gregaria* and *E. gummifera* are prevalent. Occasionally seen in rocky soil types.

### CONSERVATION STATUS

Authors of the type description (Bolin et al., 2011) state that *H. visseri* should be considered Least Concern (LC) according to IUCN criteria. It is relatively widespread and common across its range. Furthermore, neither of its hosts are listed as threatened on either the Namibian or South African red lists (Loots, 2005; Williams et al., 2008). Both host species receive some degree of protection as succulent *Euphorbia* under CITES Appendix 2. Although intensive land use for grazing occurs in southern Namibia, the host plants are not palatable food for grazers, and thus their abundance appears to be largely unaffected (Bolin et al., 2011). Indeed, as they have been observed persisting on farms in Namibia it is unlikely that *H. visseri* is currently threatened under the existing regime of land use. More local threats such as mining activity near Rosh Pinah have been noted but are unlikely to be impactful unless there is a significant increase in activity (Bolin et al., 2011).

### PHENOLOGY

Flowers primarily from October to January, although sporadic flowering has been observed all year round, likely due to flowering being dependent on adequate rainfall. Fruit maturation time is thought to be greater than eight months, with ripe fruits often existing alongside flowers from the next year (Bolin et al., 2011).

### ETYMOLOGY

The specific epithet refers to Professor Johann H. Visser (1931–1990), a South African botanist and expert on South African parasitic plants.

### VERNACULAR NAMES

All *Hydnora* species in South Africa are known as *jakkalskos*, *bobbejaankos* or *stinkblom* in Afrikaans. In Nama-Damara/Khoekhoe it is known as *kani* or *kanip* (Bolin et al., 2011).

### ETHNOBOTANY

It is unknown whether local people in Namibia discriminate between *H. africana* and *H. visseri* (See ethnobotany entry for *H. africana*). Mature fruits are reported to be enjoyed as a raw food source by local peoples and bitter immature fruits are made palatable by preparing them by roasting in coals, with or without animal milk (*Ward* 10117 [PRE0796363], Bolin et al., 2011).

### HOST SPECIFICITY

*Euphorbia gregaria* Marloth. and *E. gummifera* Boiss. are confirmed hosts, both in *E.* subgenus *Euphorbia* sect. *Tirucalli*, and there is a single report of *E. gariepina* Boiss., in *E.* subgenus *Rhizanthium* (Table 1). Even in areas where other known *Euphorbia* hosts of *H. africana* co-occur with these species, *H. visseri* seems to exclusively favour *E. gregaria* and *E. gummifera* (Bolin et al., 2011).

### NOTES

The largest of the species in subgenus *Hydnora*, *H. visseri* is distinguished from the similar *H. africana* and *H. longicollis* by its larger, more elongate, erect tepals, and by its unique hosts, *Euphorbia gregaria* and *E. gummifera*. It was only recently separated from *H. africana* as a distinct species (Bolin et al., 2011).

**3. Hydnora longicollis** *Welw.* (Welwitsch 1869: Tab. XXI). Baker & Wright in Thistleton-Dyer (1909: 132); Dinter (1923: 424); Vaccaneo (1934: 439); Harms in Engler (1935: 290); Bolin (2009b: 85); Bolin et al. (2018: 107). Type: Angola, Mossamedes Distr. As far as Cabo Negro, maritime sandy areas, 1859, *Welwitsch* 530 (syntype K!, BM!, G!, LISU!).

*Hydnora africana* var. *longicollis* Welw. (Welwitsch 1869: 66); Hiern et al. (1900: 910); Musselman & Visser (1989: 323). Type: as for *Hydnora longicollis* Welw.

*Hydnora longicollis* Welw. sub var africanae (Welw.) Solms-Laubach in Engler (1901: 6).

Type: as for *Hydnora longicollis* Welw.

Herbaceous subterranean perennial root holoparasite, without leaves or stem. *Rhizome* angular (4-6 sided) in cross-section, 1 – 3 cm in diameter; rhizome surface coriaceous, brown to dark brown; rhizome spreads laterally, usually at shallow depths, may occasionally bi- or trifurcate; rhizome ornamented with numerous lateral tubercles arranged in distinct parallel lines, with these lines defining the number of angles of the rhizome in cross-section, tubercles remain same or develop into flower or develop into haustorium; rhizome fleshy, internally coloured rich red. *Flower* usually 3-merous (rarely 4); flower emerges only partially from the soil; two floral chambers, androecial chamber subtended by gynoecial chamber, inner surfaces of chambers glabrous. *Perianth* external tissues coriaceous, brown, often resembling the soil, total length 5.1 – 14.5 cm long; perianth tissues fleshy; perianth tube pink (in life), then darkening to orange and red over several days, 1.2 – 2.7 cm wide; tepals pinkish, tepal (measured from apex to point of connation with adjacent tepal) 0.9 – 2.4 cm long × 1.2 – 2.7 cm wide (measured at midpoint); tepals elliptic-lanceolate, curving inwards towards apex, not fused at apex; tepal margin with setae up to 2 mm, with almost complete coverage across surface, largest on dorsal side but still visible even on ventral side; dorsal tepal edge lined with single row of considerably larger, conspicuous, broad, pale setae up to 4 mm. *Osmophore* recessed in tepal cavity, spongy and white (at anthesis), darkening to grey/brown over several days, generating foul odour. *Androecium* antheral ring formed by connate anther lobes, 1.0 – 2.1 cm wide, forming a central orifice, transversely striate and divided into numerous horizontal pollen sacs; pollen bisulcate. *Gynoecium* ovary inferior, unilocular with numerous ovules, ovary 1.7 – 3.3 cm wide, sessile, cushion-like stigma on the floor of the gynoecial chamber, stigma 1.0 – 1.8 cm wide. *Fruit* subterranean, turbinate to globose berry, leathery pericarp, 3 – 5 cm in diameter. *Seed* spherical, black-brown, 0.7 – 1.2 mm, thousands of seeds within fruit, embedded within white pulp. Fig. 8.

**Fig. 8.**
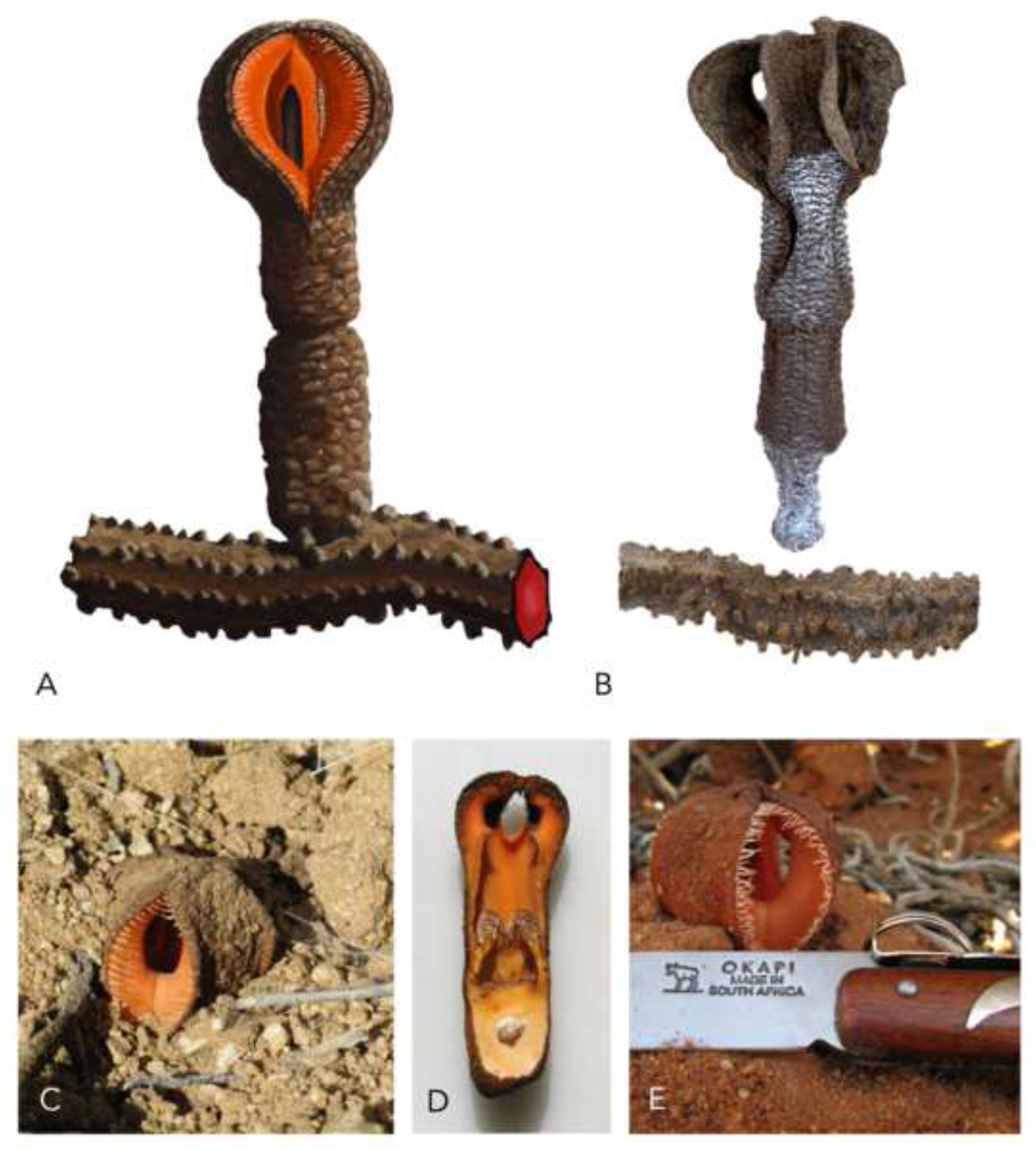
*Hydnora longicollis*. **A** illustration of flower and rhizome; **B** example of herbarium specimen flower (*Carisso & Sousa* s.n. [BM]) and rhizome (*Humbert* s.n. [P]); **C–E** live flowers. **A** ILLUSTRATED BY SEBASTIAN HATT.

### DISTRIBUTION

Known from southwestern Angola and the Erongo region of Namibia. In Namibia, it largely follows the distribution of its host, *Euphorbia damarana*. There are only a few collections in Angola, most of which are in the sandy, coastal Namibe Province. However, a recent collection was made considerably further inland in the Huíla Province. The full extent of its distribution in Angola is likely undiscovered, especially given much of the country has been considerably understudied in the last century due to civil unrest (Bolin, 2009b). Its distribution is largely disjunct from and adjacent to *H. visseri* and *H. africana*. Map 4.

### SPECIMENS EXAMINED

**ANGOLA:** Namibe Prov.: Approx 60 km E of Namibe, 2009, *Voigt* 67 (GHPG!); South of Namibe, 2006, *Bird* s.n. (ODU!); Mossamedes Distr. as far as Cabo Negro, 1859, *Welwitsch* 530 (Syntype K!, BM!, G!, LISU!); Mossamedes, desert SE of Mossamedes, 11 June 1937, *Carrisso & de Sousa* s.n. (BM!); desert SE of Mossamedes, 1937, *Humbert* s.n. (P!). **NAMIBIA.** Erongo Region: 25 km SE of Uis [S21 26.346, E15 03.115], 3 Feb. 2009, *Bolin* 0-92 (WIND!); Omaruru, approx. 15 km S of Omaruru river [21°30’18”S, 15°09’41”E], 10 April 2009, *Bolin* 09-30 (WIND!); Swakopmund, no date, *Weiss* s.n. (M!); Henties Baai, Messum Crater, 2003, *Maass* s.n. (ODU!).

### HABITAT

Sandy, rocky or gravel plains, sometimes in relative proximity to the coast or in dry riverbeds.

### CONSERVATION STATUS

There is very limited data on the conservation status of *H. longicollis*, largely because the full extent of its distribution remains unclear. Much of Namibia’s west coast falls within designated areas with at least some degree of protection. Alternatively, conservation in Angola has largely been neglected in the past, with much of its natural landscape devastated by extensive conflict or unregulated development (Huntley et al., 2019). However, one of its primary hosts, *Euphorbia damarana*, has been listed as ‘not threatened’ according to the national red list, which is encouraging for *H. longicollis* (Ministério do Ambiente, 2018). Considerable further research is clearly required on this matter to make a confident decision on the threat status.

### PHENOLOGY

In Angola, flowering has mainly been observed in January, while in northern Namibia it occurs more in February to April. However, sporadic flowering has been observed all year round, likely due to flowering being dependent on adequate rainfall. Fruit maturation time is thought to be at least several months (Bolin, 2009b)

### ETYMOLOGY

The specific epithet translates to ‘long neck’ from latin. This likely refers to the proportionately long perianth tube seen in some specimens, although this a variable character.

### VERNACULAR NAMES

In Nama-Damara/Khoekhoe, ‘Hydnora’ is known as *kani* or *kanip* (Bolin et al., 2011). It is unclear whether this is specific to *H. visseri* or whether it also applies to *H. longicollis* as well. Vernacular names for *Hydnora* in Angola are currently unrecorded. **ETHNOBOTANY.** Data for the uses of *H. longicollis* are limited to the type description in 1869. Welwitsch (1869) writes that all parts of the plant, particularly the rhizomes, are astringent. At the time, they were sometimes used in the fishing nets and fabric dyes by the Mossamedes locals. He also notes a potential value in healing sores as a mucous membrane. It is unknown whether local people discrimate between *H. longicollis, H. visseri* and *H. africana* in terms of usage, particularly in Namibia where there is overlap in their distributions. One specimen from Munich reports that it belongs to a collection of ‘plants that provide tannins’ (*Weiss* s.n. [M]). Tannins have been reported from other *Hydnora* species (see *H. solmsiana*: Ethnobotany).

### HOST SPECIFICITY

*Euphorbia damarana* L.C.Leach (*E.* subgenus *Euphorbia* sect. *Tirucalli*) is a confirmed host (Table 1). There is a single report on *E. gariepina* and two reports on *E. virosa* Willd. The type description notes *Roepera orbiculata* (as *Zygophyllum orbiculata*), although this was most likely a mistake as it belongs to an entirely unrelated family to *Euphorbia* and no reports have corroborated this since (Welwitsch, 1869) The shortage of herbarium specimens and observational data has considerably limited our understanding of host preference for this species.

### NOTES

Species description adapted from (Bolin, 2009b). *H. longicollis* has suffered much neglect since its discovery in 1869. This is largely due to the political instability in Angola that has made collecting there a challenge, hence the relatively small number of herbarium specimens (Bolin, 2009b). There is some taxonomic confusion surrounding its name, as the protologue cites it as *H. africana var. longicollis* Welw., but the illustration in the same report is labelled as *H. longicollis* Welw (Welwitsch, 1869). Therefore, both names are technically valid according to the International Code of Nomenclature (Turland et al., 2018). Since then, both names seem to have been used interchangeably (Vaccaneo, 1934; Harms, 1935). Bolin et al. (2011) consider *H. longicollis* Welw. to be a species in its own right due to its morphological distinction from *H. africana*, as well as its non-overlapping distribution and distinct host species. Furthermore, an unpublished phylogeny of *Hydnora* from Jay Bolin’s PhD thesis supported this species concept (Bolin, 2009b). Therefore, this species is accepted here.

**4. Hydnora triceps** *Drège & E.Mey.* (1833: 779); Harvey (1863: 187); Solms-Laubach in Engler (1901: 6); Thistleton-Dyer (1912: 487); Marloth (1913: 178); Vaccaneo (1934: 446); Harms in Engler (1935: 293); Musselman & Visser (1989: 323); Bolin et al. (2018: 107). Type: South Africa. Zwischen Zilverfontein, Kooperberg und Kaus, Sep./Oct. before 1833, *Drège* s.n. (syntype K!, HBG!, L!).

*Aphyteia triceps* Steud. (Steudel 1840: 111). Type: as for *Hydnora triceps* Drège & E.Mey.

*Aphyteia multiceps* Burch. (Burchell 1822: 213). Type: as for *Hydnora triceps* Drège & E.Mey.

Herbaceous subterranean perennial root holoparasite, without leaves or stem. *Rhizome* angular, 4–7(–9) sided, in cross-section, 1-3 cm in diameter, rhizome surface coriaceous, dark-brown to almost black; rhizome spreads laterally at various depths between 5–15 cm below ground; rhizome ornamented with numerous lateral tubercles arranged in distinct parallel lines, with these lines defining the number of angles of the rhizome in cross-section, tubercles remain same or develop into flower or develop into haustorium; rhizome fleshy, internally coloured deep scarlet (thicker portions) to pinkish (near growth tip). *Flower* usually 3-merous; flower almost entirely underground, usually mature flowers are up to 5 cm underground although sometimes the top 2 cm of the perianth represented by the 3 opening between tepals will be exposed; two floral chambers, androecial chamber subtended by gynoecial chamber, inner surfaces of chambers glabrous. *Perianth* external tissues coriaceous, pinkish brown, total length 10–15 cm long; perianth tissues fleshy; perianth tube pinkish-yellow, 2–10 cm long and 2.5–3.5 cm wide; tepals broadly clavate but incurved and apically fused to form 3 distinct lateral apertures, each with a pronounced cucullate flesh-coloured mouth; well-developed horizontal flange fused into a prominent circular ridge on the inside below the apertures; tepal interior glabrous. *Osmophore* recessed below each horizontal flange, below the apertures, spongy and white (at anthesis), darkening to dark brown over several days, generating strong foetid odour shortly after opening. *Androecium* 3 erect connate anther lobes, inserted near apex of perianth tube, forming central orifice, transversely striate and divided into numerous vertical pollen sacs with abundant, yellow, slightly sticky pollen. *Gynoecium* ovary inferior, unilocular with numerous pendulous ovules; sessile, cushion-like stigma on floor of the gynoecial chamber, stigma distinctly trilobed. *Fruit* subterranean, turbinate to globose berry, leathery external pericarp, internally bright pink, astringent taste, faint coconut odour, slightly dehiscent at apex when ripe, 3–10 cm in diameter. *Seed* spherical, black-brown, embedded within white pulp. Fig. 9.

**Fig. 9.**
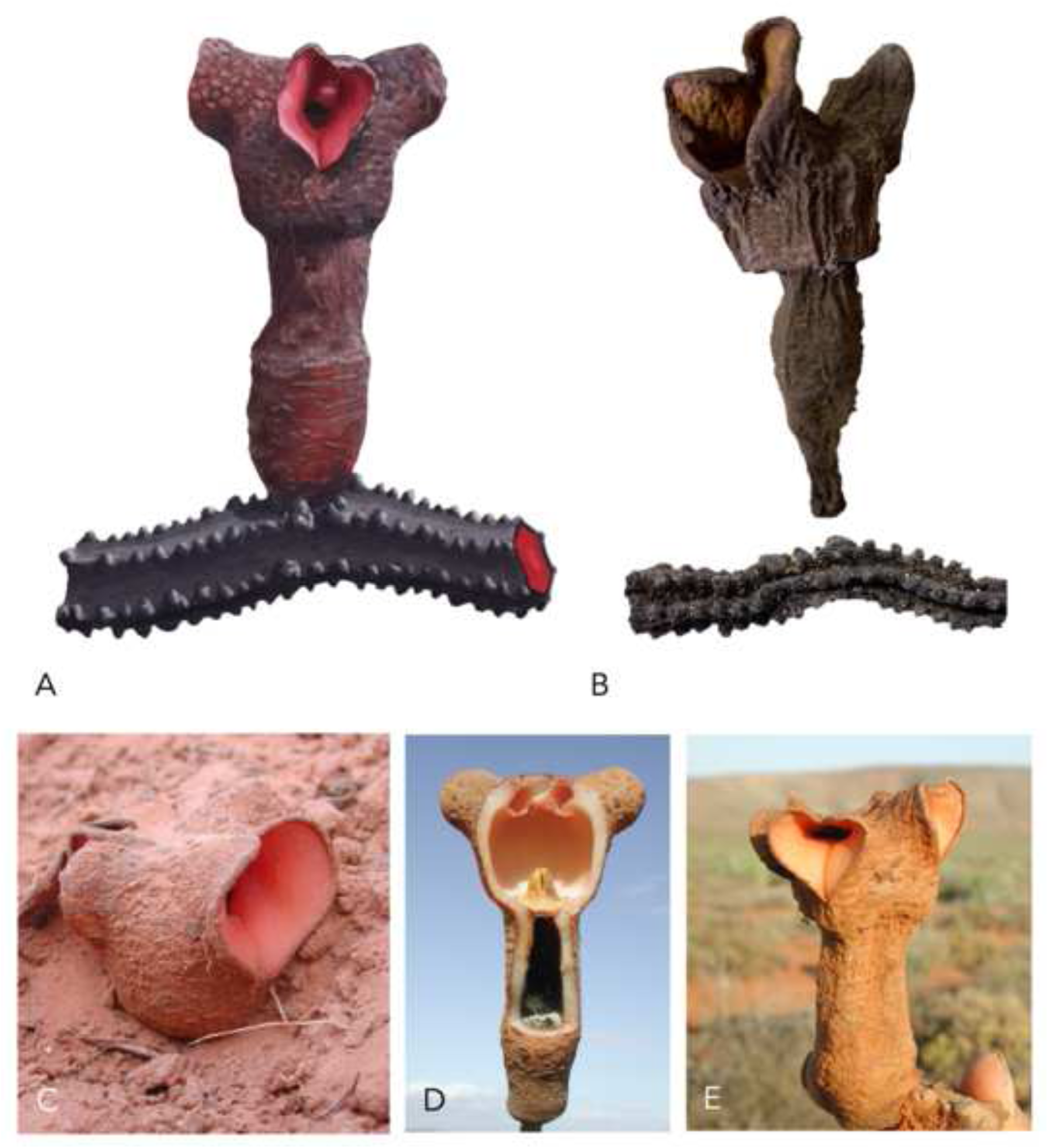
*Hydnora triceps*. **A** illustration of flower and rhizome; **B** example of herbarium specimen flower and rhizome (both *Bolin* 08-7 [WIND]); **C–E** live flowers. **A** ILLUSTRATED BY SEBASTIAN HATT.

### DISTRIBUTION

Known from the Northern Cape of South Africa and the Karas Region of Namibia, following the narrow distribution of *Euphorbia dregeana*. Map 5.

### SPECIMENS EXAMINED

**NAMIBIA.** Karas Region: Rosh Pinah, road verge next to Farm Namuskluft, 14 Sep. 2001, *Maass & Musselman* 1 (WIND!); Rosh Pinah, Farm Namuskluft [S27 56.427, E16 48.141], 3 Dec. 2008, *Bolin* 0-87 (WIND!); Rosh Pinah, Farm Namuskluft [S27 56.427, E16 48.141], 11 July 2009, *Bolin* 09-41 (WIND!); Rosh Pinah, Farm Namuskluft 88 [27°53’27”S, 16°51’21”E], 3 Aug. 2002, *Maass* EM18 (WIND!); Namian’s Saddle [27°53’27”S, 16°51’21”E], 20 Sep. 2003, *Mannheimer* CM2438 (WIND!). **SOUTH AFRICA.** Northern Cape: Port Nolloth, Farm Gemsbokvlei [29°23’47”S, 17°12’40”E], 21 Dec. 2002, *Maass & Musselman* 2 (WIND!); Port Nolloth, Nick Kotze Farm, 17 Feb. 2009, *Bolin* 09-14 (WIND!); Modderfontein, Whitehead, Namaqualand, near O’Kiep, c. 1887, *Hofmeyer* s.n. (K!, SAM!); Zwischen Zilverfontein, Kooperberg und Kaus, Sep./Oct. before 1833, *Drège* s.n. (syntype K!, HBG!, L!); Farm Droëskraal, in the vicinity of Naroegas se Berge, 1988, *Visser* STE31789 (PRE!). **UNLOCALISED.** 1887, *Marloth* 1863 (PRE!); no date, *collector unknown* s.n. (K!, mixed specimen with *H. africana*).

### HABITAT

Sandy plains and stony hills, following the preferred habitats of *Euphorbia dregeana*. Visser (1988) noted a preference in *H. triceps* for sandy habitats.

### CONSERVATION STATUS

Both *Hydnora triceps* and its host *Euphorbia dregeana* appear as ‘Least Concern’ on the Red List of South African Plants (Williams et al., 2008). While there are some potential local threats such as mining activities near Rosh Pinah, the host population seems to be largely stable. While there is intensive land use in the area, populations of both host and parasite appear to be able to survive regardless. This is perhaps because the host plant is not eaten by grazing animals. Indeed, a significant population was found growing on a farm in Namibia in 2004 (Maass & Musselman, 2004).

### PHENOLOGY

Full details of phenology are unknown, but *H. triceps* has been collected in flower during September on two occasions and collected in fruit in December on one occasion. (Visser, 1988; Maass & Musselman 2004).

### ETYMOLOGY

The specific epithet translates to ‘three-headed’, referring to the three distinct apertures formed by the incurved tepals.

### VERNACULAR NAMES

None recorded.

### ETHNOBOTANY

No uses recorded. It is possible that it is used indiscriminately from *H. africana* or *H. visseri*, given the overlap in range. However, given that most specimens don’t even breach the surface of the soil, it is also possible that they go largely unnoticed by local people.

### HOST SPECIFICITY

The only known host is *Euphorbia dregeana* E.Mey. ex Boiss. (in *E.* subgenus *Rhizanthium*) (Table 1).

### NOTES

Species description adapted from (Maass & Musselman, 2004). *H. triceps* is very distinctive and unlikely to be confused with any other species. No other species has fused tepals and an entirely hypogeous habit. It has only been collected a handful of times since its discovery in 1833. It was thought extinct until it was rediscovered in South Africa in 1988 by Johann Visser and later in Namibia in 2001 by Erika Maass (Visser et al., 1988; Maass, 2001). The rarity of this species is unclear. Despite its occurrence in South Africa, a relatively thoroughly surveyed country, the small number of collections may simply reflect that the fact that it is notoriously difficult to see. In most cases, the only evidence of its presence of small cracks in the soil, over a small bulge in the soil, leading to the openings of the flower.

**II) Hydnora** subgenus **Neohydnora** Harms in Engler (1935: 293). Type: *Hydnora esculenta* Jumelle & Perrier

*Rhizome* angular (4 – 7 sided) in cross-section; tubercles arranged in distinct parallel lines, with these lines defining the number of angles of the rhizome in cross-section. *Flower* (3 – (4 – 5) – 6)-merous. *Perianth* external tissues coriaceous, brown to purplish-brown; perianth tube whitish inside, fading to brown-black at the tepals; tepal margins whitish-cream, darkening to brown with age; tepal often strongly recurved towards the apex, not fused at apex; tepal margin with numerous setae, c. 3 mm in diameter, transitioning into a distinctly ribbed surface. *Osmophore* apical, not recessed, often split down the middle such that it is divided into two triangles barely fused at the apex, spongy and white to tan (in life), darkening to yellow when dried, generating musty, foetid odour, but not one of rotting flesh. *Gynoecium* fleshy pedicel sometimes present, 0 – 5 cm long.

**5. Hydnora esculenta** *Jumelle & Perrier* (1912: 327); Vaccaneo (1934: 446); Harms in Engler (1935: 293); Musselman & Visser (1989: 323); Bolin et al., (2013: 1); Bolin et al. (2018: 107). Type: Madagascar, SW Region. Linta and Menandra watersheds, 7 Dec. 1910, *Perrier* 8911 (holotype P!; isotype K!). Epitype (designated by Bolin et al., 2013): Madagascar, Tulear, grounds of hospital, 6 Jan. 1947, *Humbert* 19805 (epitype P!, isoepitypes MO!, G!, K!).

Herbaceous subterranean perennial root holoparasite, without leaves or stem. *Rhizome* angular (4 – 7 sided) in cross-section, 0.7 – 5 cm in diameter; rhizome surface coriaceous, dark brown to orange-brown, lighter coloured near growth tip (in life); rhizome spreads laterally, may occasionally bifurcate; rhizome ornamented with numerous lateral tubercles arranged in distinct parallel lines, with these lines defining the number of angles of the rhizome in cross-section, tubercles remain same or develop into flower or develop into haustorium; rhizome fleshy, internally coloured light pink to red (in life). *Flower* (3 – (4 – 5) – 6)-merous; flower emerges only partially from the soil; two floral chambers, androecial chamber subtended by gynoecial chamber, inner surfaces of chambers glabrous. *Perianth* external tissues coriaceous, brown to purplish-brown, total length 11.6 – 26.9 cm; perianth tissues fleshy; perianth tube whitish inside, fading to brown-black at the tepals; tepals whitish-cream, darkening to brown with age, tepal (measured from apex to point of connation with adjacent tepal) 4.3 – 7.4 cm long × 1.2 – 3.9 cm wide (measured at base); tepals elongate-lanceolate, often strongly recurved towards the apex, not fused at apex; tepal margin with numerous setae, c. 3 mm in diameter, transitioning into a distinctly ribbed surface. *Osmophore* apical, not recessed, often split down the middle such that it is divided into two triangles barely fused at the apex, spongy and white to tan (in life), darkening to yellow when dried, generating musty, foetid odour, but not one of rotting flesh. *Androecium* antheral ring formed by connate w-shaped anther lobes, forming a central orifice, transversely striate and divided into numerous horizontal pollen sacs, pollen bisulcate. *Gynoecium* ovary inferior, unilocular with numerous ovules, ovary 2.1 – 4.6 cm wide; lobed and cushion-like stigma on the floor of the gynoecial chamber, stigma 1 – 3 cm wide, fleshy pedicel sometimes present, 0 – 5 cm long. *Fruit* partially subterranean turbinate berry, 7 – 15 cm in diameter. *Seed* spherical, black-brown, 0.7 – 1.2 mm, thousands of seeds within fruit, embedded within white pulp. Fig. 10.

**Fig. 10.**
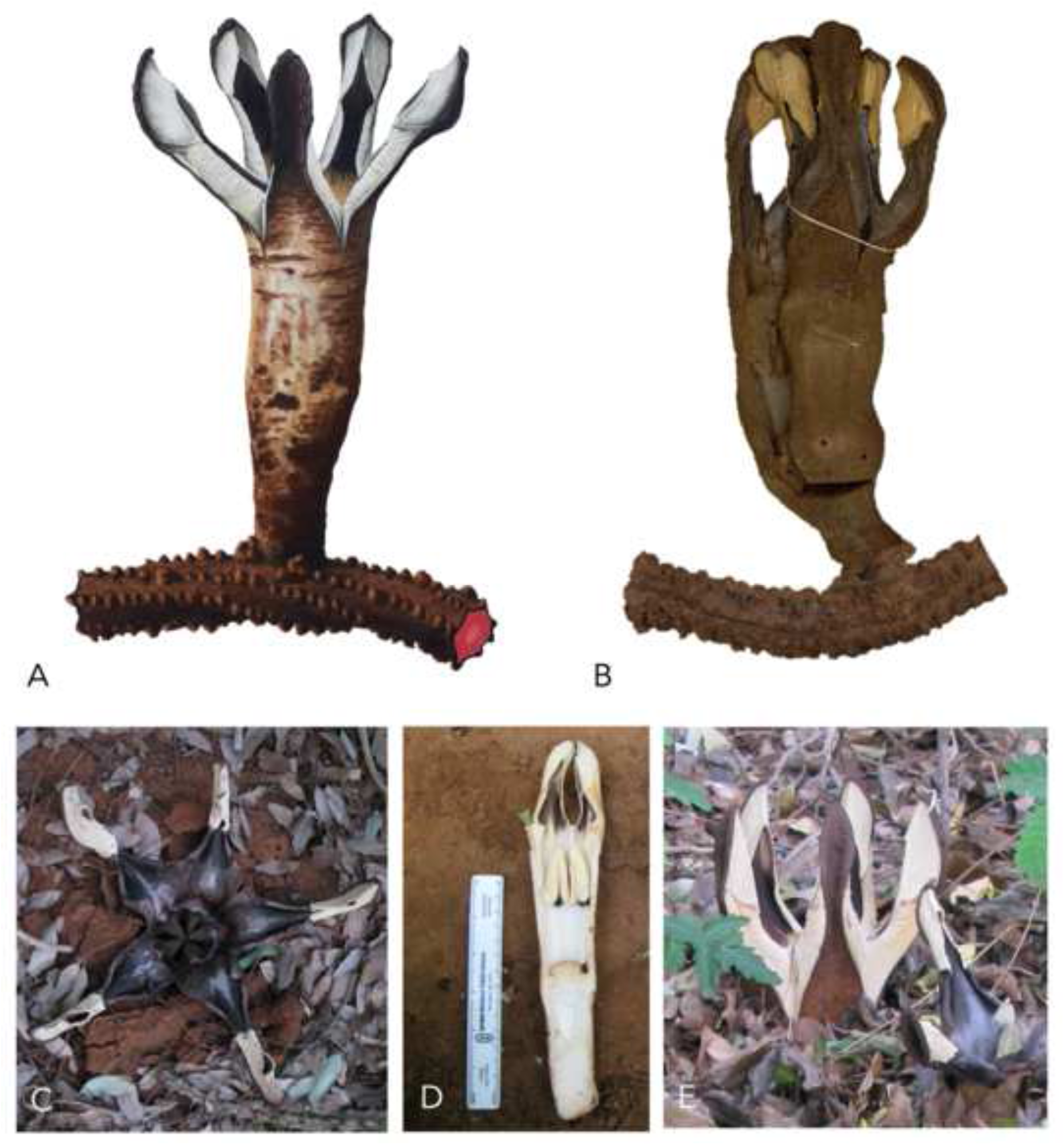
*Hydnora esculenta*. **A** illustration of flower and rhizome; **B** example of herbarium specimen flower and rhizome (both *Humbert* 19805 [P]); **C–E** live flowers. **A** ILLUSTRATED BY SEBASTIAN HATT.

### DISTRIBUTION

Endemic to southern Madagascar, specifically the Atsimo-Andrefana, Anosy and Androy regions. There exists a single report of its occurrence on Réunion Island, although this was later thought to be a short-lived population attached to an introduced host species, *Pithecellobium dulce* (Roxb.) Benth (Bosser, 1994). While there have been no reports of Hydnora from the island since, despite the efforts of Baillon to search for it, it is possible it still grows there and has not been formally recorded (Bosser, 1994). The initial description indicates it was recorded growing in abundance in two separate towns on the island. Although the description provided makes it impossible to confirm whether or not this population was *H. esculenta*, it is perhaps likely given the geographic position of the island relative to Madagascar, and that it was reported on a known host of *H. esculenta*. It is possible that the range of *H. esculenta* is increasing given its ability to parasitise the now widely distributed and planted non-native host *Pithecellobium dulce*, although this has not been studied and is mere speculation at present (Bolin et al., 2013). Map 6.

### SPECIMENS EXAMINED

**MADAGASCAR.** Toliara: NW of Amboasary, Berenty Reserve [25°01’S, 46°19’E], 23 Dec. 1987, *Phillipson* 2716 (P!, MO!, TAN!); Berenty Reserve, Ankoba Forest [25°00.053′′S, 46°17.834′′E], 7 Dec. 2007, *Bolin & Razafindraibe* 71 (TAN!); Anosy Region, Parc Andohahela-Tsimelahy [24°56.186′′S, 46°38.546′′E], 10 Dec. 2007, *Bolin & Razafindraibe* 72 (TAN!); Stream bank in Ampanihy [24°41.808′′S, 44°44.441′′E], 15 Dec. 2007, *Bolin & Razafindraibe* 73 (TAN!); Tuléar, no date, *Richey* 2 (P!); Tuléar, Sudwestliches kustengebiet, 26 Dec. 1959, *Schlieben* 8260 (BR!, M!, G!, BM!, K!, B!); Tuléar, cours de l’Hopital, 1 Jan. 1947, *Humbert* 19805 (epitype P!, isoepitypes G!, K!, MO!); near hospital, 1956, *Abbays* s.n. (TAN!); Sud de Madagascar, 7 Dec. 1910, *Perrier* 8911 (holotype P!, isotype K!).

### HABITAT

Details of habitat preference for *H. esculenta* are scarce. The region of Madagascar in which it grows is sub-arid, dominated by spiny thicket and occasional woodland cover. Its distribution even edges into the transitional-rainforest areas in the south-east of Madagascar (Bolin et al., 2013).

### CONSERVATION STATUS

*H. esculenta* has been described in 2013 as uncommon, but locally abundant in the areas in which it occurred (Bolin et al., 2013). The conservation status of this species remains unclear and largely unstudied. However, there are notable potential threats such as widespread deforestation, with a 45% loss in woodland from 1973 to 2013, particularly in remote regions such as the Tulear province due to ‘clandestine pioneer-agriculture’ (Brinkmann et al., 2014). While there are a few protected areas in southern Madagascar known to be home to *H. esculenta* such as the Berenty Reserve, much the spiny forest habitat in the region remains unprotected and at risk of destruction over the coming years. Furthermore, increased browsing by small livestock has already been noticed by local Tsimelahy villagers as having depleted the *Hydnora* populations (Bolin et al. 2013). As such, the conservation status of *H. esculenta* is perhaps the most concerning within the genus and warrants more attention.

### PHENOLOGY

Flowers primarily from November to January, with fruiting from April to May (Bolin et al., 2013).

### ETYMOLOGY

The specific epithet ‘esculenta’ is latin for ‘edible’. This has roots in the Malagasy name for Hydnora, which is *voantany*. This can be translated and divided into *voa*, meaning ‘fruit’, and *tany*, meaning ‘earth’.

### VERNACULAR NAMES

Hydnora is known as *voantany* in Malagasy, translating into ‘fruit of the earth’ (Bolin et al., 2013).

### ETHNOBOTANY

The fruit is known to be something of a delicacy in the southern regions, apparently delectable with a scent reminiscent of a rennet apple (Jumelle & Perrier, 1912). The rhizomes are sold at local drug markets and have also been recorded as being used by the local semi-nomadic ‘Vezo’ people (fishermen, or those that live from sea fishing) of southern Madagascar in the composition of the ‘mohara’ or ‘mandemilahy’ (Jumelle & Perrier, 1912). This appears to be a fetish primarily made from cow’s horn that is used, likely among other things, to help cure rheumatism (Boiteau, 1997).

### HOST SPECIFICITY

The only confirmed hosts belong to the Fabaceae (Table 1). Confirmed hosts include the native *Albizia tulearensis* R.Vig. and *Alantsilodendron decaryanum* (R.Vig.) Villiers., whose distributions are a close match for *H. esculenta*, and also on the non-native introduced *Pithecellobium dulce* (Roxb.) Benth., whose range is now almost global. Curiously, Bolin et al. (2013) reported that *P. dulce* was the most commonly encountered host in the region. There are historic reports of *Casuarina* sp. as a host (*Schlieben* 8260 [K!]), although this was likely a mistake as this is not in the Fabaceae. *Tamarindus indica* has also been a recorded host in the past, although this was in 1947 [*Humbert* 19805 (P!)] and has not been observed since. However, *T. indica* is within the Fabaceae, and is a recorded host on *H. abyssinica*, so this association is entirely plausible. There is a single report of *H. esculenta* growing on *P. dulce* and *Albizia lebbeck* on Ile de Réunion. Both these trees are non-native, so it is suspected that *Hydnora* was transferred to the island accidentally on the roots of these introduced trees (Bosser, 1994).

### NOTES

Species description adapted from (Bolin et al., 2013). *H. esculenta* is the only species found on Madagascar, and the only species belonging to subgenus *Neohydnora*. It is distinguished from all other species by its unique combination of an angular rhizome with tubercles in parallel lines, together with apical, but not recessed osmophores. This species was described in 1912, although the type specimen is little more than a shattered fragment of fruit. Therefore, it was epitypified and re-described by Bolin et al. in 2013.

**III) Hydnora** subgenus **Dorhyna** *Decaisne* (1873: 75) Harms in Engler (1935: 291). Type: Hydnora abyssinica A.Braun.

*Rhizome* irregular, often both terete, subterete and compressed at different points, rhizome ornamented with numerous lateral tubercles of variable size and shape, randomly and densely distributed on the surface, may occasionally form what appear to be lines. *Flower* usually 4-merous (3 and 5 also observed several times); pedicel absent or present up to 2.5 cm long. *Perianth* external tissues coriaceous, brown to reddish brown; tepal margins with setae; sometimes glabrous outer cucullus present dorso-apical to the apical osmophore and tepal margin. *Osmophore* apical always present but may be reduced; recessed osmophore usually present to at least some extent.

**6. Hydnora abyssinica** *A.Braun* (Braun in Schweinfurth 1867: 217); Decaisne (1873: 76); Martelli (1886: 70); Engler (1895: 169); Rendle (1896: 55); Engler (1900: 386); Baker & Wright in Thistleton-Dyer (1909: 133); Musselman (1997: 18); Beentje & Luke (2002: 1); Bolin et al. (2018: 106). Type: Ethiopia, mountains near Dehli-Dikeno, 1853, *Schimper* 963 (syntypes B [not seen, may be destroyed?], P!).

*Hydnora abyssinica var quinquefida* Engl. (Engler 1900: 386); Baker & Wright in Thistleton-Dyer (1909: 134). Type: Tanzania, Uhehe, Lukosse river, before 1900, *Goetze* 487 (holotype B [not seen, may be destroyed?]).

*Hydnora aethiopica* Decne. (Decaisne 1873: 77); Solms-Laubach in Engler (1901: 6); Baker & Wright in Thistleton-Dyer (1909: 132); (Vaccaneo 1934: 441). Type: Sudan, voyage aux sources du Nil Blanc, before 1873, *Sabatier* s.n. (holotype P).

*Hydnora angolensis* Decne. (Decaisne 1873: 76); Solms-Laubach in Engler (1901: 7); Baker & Wright in Thistleton-Dyer (1909: 134); (Vaccaneo 1934: 444). Type: Angola, ad oram angolensem, before 1873, *collector unknown* s.n. (P [not seen, may be missing?]).

*Hydnora bogosensis* Becc. (Beccari 1871: 6); Martelli (1886: 70); Rendle (1896: 55); Solms-Laubach in Engler (1901: 7); Baker & Wright in Thistleton-Dyer (1909: 134); Chiovenda (1916: 156); Vaccaneo (1934: 432); Harms in Engler (1935: 291); Cufodontis (1972: 36). Type: Eritrea, Keren, July–Aug. 1870, *Beccari* s.n. (holotype FT).

*Hydnora gigantea* Chiov. (Chiovenda 1916: 156); Harms in Engler (1935: 291). Type: Somalia, Berdale to El Ualac, c. 1911, *Paoli* 980, 1035 (both syntypes FT [not seen, may be missing?]).

*Hydnora gigantea var trimera* Chiov. (Chiovenda 1916: 157); Harms in Engler (1935: 291); Cufodontis (1972: 37). Type: as for *Hydnora gigantea* Chiov.

*Hydnora johannis* Becc. (Beccari 1871: 5); Solms-Laubach in Engler (1901: 7); Chiovenda (1916: 156); Vaccaneo (1934: 427); Harms in Engler (1935: 291); Peter (1932: 185); Lebrun (1948: 393); Cufodontis (1972: 37); Troupin (1978: 289); Malaisse (1982: 115); Musselman (1984: 23); Musselman & Visser (1987: 81); Musselman & Visser (1989: 323); Musselman (1993: 31). Type: Eritrea, Keren-Bogos, 1870, *Beccari* 170 (holotype FT!).

*Hydnora johannis var. gigantea* (Chiov.) Vacc. (Vaccaneo 1934: 448); Cufodontis (1972: 37). Type: as for *Hydnora gigantea* Chiov.

*Hydnora johannis var. quinquefida* (Engl.) Solms-Laubach (Solms-Laubach in Engl. 1901: 7); Vaccaneo (1934: 430); Harms in Engler (1935: 291); Peter (1932: 185). Type: as for *Hydnora abyssinica var. quinquefida* Engl.

*Hydnora michaelis* Peter (1932: 185). Type: Tanzania, Usagara, Kidete, Pori on Romuma, before 1932, *Peter* 32690 (holotype B!).

*Hydnora ruspolii* Chiov. (Chiovenda 1917: 57); Vaccaneo (1934: 447); Harms in Engler (1935: 291); Cufodontis (1972: 37). Type: Somalia, Marro Umberto I ville, April 1893,

*Ruspoli & Riva* 1091 (syntypes F!, RO!, G!).

Herbaceous subterranean perennial root holoparasite, without leaves or stem. *Rhizome* irregular, often both terete, subterete and compressed at different points, 5 – 8(– 10) cm in diameter; rhizome surface coriaceous, dark brown, lighter coloured (when fresh) near growth tip; rhizome spreads laterally and often bi- or trifurcating or branching irregularly to form smaller terminal branches; rhizome ornamented with numerous lateral tubercles of variable size and shape, randomly and densely distributed on the surface, not in clear consistent lines (although may occasionally form what appear to be lines), tubercles remain same or develop into flower or develop into haustorium; numerous flowers and flower buds on single rhizomes; rhizome fleshy, brick red to reddish pink internal tissue when broken, producing sticky, astringent exudate, lighter colours at growing tip (in life). *Flower* usually 4-merous (3 and 5 also observed several times); flower emerges only partially from soil; two floral chambers, androecial chamber subtended by gynoecial chamber, inner surfaces of chambers glabrous, pedicel absent or present up to 2.5 cm long. *Perianth* external tissues coriaceous, brown to reddish brown, often with thin fleshy white streaks where the outer layer has cracked, total length (5 –)11 – 25(– 33) cm; perianth tissues fleshy; perianth tube and tepal cavity pinkish to ivory white inside (in life), darkening to brick red-brown (dried), 3 – 5.5 cm wide; tepal margin reddish pink to pink-orange inside (in life), darkening to brick-red brown, tepal (measured from apex to point of connation with adjacent tepal) (4 –)6 – 9(– 14) cm long × 1.7 – 3.8 cm wide (measured at midpoint); tepals lanceolate to spathulate, initially erect, gently curved and connivant, later spread out flat on ground, this change hastened by moist weather but usually inevitable with age; tepal margins covered in diffuse setae, longest on dorsal side, often entirely absent on ventral side, from 1 – 7 mm, generally quite variable between individuals. *Osmophore* apical osmophore present, border between osmophore and hairy tepal margin sloping down basally in a sharp diagonal from the dorsal side to the ventral side; recessed osmophore present but barely so, usually extending no more than 1 cm down the length of the tepal margin; spongy, cream to white (in life), 2.6 – 5 cm long × 1.6–2 cm wide, darkening to tan when dried, generating very foetid odour. *Androecium* antheral ring formed by connate w-shaped anther lobes, forming a central orifice, transversely striate and divided into numerous horizontal pollen sacs. *Gynoecium* ovary inferior, unilocular with numerous ovules produced from apical placenta, ovary 2.2 – 2.8(– 4.5) cm wide; lobed and cushion like stigma on the floor of the gynoecial chamber. *Fruit* subterranean, globose, 10 – 15 cm in diameter, leathery brown pericarp. *Seed* spherical, black-brown 1 – 1.8 mm, thousands of seeds embedded within mealy white pulp. Fig. 11.

**Fig. 11.**
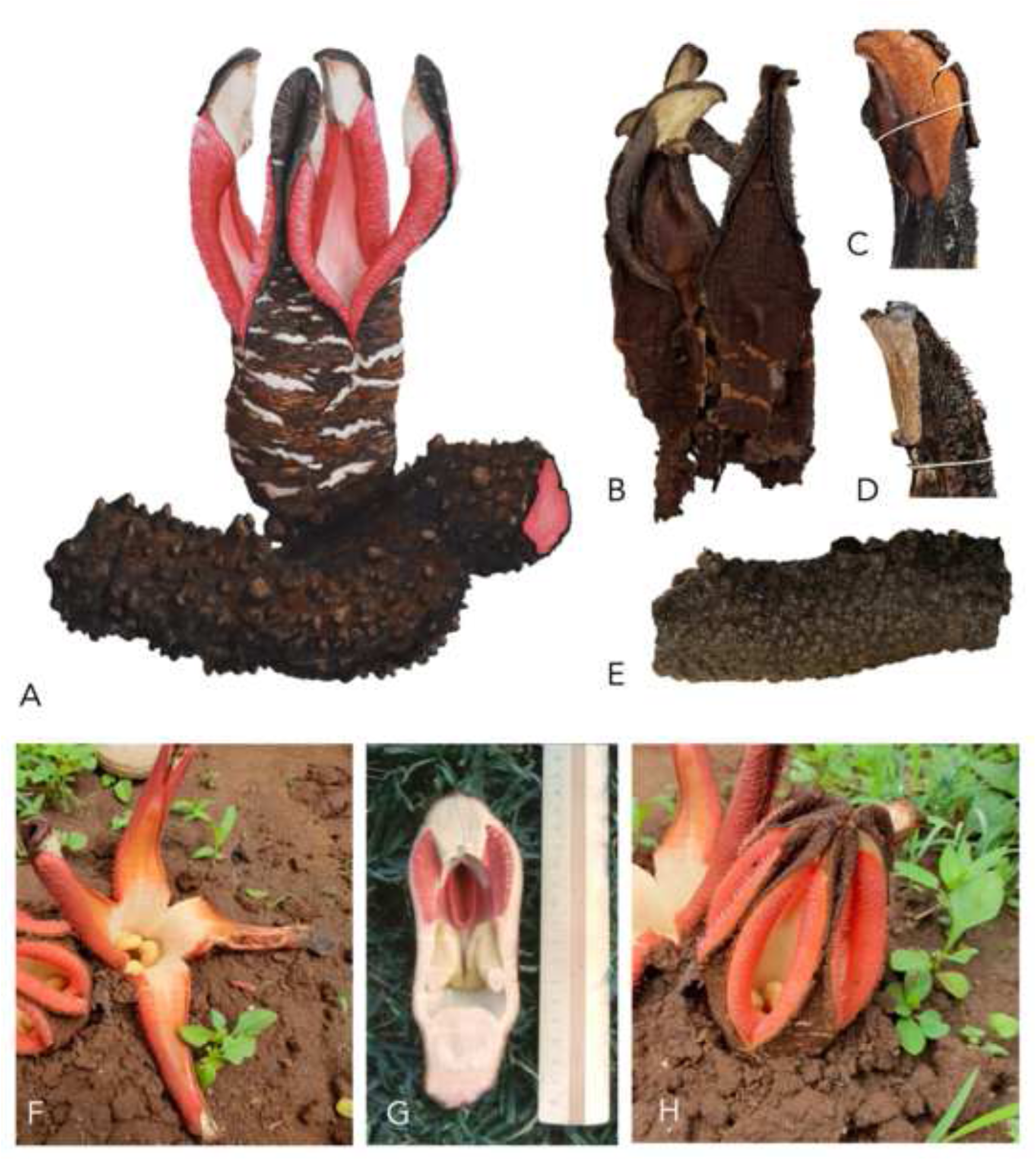
*Hydnora abyssinica*. **A** illustration of flower and rhizome; **B–E** examples of herbarium specimens: **B** flower (*Luke* 3016 [K]); **C** osmophore (*Greenway & Kirrika* 11132 [K]); **D** osmophore and diffuse tepal setae (*Gillett* 14151 [K]); **E** rhizome (*Jameson & Michelmore* 134 [K]); **F–H** live flowers. **A** ILLUSTRATED BY SEBASTIAN HATT. PHOTOS: **F & H** MATHEW REES; **G** WILLIAM BURGER, slide no. 4174 [K].

### DISTRIBUTION

*H. abyssinica* has a considerably larger distribution than all the other species. It has been recorded in northern South Africa, Eswatini, Mozambique, Tanzania, Kenya, Uganda, the DRC, Rwanda, Ethiopia, Somalia, Eritrea and Nigeria. There is a single report from 1873 of a section of rhizome collected from Gabon, although this material has since been lost (Decaisne, 1873). However, given the recent discovery of *H. abyssinica* in Nigeria, far outside the previously accepted range, it is possible that *Hydnora* may be found in Gabon or surrounding countries such as the Central African Republic or Chad (Agyeno et al., 2018). Map 7.

### SPECIMENS EXAMINED

**NIGERIA.** LGA Plateau State: Nekong Kanke, c. 2018, *Gosomji* s.n. (JUHN!). **ERITREA.** Keren, July 1870, *Beccari* s.n. (holotype FT); Abita, Keren, July 1870, *Beccari* 170 (holotype FT!); Barka Valley, Keren-Bogos, Sciotel?, c. 1870, *Beccari* 417, 423, 424, 3685, 3687, 3690 (FT [not seen, may be missing?]); Sciotel, c. 1870, *Martelli* s.n. (FT!); Beni-Amer, Monte Damba, 25 Sep. 1903, *Pappi* 6110 (FT!); Habab, Melchet-Tzaroba, 18 April 1909, *Pappi* 8222 (FT!). **SOMALIA.** Somaliland: Meid, April 1875, *Hildebrandt* s.n. (K!, BM!, K!, L!); West of Gacan Libaax camp on road, 17 May 2017, *Awale* AIA20 (HARG!); Hargeisa, 26 Sep. 1932, *Gillett* 4071 (K!, FT!); Northern rangelands [9°58’N, 46°12’E], 11 March 1981, *Beckett* 929 (EA!); Hahi, June 1885, *James & Thrapp* s.n. (K!); Harrar, da Gildessa a Zeila, 1889, *Robecchi-Bricchetti* 184 (FT!); Northeast Region: Sultanato di Obbia, 26 April 1924, *Puccioni & Stefanini* 443 (FT!); Central Region: Marro Umberto I ville, April 1893, *Ruspoli & Riva* 1091 (syntypes FT!, RO!, G!); Harli Bomani, April 1893, *Ruspoli & Riva* s.n. (G!); South Region: Berdale to El Ualac, c. 1911, *Paoli* 980 (syntype FT [not seen, may be missing?]); Berdale to El Ualac, c. 1911, *Paoli* 1035 (syntype FT [not seen, may be missing?]). **ETHIOPIA.** Oromia Region: Harar, slopes W of Midaga above Gobelli River Valley [42°4’E, 8°46’N], 9 May 1964, *Burger* 3496 (K!, US!); Pozzi di el Banno, 2 May 1939, *Corradi* 2866 (FT!); Road Ginir, 119 km E of Ginir, 17 Nov. 2015, *Friis, Abebe & Getachew* 15708 (K!); Sidamo, 20 km S of Neghelle, 18 May 1982, *Friis, Tadesse & Vollesen* 3101 (K!); Sidamo, El Siro water holes, 22 May 1983, *Gilbert & Ensermu* 7727 (K!); Kilil, Lake Margerhita, 5 April 1958, *Eriksson* 645 (S [not seen, identity not confirmed]); Amhara Region: Gondaraba nei dintorni del fortino, 28 May 1939, *Corradi* 2862 (FT!); mountains near Dehli-Dikeno, 1853, *Schimper* 963 (syntypes B [not seen, may be destroyed], P!); Dehli- Dikeno, near Dschadscha, Agau, 19 July 1853, *Schimper* 1189 (P!); Tigray Region: Amhara-Uolcait, 5 Feb. 1913, *Pappi* 9007 (FT!); SNNP Region: Balkakà-Neldà, 1893, *Ruspoli & Riva* 1623 (FT!); Dire Dawa Region: 20 km from Haresa, 26 April 1969, *Ebba* 734 (K!); 27 km NE of Dire Dawa on road to Djibouti, 15 April 1972, *Gilbert* 2390 (K [appears to be missing], EA!); Somali Region: 22 km from Qarsonney, 14 May 2006, *Thulin et al.* 11140 (UPS); Unlocalised: likely 1854, *Schimper* s.n. (P! [P05591526]); 4 April 1854, *Schimper* s.n. (P! [P05591525]). **KENYA.** K1: Balambala, no date, *Adamson* 450 or *Bally* 6151 (K!, G!, S); a few miles S of Bura, 3 Jan. 1943, *Bally* 2050 (K!, EA!); Mandera, 30 km on Ramu-Malka Mari road [4°4’N, 40°59’E], 6 May 1978, *Gilbert & Thulin* 1534 (K!, EA!, UPS); Moyale, 15 km out on Wajir road, 6 Nov. 1952, *Gillett* 14151 (K!, EA!); Kilifi, Sala E [3914E, 0306N], 28 Dec. 1991, *Luke* 3016 (K!, EA!); Mandéra, Oct. 1889, *Sacleux* 965 (P); K2: Munyen, Turkana, March 1947, *Bally* 5042 (K!); Wei Wei, Katuw, 23 July 1978, *Meyerhoff* 71M (K!); K3: Baringo Distr., Lake Hannington E shore, 28 June 1972, *Bally* 15177 (EA!); Baringo Distr., Chemolingot borehole area [0°58’N, 35°57’E], 6 Sep. 1976, *Timberlake* 712 (EA!); Rumarute, 1 Nov. 1937, *Searle* s.n. (K!); K4: Nairobi, Feb. 1964, *Beecher* s.n. (K!, EA!); Nairobi National Park, just outside Cheetah Gate [1°26’S, 36°58’E], 26 April 1975, *Friis & Hansen* 2605 (K!); Machatos Distr., Kanza, 14 Nov. 1982, *Howard* 16717 (EA!); K6: Kampi ya Bibi [1°33’S, 36°33’E], 25 April 1965, *Archer* 13150 (EA!); Ngong Hills, Nov. 1938, *Bally* 12514 (K!); Nairobi-Magadi Rd, prehistoric site, 13 Dec. 1968, *Early* 14080 (EA!); Chyulu Hills, 18 Nov. 1972, *East African Herbarium* 15231 (EA!); Kajaido Distr., Chyulu Hills [2°31’S, 37°45’E], 18 Oct. 1969, *Gillett & Kariuki* 18797 (K!, EA!); Chyulu Hills, Mbirikani-Nbi [02320S, 3731E], 12 Dec. 1991, *Luke* 2963 (K!, EA!); Chyulu Hills, below Bonham House [0232S, 3744E], 22 Dec. 2000, *Luke & Luke* 7182 (K!, EA!, MO, US!); Olorgasailie Prehistoric Site, 28 Dec. 1958, *Napper* 1220 (K!, EA!); Loita Hills [0140S, 3555E], 26 April 1948, *Vesey Fitzgerald* 6359 (K!); K7: no locality, Feb. 1977, *Adamson* 16079 (EA!); Voi, Dec. 1962, *de Jong* 362/CL (EA!); Mackinnon road, 14 Nov. 1950, *Lt. Col. Venour* 114/50 (EA!); Mt. Kisigau, Nov. 1938, *Joanna* 8635 (K!, EA! [both just rhizomes, identity impossible to confirm]). **UGANDA.** U1: N. Karamoja, near Kaabong, June 1945, *Dale* 416 (EA!); Lira, Lango, Dec. 1929, *Davies* 1038 (K!); Amudat, Karamoja, May 1948, *Eggeling* 5812 (K!); 30– 50 miles N of Kacheliba–Karamoja, 9 May 1953, *Padwa* 106 (K!, EA!); Lochoi, Karamoja, 24 May 1940, *Thomas* 3539 (K!); U2: Kijumbura, Victoria Nile, 31 Jan. 1907, *Bagshawe* 1559 (BM!); Kasumba, Ankole, 9 April 1953, *Jameson & Michelmore* 134 (K!); Ruizi river, 24 Feb. 1951, *Jarrett* 444 (K!, EA!); U4: Sekamuli, near Bamunanika [0°41’N, 32°36’E], 13 June 1967, *Ferreira* 133 (K!, EA!, MO); Unlocalised: 28 April 1997, *Badaza* 64808 & 64809 (K!). **RWANDA.** Gabiro, no date, *Hoier* s.n. (BR); Kibugabuga [Bugesera], 17 Nov. 1953, *Liben* 953 (BR); Akagera National Park [1.5088°S, 30.6814°E], 9 Jun 2018, *Luke* 18431 (EA!). **TANZANIA.** T1: Mwanza Distr., Ukiriguru, 7 Dec. 1970, *Ebbela* 71/1 (EA!); Musoma, below Sid Downies Dam, 30 Dec. 1962, *Greenway & Harvey* 10917 (K!, EA!); Mwanga, Ukiriguru, 15 Jan. 1948, *Prentice* 1298 (EA!); Benagi Hill, Kikima, 24 March 1961, *Pullman* 12381 (EA!); Bukoba Distr., Nov. 1983, *Kanywa* 84 (K! [just rhizome, impossible to confirm identity]); T2: Kilimanjaro, N of Loloari Hill, 26 Nov. 1967, *Bigger* 1442 (EA!); Lake Manyara National Park, near Mto wa Mbu river, 6 Jan. 1962, *Dingle* HD1002 (EA!); Arusha, Ngorongoro Conservation Area [3°20’S, 35°2’E], 15 Oct. 1993, *Ellemann* 654 (AAU, MO!); Mbuka Distr., Tamamgwe river, 3 Jan. 1959, *Mahinde* 415 (EA!); T3: Msasa, Lake Manyara National Park, 4 Dec. 1963, *Greenway & Kirrika* 11132 (K!, EA!); Usambara Mts, May 1998, *Schlage* CS265 (B!); Muhesa, 4 April 1924, *Shantz* 198 (K!); T5: Kongogo, 8 Jan. 1966, *Newman* 88 (EA!); Landschaft Ussagara, Kidete, Dec. 1925, *Peter* 32690 (holotype B!); Kondoa Distr., Kondoa, 11 Jan. 1962, *Polhill & Paulo* 1139 (K!, B!, BR!); T6: between Uhehe and Khutu, Jan. 1899, *Goetze* 395 (B [not seen, may be destroyed]); T7: Uhehe, Lukosse river, no date, *Goetze* 487 (holotype B [not seen, may be destroyed]); Unlocalised: Serengeti Park, near a river, 2004, *Vandervinne* s.n. (photograph, BR!). **MALAWI.** Mchiru Mts, Nov. 1989, *Jenkins* 87 (K!). **ANGOLA.** Ad oram Angolensem, before 1873, *collector unknown* s.n. (holotype P [not seen, may be missing and may not be *H. abyssinica*). **NAMIBIA.** Hardap Region: Karibib, Farm Nuiskasaken, 24 July 1963, *Häblich & Giess* 7652 (M!). **SOUTH AFRICA.** KwaZulu-Natal Prov.: Tembe Park [27°2’40”S, 32°25’6”E], 12 Dec. 2001, *Vahmeijer* HV00385 (PRE!). **UNLOCALISED.** Voyage aux sources du Nil Blanc, before 1873, *Sabatier* s.n. (holotype P); no date, *Bouxin* 1257BIS (BR); no date, *Musselman* 198 (E). **HABITAT.** Wide range of arid to sub-arid habitat types following the distribution of its generally *Acacia* host species. Recorded from dry woodland, wooded grassland, bushland, scrubland, often on black cotton, clay or sandy alluvial soils or rocky slopes (Musselman, 1997; Beentje & Luke, 2002).

### CONSERVATION STATUS

*H. abyssinica* has a vast range across Africa, although the population density within that range is unclear. The species as a whole is unlikely to be threatened given the size of the distribution, although certain localities will be more at risk than others. The population in Nigeria is reported to be under considerable threat from over-harvesting and land development in the area (Otuwose Agyeno, pers. comm.). Similarly, in Uganda there are concerns about increasing awareness of its medicinal potential leading to over-harvesting (Nyafuono, 2000). The South African Red List records *H. abyssinica* as ‘Not Threatened’, although nearby populations in Mozambique are known to be harvested to a considerable degree (Williams et al., 2008; 2011). It is difficult to comment with confidence on the threat harvesting causes, as the full extent of the populations are largely unknown. Furthermore, *H. abyssinica* may later prove to be composed of distinct species or subspecies, some of which may be under more threat than others.

### PHENOLOGY

Flowering time varies considerably across its vast range but tends to occur following the rains and lasts for a few days to a month (Beentje & Luke, 2002). Fruits tend to mature around five months after flowering (Musselman, 1997).

### ETYMOLOGY

The specific epithet refers to the locality of the type specimen in Ethiopia (formerly known as Abyssinia).

### VERNACULAR NAMES

In Somalia, it is known as *likke*, *lécche*, *lipti* or *leka* (Vaccaneo, 1934) or *laka* (*Gilbert & Thulin* 1534 [K!, EA!, UPS]) or *liki*, *like* or *dingah* (Mandu et al., 1999) or *likaha* (*Ebba* 734 [K!]). In Gobato, Galla and Malakati, spoken in Ethiopia, it is known as *tuka* (*Ebba* 734 [K!]; *Adamson* 450 [K!, G!, S]). In East Africa, it has names in many languages. In Maasai or Maa it is known as *orukunyi* (*Luke & Luke* 7182 [K!, EA!, MO, US!]; Beentje & Luke, 2002) or *erukunyi* or *erkyunyi* (Mandu et al., 1999). In Borana it is known as *toga* (*Gilbert & Thulin* 1534 [K!, EA!, UPS]; Mandu et al., 1999). In Burji it is known as *guli* (Mandu et al., 1999). In Kikuyu it is known as *muthigira* (Mandu et al., 1999). In Pokot it is known as *aurieng’o* or *kaworiongo* (Mandu et al., 1999). In Turkana it is known as *auriong’o* (Mandu et al., 1999). In Swahili it is known as *kioga* (Engler, 1900) or *nyambo* or *mnyambo* (in Tanzania according to *Goetze* 395 [B]; Mandu et al., 1999). In Uganda, in Luganda, it is known as *omutimagwensi* or *omutimagwettaka* (Nyafuono, 2000) or *mutima gwenai/gwensi* which translates to ‘the heart of the earth/world’ (*Bagshawe* 1559 [BM!]; *Kanywa* 84 [K!]) or *awuriongo* (*Thomas* 3539 [K!]) or in Ankole, spoken in Uganda, it is known as *otumimawasi* (*Jarrett* 444 [K!, EA!]). In Tanzania, it is known also known as *mulumbi* (*Goetze* 487 [B]; Vaccaneo, 1934) or *amamaso* (*Newman* 88 [EA!]). In Wasukuma, spoken in Tanzania, it is known as *wija wa hasi*, which translate to ‘half-blind person of the earth’, in reference to its ‘screwed-up peering look’ when it first breaches the ground surface (*Prentice* 1298 [EA!]). In Nigeria, the Ngas and Hausa speaking community know it as *kurshim* or *kaushekasa* (Agyeno et al., 2018). In southern Africa, in Xitsonga/Xichangana, it is known as *mavumbule* (Williams et al., 2011). It is known as *uMavumbaka* in Zulu, meaning ‘the one that pops up’, a name that is shared with *Sarcophyte sanguinea* (Williams et al., 2010).

### ETHNOBOTANY

*H. abyssinica* has a wide range of uses across its range. In Somalia, locals cut it into thin slices and heat it in water for a few hours, obtaining a strongly coloured liquid use to dye mats (Vaccaneo, 1934). The fleshy rhizome and fruits are known to be eaten in Somalia (*Ruspoli & Riva* 1091 [FT!, G!, RO!]; Vaccaneo, 1934). The fruit are reported to be cooked like a baked potato, then cut open with the seeds eaten with sugar and milk (*Gillett* 4017 [K!]). In Ethiopia, the fruit is reported to be eaten by local people (*Burger* 3496 [K!, US!]). In Kenya, the flower buds are reported to be eaten by the Pokot and Maasai (*Gillett* 14151 [K!, EA!]; *Timberlake* 712 [EA!]), with additional minor medicinal use during childbirth (Beentje & Luke, 2002) or to treat post-partum haemorrhage (*Searle* 2575 [K!]). In northern Uganda it is used as an antidiarrheal agent, while in eastern and central Uganda, the root is used to treat cardiovascular disorders and diabetes (Nyafuono, 2000). One report indicates that in Uganda, if it is used to treat heart infections, as reported by Nyafuono (2000), the sufferer who used this treatment must for-go goat hearts for the rest of their life (*Bagshawe* 1559 [BM!]). In Tanzania, it is used in the treatment of sore throat, swollen tonsils and throat inflammations (*Schlage* CS265 [B!]; Nyafuono, 2000). Furthermore, in Tanzania it is eaten with milk by herd boys, who also use the flower buds as a small cup to milk the cows in the daytime, although apparently this is strictly prohibited (*Elleman* 654 [AAU, MO!]). The usage of *H. abyssinica* in local medicine in Nigeria far pre-dates its discovery there in 2018. The plant is harvested at considerable rates for use by doctors in the treatment of diarrhoea, dysentery, stomach-ache, piles and erectile disfunction, with some locals reported using it to ‘rejuvenate the body from the physical and mental effects of the sun’ (Agyeno et al., 2018). It is sold at traditional medicine markets in South Africa, although harvested from Mozambique, in the treatment of bleeding, diarrhoea, acne, piles, menstrual problems, stomach cramps (Williams et al., 2010; 2011). *H. abyssinica* has been reported being consumed by wild pigs and warthogs in East Africa (*Bally* 12514 [K!]; *Bally* 5042 [K!]; Beentje & Luke, 2002). Beyond uses, *H. abyssinica* has been reported causing damage to local streets, driveways and patios in Ethiopia, where its rhizomes have burst through the road or tile surface (Maass & Musselman, 2001).

### HOST SPECIFICITY

*H. abyssinica* appears to be restricted to Fabaceae hosts, mostly *Senegalia* and *Vachellia* (*Acacia*), but also a few other trees from different genera in the same family (Table 1). The following hosts have been reported: *Albizia lebbeck* (L.) Benth., *Delonix regia* (Bojer ex Hook.) Raf., *Piliostigma thonningii* (Schumach.) Milne-Redh., *Senegalia asak* (Forssk.) Kyal. & Boatwr [*Acacia glaucophylla*], *Senegalia mellifera* (Benth.) Seigler & Ebinger [*Acacia mellifera*], *Senegalia nigrescens* (Oliv.) P.J.H.Hurter [*Acacia nigrescens*], *Tamarindus indica* L., *Vachellia amythethophylla* (Steud. ex A. Rich.) Kyal. & Boatwr [*Acacia macrothyrsa*], *Vachellia drepanolobium* (Harms ex Y.Sjöstedt) P.J.H.Hurter [*Acacia drepanolobium*], *Vachellia elatior* (Brenan) Kyal. & Boatwr. [*Acacia elatior*], *Vachellia gerrardii* (Benth.) P.J.H.Hurter [*Acacia gerrardii*], *Vachellia grandicornuta* (Gerstner) Seigler & Ebinger [*Acacia grandicornuta*], *Vachellia hockii* (De Wild.) Seigler & Ebinger [*Acacia hockii*], *Vachellia nilotica* subsp. tomentosa (Benth.) Kyal. & Boatwr. [*Acacia arabica*], *Vachellia tortilis* (Forssk.) Galasso & Banfi [*Acacia tortilis*], *Vachellia xanthophloea* (Benth.) Banfi & Galasso [*Acacia xanthophloea*]. An entry from iNaturalist cited a *Prosopsis* sp. (Fabaceae) as the host [*bryanadkins* 13223800]. Given the rhizome appeared excavated in the image, this could indeed be the host. There are a handful of other host species that have been recorded in the past on old herbarium specimens, although the validity of these is doubtful given the ease with which the host species can be mistaken. As they belong to completely different families and have only been reported once, they are not considered potential hosts here. These erroneous hosts are *Adansonia digitata* (Malvacaea), *Dobera glabra* (Salvadoraceae), *Hildebrandtia* sp. (Convolvulaceae), *Kigelia africana* (Bignoniaceae) and *Terminalia sericea* Burch. Ex DC. (Combretaceae).

### NOTES

Species description adapted from (Beentje & Luke, 2002). *H. abyssinica* is the most widespread and variable species of *Hydnora*, hence the numerous synonyms described since its initial publication in 1867. *H. abyssinica* was previously known as *H. johannis*, Becc., until Beentje & Luke (2002) noticed that an earlier description for *H. abyssinica* was indeed valid. As such, many specimens around the world are still labelled as *H. johannis*. The various synonyms have been studied to as much detail as possible in this monograph, despite the loss of many type specimens and obscurity of many of the protologues. None of them appeared to be sufficiently distinct from *H. abyssinica* besides *H. solmsiana*, which has been reinstated here. Further field collections in under-studied areas may well provide new specimens that may lead to *H. abyssinica* being divided further.

*H. abyssinica* is also similar to *H. hanningtonii*, but is distinguished by having a proportionately smaller apical osmophore that only occupies considerably less than ½ the tepal. *H. hanningtonii* has dense strigose setae that cover the entire tepal margin and a red perianth tube and tepal cavity, while *H. abyssinica* has longer setae, that rarely reach the ventral side of the tepal margin, and a pinkish-white perianth tube and tepal cavity.

**7. Hydnora hanningtonii** Rendle (1896: 55); Solms-Laubach in Engler (1901: 7); Baker & Wright in Thistleton-Dyer (1909: 133); Chiovenda (1916: 156); Vaccaneo (1934: 448); Harms in Engler (1935: 291); Cufodontis (1972: 37). Type: Tanzania, Jordan’s Nullah, Lake Victoria, E arm of Mwanza Gulf, Dec. 1882, *Hannington* s.n. (syntype BM!) & Ethiopia, Ginia, Galla highlands, 1895, *Donaldson Smith* s.n. (syntype BM [not seen, likely missing]).

*Hydnora arabica* Bolin & Musselman (2018: 1). Type: Oman, Dhofar Region, Ayn Ayuoon, 0.66 km SE of the spring [17.241294, 53.891644], 16 Dec. 2014, *Bolin, Rahbi & Musselman* JFB2014OM3 (holotype OBG!, isotype US!).

*Hydnora cornii* Vacc. (Vaccaneo 1934: 414); Harms in Engler (1935: 291); Cufodontis (1972: 37). Type: Somalia, Mogadishu, Villa Governationale di Afgoi, *Corni* s.n. (holotype TOM).

Herbaceous subterranean perennial root holoparasite, without leaves or stem. *Rhizome* terete, subterete or compressed, 1–5 cm diam., may be much larger on mature specimens (not seen); rhizome surface coriaceous, dark brown to light tan, lighter coloured (when fresh) near growth tip; rhizome spreads laterally and may bifurcate or branch irregularly; rhizome ornamented with numerous lateral tubercles, randomly and densely distributed on the surface, not in clear consistent lines, tubercles remain same or develop into flower or develop into haustorium; numerous flowers and flower buds on single rhizomes; rhizome swollen and irregular at haustorial interface with host root; rhizome fleshy, pink to red internal tissue when broken, lighter colours at growing tip (in life). *Flower* usually 4-merous, (though 3 and 5 observed); flower emerges only partially from soil; two floral chambers, androecial chamber subtended by gynoecial chamber, inner surfaces of chambers glabrous. *Perianth* external tissues coriaceous, brown to reddish-brown, total length 10–28 cm; perianth tissues fleshy; perianth tube red inside (in life), darkening to brick red-brown, 1.9–3.6 cm wide; tepal margins red inside (in life), darkening to brick red-brown, tepal (measured from apex to point of connation with adjacent tepal) 4.0–8.5 cm long × 1.9–3.5 cm wide (measured at midpoint) (this on 4-merous flowers, wider on 3-merous flowers and narrower on 5-merous flowers); tepals clavate to elongate-linear, typically curved, not fused at apex; tepal margins always with dense strigose setae covering most or all of the tepal margin from ventral to dorsal edge, max. 3.3 mm. *Osmophore* apical and very prominent, greater than or equal to ½ length of tepal, spongy and white (in life), darkening to tan when dried, generating foetid odour, described as a strong ammoniac smell. *Androecium* antheral ring formed by connate w-shaped anther lobes, forming a central orifice, transversely striate and divided into numerous horizontal pollen sacs. *Gynoecium* ovary inferior, unilocular with numerous ovules, ovary 2.5–4.5 cm wide; lobed and cushion-like stigma on the floor of the gynoecial chamber, stigma 1.9–2.2 cm wide. *Fruit* only one observed, subterranean, globose, 7 cm in diameter. *Seed* spherical, black-brown, 0.7–1.2 mm, thousands of seeds within fruit. Fig. 12.

**Fig. 12.**
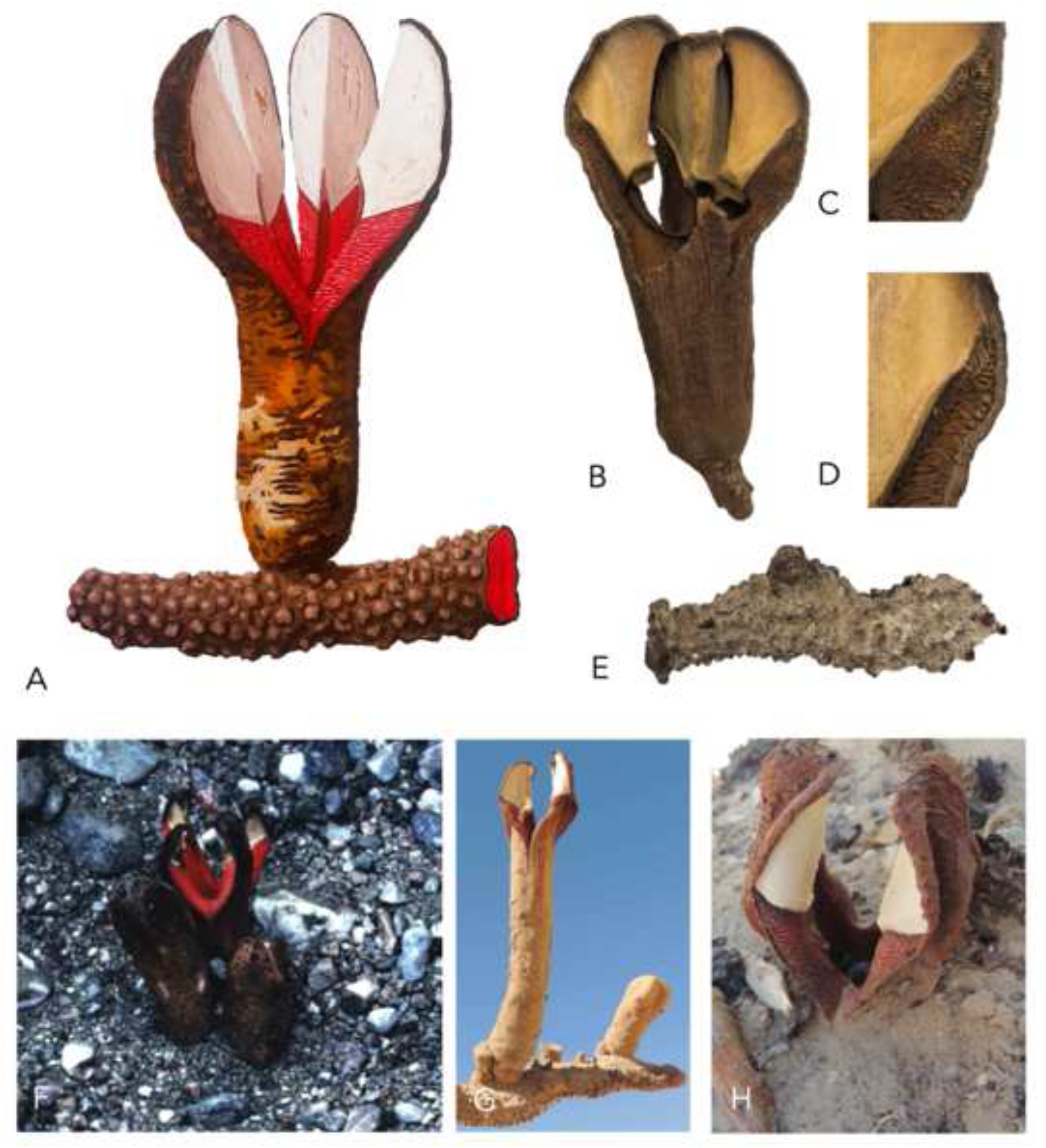
*Hydnora hanningtonii*. **A** illustration of flower and rhizome; **B–E** examples of herbarium specimens (all *Thesiger* s.n. [BM]): **B** flower; **C, D** strigose tepal setae; **E** rhizome; **F–H** live flowers. **A** ILLUSTRATED BY SEBASTIAN HATT.

### DISTRIBUTION

Known from the southern Arabian Peninsula (recorded from southwestern Saudi Arabia, southern Oman [Dhofar region], and Yemen). Also recorded from East Africa (Somalia, Tanzania and Kenya). Collected from an elevation of 200–1900 m. Map 8.

### SPECIMENS EXAMINED

**SAUDI ARABIA.** Najran Region: ∼42 km NW of Najran, 30 April 1979, *Collenette* 1485 (K!). **OMAN.** Dhofar Region: 50 km W of Mudhai, 13 Sep. 1985, *Miller* 7619 (E!, ODU!); Wadi Haluf, SE of Rub el Khali, 8 Nov. 1945, *Thesiger* s.n. (BM!); Umshadid, 31 Oct. 1946, *Thesiger* s.n. (BM!); Ayn Ayuoon, 0.66 km SE of the spring [17°14’28.66”N, 53°53’29.92”E], 16 Dec. 2014, *Bolin, Rahbi & Musselman* JFB2014OM3 (holotype OBG!, isotype US!); ∼ 2 km S of Haluf, Wadi Haluf [17°20’3.56”N 53°57’35.82”E], 16 Dec. 2014, *Bolin, Rahbi & Musselman* JFB2014OM2 (OBG!); ∼17 km NE of Mirbat, Wadi Ayn [17° 7’2.64”N 54°47’53.21”E], 15 Dec. 2014, *Bolin, Lupton, Musselman, Rahbi* JFB2014OM1 (OBG!); ∼16 km NE of Mirbat, 17 Dec. 2014, *Bolin, Lupton, Musselman, Rahbi* JFB2014OM4 (OBG!). **YEMEN.** Hadramawt, Wadi Masila, 14 April 1947, *Thesiger* s.n. (BM!). **SOMALIA.** Central Region: Mogadishu, Villa Governationale di Afgoi, before 1932, *Corni* s.n. (holotype TOM); Afgoi, 6km from Mogadishu, 13 Aug. 1959, *Moggi & Bavazzano* s.n. (FT!). **KENYA.** K7: Taita Taveta Distr., Ndara Ranch to Kajeri [0330S, 3937E], 26 Dec. 1991, *Robertson* 6537 (K!, EA!). **TANZANIA.** T1: Jordan’s Nullah, Lake Victoria, E arm of Mwanza Gulf, Dec. 1882, *Hannington* s.n. (syntype BM!); T3: near Gonja, Jan. 1950, *Bally* 7694 (K!).

### HABITAT

In the southern Arabian Peninsula, it is typically found in stony and sandy wadi beds. It has been observed growing in sand. The type locality of *H. arabica* is mountainous with strongly dissected rocky plateaus and loamy-skeletal to sandy-skeletal shallow soils underlain by Tertiary, Hadhramaut group formations: beige bioclastic, calcarenitic and micritic limestone with abundant fossilised molluscs (Bolin et al., 2018). In East Africa, it also appears to have a degree of preference for sandy soils, as in (*Robertson* 6537 [K!, EA!]). In Somalia, it has been reported from a sand dune (*Robecchi-Bricchetti* 184 [FT!]). *H. cornii* was described growing in an *Albizia* grove in the garden of the Consulate’s villa, though it is unclear how sandy the soil is there. The paucity of collections and infrequent sightings of this species mean that our understanding of its habitat or ecological requirements are limited.

### CONSERVATION STATUS

Fieldwork in Dhofar, Oman by Bolin et al. (2018) has indicated that populations of both *H. hanningtonii* and its host *Acacia tortilis* are abundant in the wadi beds, despite being difficult to locate due its hypogeous habit. They suggest that the conservation status of *H. hanningtonii* in southern Oman is secure. Details of its abundance and distribution in Yemen are unclear. The conservation status in East Africa is unknown, although the ability of the plant to grow on artificially planted, introduced species, as seen in Afgoi where it grows on an *Albizia* grove in the Consulate’s villa garden (Vaccaneo, 1934), is encouraging as this suggests it can survive disruption to the natural habitat so long as a suitable host is present.

### PHENOLOGY

Observed flowering in southern Oman in December, but local villagers report that flowering also occurs in other months (Bolin et al., 2018). At some sites, flowering was reported in July and at others in May. In East Africa, flowering has also been reported around December and January, although in Somalia it has been reported flowering in April. The apparently sporadic flowering may be dependent on adequate rainfall.

### ETYMOLOGY

The specific epithet refers to the collector of the type, Bishop Hannington, whom Rendle apparently named this species after.

### VERNACULAR NAMES

Known in Oman as *dhanuna*, *xamleg* (Jibbali), *khamlayyeh*, *khumla’ah* (Dhofari) (Bolin et al., 2018). Known in Yemen as *nabeekh*, *fateekh* and *tarateef* (Lawdar and Dathina) (Al-Fatimi et al., 2016). Vernacular names in East Africa for this species are likely the same as *Hydnora abyssinica* (see Vernacular Names), as there is currently no evidence that local people distinguish between these two species.

### ETHNOBOTANY

In Oman, Jibbali settlers in Dhofar use all parts of the plant, and typically collect it following a rainstorm (Miller & Morris, 1988). The flower heads are eaten, which smell of strong ammoniac and taste like very green cheese. However, the fruit is the more desired part for consumption. The matured fruit is apparently very sweet and is enjoyed by both humans and foxes. The dried remnants of the plant are used by locals as an indicator of where to dig for it the next time the rains fall. The rhizomes are crushed to a rough paste, which has been used to treat odorous or crusted leather. It has also been used to strengthen cotton material in fishing nets and ropes. Both the paste and dried out remnants have been used as a black ‘antimony’ powder which can serve as a dye or as eye shadow. In South Yemen, in the Lawdar and Dathina districts of Abyan, local people collect the flowers after the rains (Al-Fatimi et al., 2016). They eat the flowers, fresh or grilled, and use them in medicinal therapy to cure various stomach diseases, gastric ulcers and cancers. The dried flowers are ground down and applied with milk or water to the afflicted area. The rhizomes and fruits are not known to be used in Yemen. In Kenya it has been recorded being dug up by rhinoceroses (*Bally 7694* [K!]). Uses in East Africa for this species are likely the same as *Hydnora abyssinica* (see Ethnobotany entry), as there is currently no evidence that local people distinguish between these two species.

### HOST SPECIFICITY

All known hosts are within the Fabaceae (Table 1). In Oman and Yemen, the primary host is *Vachellia tortilis* (Forssk.) Galasso & Banfi (*Acacia tortilis*) (Bolin et al., 2018). However, Bolin et al. (2018) found a robust population of *H. arabica* growing on *Pithecellobium dulce* (Roxb.) Benth. in a goat yard. Goats feeding on in the nearby wadi where *H. hanningtonii* is abundant are thought to be the likely vector for the *Hydnora* seeds into the goat yard. This tree is native to the Americas and has been introduced into much of Africa and Asia. Similarly, the type of *H. cornii* (now *H. hanningtonii*) was found on *Albizia lebbeck*, another introduced tree.

### NOTES

Species description adapted from Vaccaneo (1934) and Bolin et al. (2018). *H. arabica* was described in 2018 and was thought to be restricted to the southern Arabian Peninsula. However, examination of the protologues and type specimens of *H. hanningtoni* Rendle and *H. cornii* Vaccaneo revealed that both species share the same morphological characters as *H. arabica*; notably an apical osmophore that is equal to or greater than ½ the length of the tepal, dense strigose setae that cover most or all of the tepal margin, and an entirely red perianth tube. As *H. hanningtoni* Rendle is the oldest synonym described, it is selected here as the true name for this species. According to Art. 60.12 of the International Code of Nomenclature (Turland et al., 2018), the specific epithet is changed to *hanningtonii*, as is appropriate for an epithet named in honour of a male person. Examination of herbarium material revealed that several specimens from Tanzania, Kenya and Somalia previously labelled as *H. abyssinica* were in fact *H. hanningtonii*. Furthermore, a set of photographs online by Jabruson of (supposedly) *H. abyssinica* growing on *Acacia* roots in Kenya, Tsavo National Park, were identified as *H. hanningtonii* (https://www.mindenpictures.com/stock-photo/fly-on-flower-of-(hydnora-abyssinica)-parasite-growing-on-acacia-roots-tsavo/search/detailmodal-0_90179754.html). Morphologically, this species is quite similar to *H. abyssinica*, which can make identification from dried specimens relatively difficult, though not impossible as there are clear characters that reliably separate the two species.

**8. Hydnora solmsiana** *Dinter* (1909: 57); Marloth (1913: 178); Dinter (1923: 424); Harms in Engler (1935: 291); Schreiber in Merxmüller (1968: 41). Type: Namibia, Windhoek, Klein– Windhoek, before 1909, *Dinter* 356 (syntype B [not seen, likely destroyed]) & Namibia, am auß bei Keetmanshoop, before 1909, *Dinter* s.n. (syntype B [not seen, likely destroyed]). Neotype: Namibia, Horn River at Warmbad Township, no date, *Bayer* s.n. (neotype selected here PRE!, isoneotypes K!, US!, LD, HUH!).

Herbaceous subterranean perennial root holoparasite, without leaves or stem. *Rhizome* irregular, terete, subterete and often compressed, 1.5 – 6.5 cm in diameter, rhizome surface coriaceous, dark brown; rhizome spreads laterally and often bi- or trifurcating or branching irregularly to form smaller terminal branches; rhizome ornamented with numerous lateral tubercles of variable size and shape, randomly and densely distributed on the surface, not in clear consistent lines (although may occasionally form what appear to be lines), tubercles remain same or develop into flower or develop into haustorium; numerous flowers and flower buds on single rhizomes; rhizome fleshy, pinkish internal tissue when broken. *Flower* usually 4-merous (rarely 3– or 5–merous); flower emerges only partially from soil; two floral chambers, androecial chamber subtended by gynoecial chamber, inner surfaces of chambers glabrous. *Perianth* external tissues coriaceous, rust-brown, total length 10 – 18 cm; perianth tissues fleshy; perianth tube usually creamish-white inside, darkening to brick red brown (dried), 2 – 4.5 cm wide; tepal lobes creamish-white (in life), darkening to brick-red brown (dried), tepal (measured from apex to point of connation with adjacent tepal) 3.5 – 10 cm long × 1.2 – 3 cm wide (measured at midpoint); tepals lanceolate to spathulate, gently curved and connivant; tepal margins covered in diffuse setae, longest on dorsal side, less dense on ventral side, c. 2 mm long, generally quite variable between individuals. *Osmophore* apical osmophore present, with an indistinct border between osmophore and setose tepal margin, roughly horizontal or sloping up apically from dorsal side to the ventral side; recessed osmophore present, reaching down to basal end of setose tepal margins and often slightly beyond, elliptic or spathulate, often fleshy and pitted in life; both osmophores visible in dried specimens as distinctly paler regions; no unpleasant smell reported. *Androecium* antheral ring formed by connate w-shaped anther lobes, forming a central orifice, transversely striate and divided into numerous horizontal pollen sacs. *Gynoecium* ovary inferior, unilocular with numerous ovules produced from apical placenta, ovary c. 1.5 – 2 cm wide; lobed but nearly flat and cushion-like stigma on the floor of the gynoecial chamber. *Fruit* subterranean, globose, c 4.5 cm in diameter, leathery brown pericarp. *Seed* spherical, black-brown, c. 1 mm, thousands of seeds embedded within mealy pink-white pulp. Fig. 13.

**Fig. 13.**
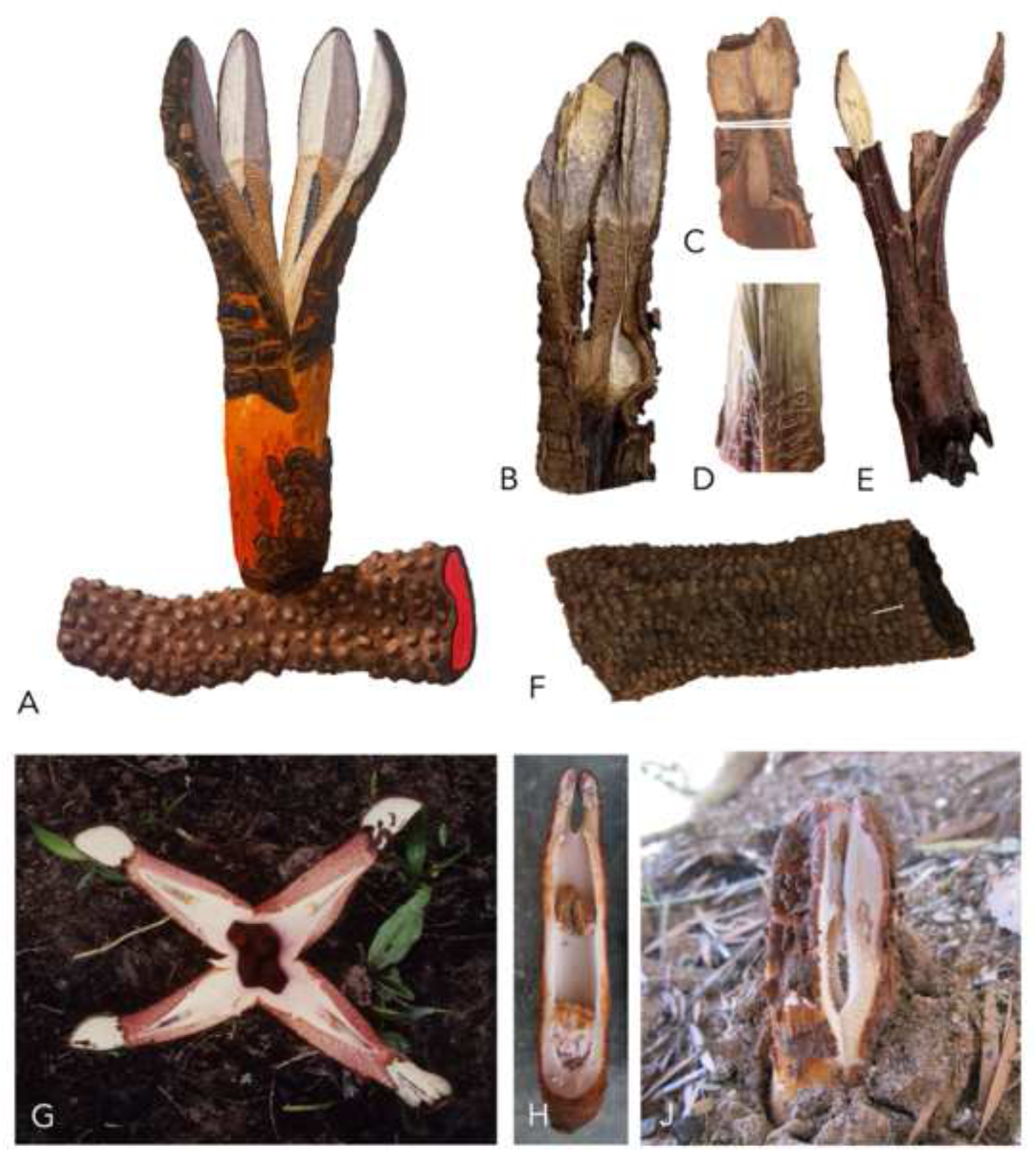
*Hydnora solmsiana*. **A** illustration of flower and rhizome; **B–F** examples of herbarium specimens: **B** tepals (*Kotze & Giess* 10892 [WIND]); **C** tepal showing apical and recessed osmophore (*Bayer* s.n. [US]); **D** indistinct border between osmophore and tepal setae region (*Bolin* 09-06 [WIND]); **E** flower (*Bolin* 09-06 [WIND]); **F** rhizome (*Musselman* 6279a [BM]); **G** open flower (*Malaisse* 11290 [BR]); **H** flower cross-section; **J** freshly-opened flower. **A** ILLUSTRATED BY SEBASTIAN HATT.

### DISTRIBUTION

*H. solmsiana* has been recorded from Namibia, Botswana, South Africa, Eswatini, Zimbabwe, the DRC, Sudan, Rwanda and western Tanzania. Given how patchy this known distribution is, it is likely that *H. solmsiana* also grows in Angola and Zambia but has not been reported yet. *H. solmsiana* appears to be distributed more across southern and central Africa, while *H. abyssinica*, *H. hanningtonii* and *H. sinandevu* tend to be found more across East Africa. Map 9.

### SPECIMENS EXAMINED

**SUDAN.** Central Sudan: Gezira, Um Barona, 27 Aug. 1982, *Musselman* 6129 (K, E); Blue Nile Prov., Abu Naama, no date, *Musselman* 6228 (E!); Khartoum, S of Khartoum, Dom Island, 4 Jan. 1989, *Musselman* 6279A (K!, E!, BM!, BR!, LD!). **TANZANIA.** T4: Katisunga, 19 Jan. 1950, *Bullock* 2273 (K!). **DEMOCRATIC REPUBLIC OF THE CONGO.** Katanga: Dikulushi, 20 km N of Kiankalamu, likely 1980, *Malaisse* 11290 (BR!); Dikulushi, 15 Dec. 1980, *Malaisse* 11348 (BR!). **ZIMBABWE.** Midlands: Gwelo, Gwelo Teacher’s College, 21 Feb. 1967, *Biegel* 1929 (K!, SRGH!); Selukwe, Uoncima Farm, 2 Jan. 1965, *Guy* 159891 (K!, SRGH!); Matabeleland South: Filabusi, 26 Dec. 1962, *Guy* 139469 (K!, SRGH!, BR!); Matabeleland North: Bulawayo, no date, *Musselman & Obilana* s.n. (M!, ODU!). **BOTSWANA.** Ngamiland: Okavango, no date, *Smith* s.n. (ODU!); Okavango, Qhusai Island, 25 Nov. 1979, *Smith* 2904 (PSUB!); Ngamiland, Aug. 1946, *Wilmot* PRE52465 (PRE!); Unlocalised: July 1994, *Bolnick* 22 (MO). **NAMIBIA.** Karas Region: Horn River at Warmbad, no date, *Bayer* s.n. (neotype K!, isoneotypes PRE!, US!, LD, HUH!); Gondwana Canyon Park [S27 24.157, E17 56.768], 11 Feb. 2009, *Bolin* 09- 06 (WIND!); Am Auß bei Keetmanshoop, no date, *Dinter* s.n. (B [not seen, likely destroyed?]); Bethanie, Plaas Umub, 13 April 1970, *Kotze & Giess* 10892 (WIND!); Gobas, 27 Aug. 1927, *collector unknown* s.n. (NBG!); Hardap Region: Maltahöhe, Bergzebrapark Naukluft, 24 Feb. 1974, *Giess & Robinson* 13256 (WIND!); Maltahöhe, Bergzebrapark Naukluft, 3 Sep. 1972, *Merxmüller & Giess* 28200a (M!); Khomas Region: Windhoek, Klein Windhoek, no date, *Dinter* 356 (syntype B [not seen, likely destroyed]); Hatsamas, 6 March 1911, *Dinter* 1951 (B [not seen, likely destroyed], SAM!); Hereroland, near Windhoek, Jan. 1912, *Bohr* 5093 (PRE!); Erongo Region: 10 km E of Uis [S21 13.454, E15 03.295], 2 Feb. 2009, *Bolin* 08-1 (WIND!); Uis [S21 13.454, E15 03.295], 31 Oct. 2008, *Bolin* 08-1-2 (WIND!); Otjozondjupa: Otavi, Grootfontein, no date, *Freyer & Giess* 7469, 7650 (WIND [not found, likely missing]); Okahandja, no date, *Gaerdes & Giess* 7651 (WIND [not found, likely missing]); Tsumkwe, no date, *Kat* 807 (WIND [not found, likely missing]); Tsumkwe, edge of Nama Pan, no date, *Story* 5162 (WIND!); Grootfontein, no date, *Bauer* s.n. (WIND [not found, likely missing]); Kavango West: Okavango Mile 46 [18°18’22”S, 19°15’3”E], 19 Feb. 2003, *Strohbach* BS5635 (WIND!); Kavango East: Okavango, Hukapama [17°53’116”S, 20°15’26”E], 7 Nov. 1996, *Burke* 95373 (WIND!); Rundu, along Kavango River [S17 53.164, E20 15.461], 3 May 2009, *Bolin* 93-9 (WIND!); Oshikoto: Halali, no date, *Musselman & Visser* s.n. (ODU!); Namutoni, no date, *Musselman & Visseri* s.n. (ODU!); Etosha Pan, Tsumeb, Namutoni, 29 Dec. 1986, *Dujardin* 10491 (M!); Oshana: Etosha National Park, Okaukuejo Rest Camp, 20 Nov. 2005, *Bolin* 05-2 (WIND!). **ESWATINI.** Nsoko, lowveld of Swaziland, Feb. 1977, *von Wissel* s.n. (K!, PRE!). **SOUTH AFRICA.** Limpopo Prov.: Transvaal, Naboomspruit, 12 Jan. 1939, *Galpin* s.n. (PRE!); Transvaaal, Potgietersrust, Jan. 1940, *McDonald* 213 (K!); Potgietersus Distr., Tugela Hotel, 3 Jan. 1978, *Muller* s.n. (PRE!); Besjeskuil, before 1931, *van der Byl* 8829 (K!); Northern Cape Prov.: Namaqualand, Richtersveld, Sendelingsdrift, 10 Jan. 1996, *Williamson* 5882 (PRE!); Little Namaqualand, banks of Orange River, Arris Drift [Beesbank], 31 Aug. 1925, *Marloth* 12526 (PRE!).

### HABITAT

Appears to have some degree of preference for riverside habitats, although not exclusively. In Namibia, it has often been collected and seen by the authors near rivers (*Bolin* 939 [WIND!]; *Bayer* s.n. [PRE!, US!, LD, HUH!, K!]; *Williamson* 5582 [PRE!]), with Dinter describing several observations along rivers in Namibia (Dinter, 1909). Similarly in Sudan it has been collected along the Blue Nile (Musselman, 1984) and in Botswana in riverine woodland in the Okavango delta (*Smith* 2904 [PSUB!]). However, it is not necessarily restricted to this habitat, and has been collected from several non-riparian habitats across its range.

### CONSERVATION STATUS

Due to the poor understanding of this species and the lack of recognition of it as distinct species over the last century, its conservation status is currently unknown. It has several reported uses across its range, although the extent of harvesting and the damage this is causing to population levels has not been ascertained. Given it appears to have a number of possible hosts and a distribution that spreads across several countries, it is hoped that the plant is unlikely to be critically threatened, though may face local threats at a population level.

### PHENOLOGY

In southern Africa, it is often recorded flowering in December or January following the rains (Marloth, 1913; pers. obs.), with many specimens being collected around December and January. Therefore, across its vast range, flowering time varies considerably.

### ETYMOLOGY

Dinter does not explain his choice of specific epithet, although it is likely that he named it in honour of the German botanist Hermann zu Solms-Laubach, who wrote about the seeds of the Hydnoraceae and Rafflesiaceae (Solms-Laubach, 1874) and produced a short monograph on the Hydnoraceae in 1901 for Adolf Engler’s Pflanzenreich (Solms-Laubach, 1901).

### VERNACULAR NAMES

Vernacular names for *H. solmsiana* may apply to other species of *Hydnora*, particularly *H. abyssinica* to which it bears close resemblance. In southern Africa, in Xitsonga/Xichangana, it is known as *mavumbule* (Williams et al., 2011). It is known as *uMavumbaka* in Zulu, meaning ‘the one that pops up’, a name that is shared with *Sarcophyte sanguinea* Sparrm. (Williams et al., 2010). In Sudan, a common name for the plant is *tartous*, *tartousch* or *dambu*, deriving from the Arabic word *utartis*, meaning ‘to hold fast’ (Musselman, 1984).

### ETHNOBOTANY

In Sudan, the dried rhizomes are used as a charcoal/fuel alternative. It is reported to be far superior to normal charcoal, producing a heat of unrivalled intensity and evenness (Musselman, 1984). Also in Sudan, the plants are powdered and boiled to produce a tannin-rich astringent paste used in the treatment of diarrhoea and assorted intestinal ailments (Musselman, 1984). The white pulp of the fruit is also eaten as a delectable food source in their own right and is reported to resemble custard apple (*Annona squamata*) in taste and an apple in texture (Musselman, 1984). Curiously, a large shipment of *H. solmsiana* [then *H. abyssinica*] was seized from the Germans during the First World War, although the purpose of this shipment remains unknown (Dinter, 1909; Musselman, 1984). Perhaps it is linked to its high tannin content and subsequent use in tanneries before the war (Musselman & Visser 1987). In the Democratic Republic of the Congo, the rhizomes are crushed into a decoction and used in the treatment of swollen legs and feet (Malaisse, 1982). This decoction is also apparently poured on the soil of *Cassava* fields in order to ensure good harvest and restore soil fertility. *H. solmsiana* is consumed as food by a wide range of animals, including monkeys and goats along the Blue Nile in Sudan (Musselman, 1984; Musselman & Visser, 1989) and elephants at Etosha Pan National Park (Musselman & Visser, 1989). Beyond uses, *H. solmsiana* has been reported causing damage to local streets, driveways and patios in both Namibia and Sudan, where its rhizomes have burst through the road or tile surface, even causing cracks in the walls of a nearby house (Maass, 2001).

### HOST SPECIFICITY

*H. solmsiana* is restricted to Fabaceae and has several overlapping host species with *H. abyssinica* (Table 1). So far, it has only been recorded on the following hosts: *Acacia saligna* (Labill.) H.L.Wendl. [*Acacia cyanophylla*], *Faidherbia albida* (Delile) A.Chev. [*Acacia albida*], *Senegalia nigrescens* (Oliv.) P.J.H.Hurter [*Acacia nigrescens*], *Senegalia polyacantha* subsp. *ampylacantha* (Hochst. Ex A.Rich.) Kyal. & Boatwr [*Acacia polyacantha subsp. campylacantha*], *Vachellia karroo* (Hayne) Banfi & Galasso [*Acacia horrida*, *Acacia karroo*], *Vachellia luederitzii* (Engl.) Kyal. & Boatwr. [*Acacia luederitzii*], *Vachellia nebrownii* (Burtt Davy) Seigler & Ebinger [*Acacia nebrownii*], *Vachellia nilotica* (L.) P.J.H.Hurter & Mabb. [*Acacia nilotica*], *Vachellia reficiens* (Wawra & Peyr.) Kyal. & Boatwr. [*Acacia reficiens*], *Vachellia rehmanniana* (Schinz) Kyal. & Boatwr [*Acacia rehmanniana*], *Vachellia seyal* (Delile) P.J.H.Hurter [*Acacia seyal*], *Vachellia sieberiana* (DC.) Kyal. & Boatwr. [*Acacia sieberiana*], *Vachellia tortilis* (Forssk.) Galasso & Banfi [*Acacia tortilis*], *Vachellia tortilis* subsp. *heteracantha* (Burch.) Kyal.& Boatwr [*Acacia heteracantha*, *Acacia litakunensis*]. It is likely not limited to the hosts above, particularly because one of the hosts, *Acacia saligna*, is an introduced species from Australia. Dinter records *Acacia horrida* as the host in the protologue, although this is likely a misidentification of *Acacia karroo*; a mistake made by many collectors in Namibia at that time (Namibia Biodiversity Database, 2021).

### NOTES

Species description adapted from (Dinter, 1909; Marloth, 1913). *H. solmsiana* was described in 1909 and was accepted and considered a valid species over the 25 years that followed. The monographs of Vaccaneo (1934) and Harms (1935) then expressed doubt about the strength of the distinction between *H. solmsiana* and *H. abyssinica* (then *H. johannis*). It was scarcely mentioned again until 1989, when Musselman officially synonymised it with *H. abyssinica* (then *H. johannis*) (Musselman & Visser, 1989). The uncertainty stems from the brevity of Dinter’s original description, which omits several key details about the defining morphological characters, and lacks a clarifying illustration (Dinter, 1909).

Although the description is brief, it does mention that *H. solmsiana* is parasitic on *Acacia karroo* (*Acacia horrida*), and that the flower is 4-merous, with a whitish pink interior and a subterete rhizome. This is barely sufficient as a distinguishing description and makes no reference to the important character of the apical osmophore. Unfortunately, the type specimen was almost certainly destroyed in Berlin during the Second World War as it is no longer found in the Berlin herbarium, making it difficult to be without any doubt that the name: ‘*H. solmsiana* Dinter’ indeed applies to the specimens described here. However, while Dinter’s original 1909 description is brief, its contents do agree with the characters described here as belonging to the species, and *H. solmsiana* is by far the most dominant species of *Hydnora* with terete rhizomes in Namibia. More thorough descriptions and illustrations with associated herbarium specimens are later provided by Marloth in 1913, and in ‘Flowering Plants of South Africa’ by Pole-Evans in 1931, accompanied by a colour illustration by Cynthia Letty. Both of these concur with, and expand on, the original description by Dinter. The specimens of *H. solmsiana* cited in this paper, along with several iNaturalist observations, have consistent morphological differences from *H. abyssinica* and these characters match those in the combined descriptions of Dinter, Marloth and Pole-Evans. As a result, the species name *H. solmsiana* Dinter is reinstated here. Molecular data would provide further confidence to this taxonomic decision.

As a result of the destruction of the type in Berlin, a neotype was designated here. (*Bayer* s.n. [PRE!, K!, US!, LD, HUH!) was selected as it has multiple duplicates and the specimens are in good condition, clearly illustrating the morphological characters of the species.

*H. solmsiana* is most similar to *H. abyssinica*, from which it is distinguished by having a conspicuous recessed osmophore that extends at least as far as the basal end of the tepal margins, and an indistinct border between the apical osmophore and the tepal margins that often slopes up apically from the dorsal side to the ventral side. While colour is not a reliable trait, the tepal margins are generally whitish-cream when fresh, particularly in southern Africa.

Examination of specimens of *H. abyssinica* across Africa revealed that many of these are in fact *H. solmsiana* and clearly display traits associated with that species. These specimens were mostly in southern or central African countries and not East African countries. It appears that the distribution of *H. solmsiana* and *H. abyssinica* are largely distinct, with *H. solmsiana* generally west of the Great Rift Valley, and *H. abyssinica* to the east. The two populations co-exist in the Kruger National Park in South Africa, although seem to maintain morphological distinction. Malaisse (1982) wrote that the specimens of *Hydnora abyssinica* [then *H. johannis*] he observed in the Democratic Republic of the Congo, Dikulushi, may in fact be *H. solmsiana*, and indeed he was correct.

Curiously, there is a single specimen of what appears to be *H. abyssinica* in central Namibia, which is unexpected as it is far from the distribution of other *H. abyssinica* (*Häblich & Giess* 7652 [M!]) in an area apparently dominated by *H. solmsiana*. It is also possible, and perhaps likely given the distribution, that the population of *H. abyssinica* reported in Nigeria is indeed *H. solmsiana*, given its proximity. However, the images published in Agyeno et al. (2018) are not sufficient to determine this.

**9. Hydnora bolinii** *Hatt* **sp. nov.** Type: Ethiopia, Harar Province, Schevelli Valley, flat area above river, 10 km E of the Fafan Valley [42°40’E, 9°16’N], 1432 m, 14 April 1963, *Burger* 2695 (holotype K!; isotype US!).

Herbaceous subterranean perennial root holoparasite, without leaves or stem. *Rhizome* irregular, often both terete, subterete and compressed at different points, up to 3 cm in diameter, though likely has the potential to be larger; rhizome surface coriaceous, dark grey-brown; rhizome spreads laterally; rhizome ornamented with numerous lateral tubercles of variable size and shape, randomly and densely distributed on the surface, not in clear consistent lines (although may occasionally form what appear to be lines), tubercles remain same or develop into flower or develop into haustorium; numerous flowers and flower buds on single rhizomes; rhizome fleshy, internal colour not seen. *Flower* usually 4-merous (3 and 5 also observed several times); flower emerges only partially from soil; two floral chambers, androecial chamber subtended by gynoecial chamber, inner surfaces of chambers glabrous. *Perianth* external tissues coriaceous, rust-brown, total length up to 16 cm, may grow even larger; perianth tissues fleshy; perianth tube salmon and white inside, up to 3 cm wide; tepal salmon and white inside, tepal (measured from apex to point of connation with adjacent tepal) 5 – 10 cm long × 2 – 3 cm wide (measured at midpoint); tepals lanceolate, gently curved and connivant; tepal apex tissue glabrous outer cucullus and apparently not an osmophore; tepal margins setose, setae c. 2 mm long, tepal margin only reaching half way from ventral to dorsal side of tepal. *Osmophore* positioned in two places; apical osmophore a narrow upside-down V-shaped strip, apical to the tepal margin and basal to the outer cucullus; recessed osmophore in tepal cavity; unknown if smell is produced. *Androecium* antheral ring formed by connate w-shaped anther lobes, forming a central orifice, transversely striate and divided into numerous horizontal pollen sacs. *Gynoecium* ovary inferior, unilocular with numerous ovules produced from apical placenta, ovary c. 1.5 – 2.5 cm wide; lobed and cushion-like stigma on the floor of the gynoecial chamber. *Fruit* and *Seed* not seen. Fig. 14.

**Fig. 14.**
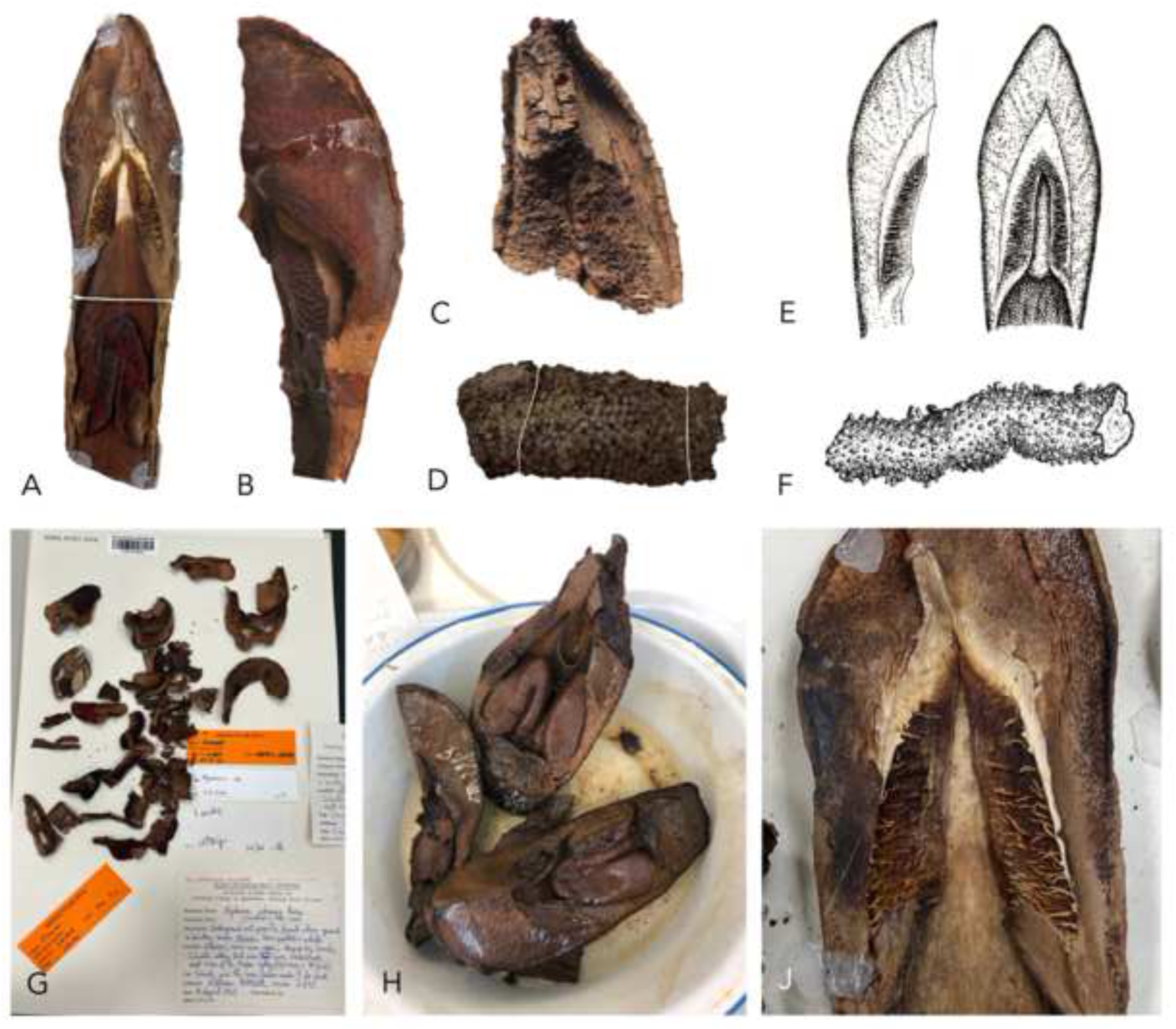
*Hydnora bolinii*. **A–D** examples of herbarium specimen material; **A** tepal (*Gillett* 12781 [K]); **B** tepal, side view (*Burger* 2695 [US]); **C** very desiccated tepal apex (*Gillett & Watson* 2358 [K]); **D** rhizome (*Gillett* 12781 [K]); **E** tepal illustration, side view (left), front view (right); **F** rhizome illustration; **G, H** type specimen (*Burger* 2695 [K]): **G** herbarium sheet; **H** spirit specimen; **J** close up of apical osmophore, reduced tepal margins and recessed osmophore (*Gillett* 12781 [K]). **E, F** DRAWN BY SEBASTIAN HATT.

### RECOGNITION

*H. bolinii* is similar to members of subgenus *Dorhyna*, but is distinguished by having an apical osmophore reduced to a narrow strip (vs apical osmophore as the actual apex of the tepal), together with a recessed osmophore that extends to the basal end of the tepal margins (as seen in *H. solmsiana* but not *H. abyssinica*). The setose tepal margin never touches the dorsal edge of tepal (vs *H. abyssinica* and *H. solmsiana* where it touches the dorsal edge for the entire length of the tepal below the osmophore). There is a glabrous outer tissue that forms a large glabrous cucullus at the tepal apex (this tissue is not present in *H. abyssinica* or *H. solmsiana*).

### DISTRIBUTION

Africa: Ethiopia, northern Kenya, Somalia. The true extent of *H. bolinii* is likely greater than we currently understand given the limited number of collections made. Map 10.

### SPECIMENS EXAMINED

**SOMALIA.** Somaliland: N Somalia site A/9 [9°50’N, 45°33’E], 23 June 1981, *Gillett & Watson* 2358 (K!, EA!). **ETHIOPIA.** Harar Prov., Schevelli Valley [42°40’E, 9°16’N], 14 April 1963, *Burger* 2695 (holotype K!, isotype US!). **KENYA.** K1: Dandu [3°26’N, 39°54’E], 14 April 1952, *Gillett* 12781 (K!).

### HABITAT

Collected in rich *Commiphora*-Acacia *scrub* scattered with larger trees such as *Delonix*, *Terminalia* and *Gyrocarpus* (*Gillett* 12781 [K!]) and in degraded open *Acacia* woodland on eroded yellow-red silty sand (*Gillett* & *Watson* 2358 [K!, EA!]). The limited number of collections likely do not represent the full range of this species’ preferred habitat.

### CONSERVATION STATUS

Due to the limited number of collections of this species and the lack of recognition of *H. bolinii* as distinct species over the last century, its conservation status is currently unknown. The very small number of collections and observations made of this species may be due to its rarity, in which case it is of likely conservation concern. However, this may simply be an artefact of it growing in fairly remote and inaccessible locations. As such it is difficult to comment on how threatened this species is, though perhaps much of what has been written about *H. abyssinica* (see species entry) applies here.

### PHENOLOGY

Specimens collected in flower in April and June. (*Gillett* 12781 [K!]) reports that it rained on 27^th^ March 1952, and it was collected on 14^th^ April that year. This may suggest that like other *Hydnora* species, it often flowers following rains.

### ETYMOLOGY

This species is named in honour of Dr Jay Bolin, in recognition of his vast contribution to the study of *Hydnora* over almost two decades of intrepid field work and innovative research.

### VERNACULAR NAMES

Known as *tukha* in Boran and *liki* in Somali (*Gillett* 12781 [K!]). It is likely that many of the vernacular names for *H. abyssinica* also apply to this species, as it is currently unknown whether local people differentiate between the two species.

### ETHNOBOTANY

The fruit is reported to be eaten by people (*Gillett* 12781 [K!]). (*Burger* 2695 [K!]) writes that it is ‘dug up by Somalis’. It is likely that many of the uses of *H. abyssinica* also apply to this species, as it is currently unknown whether local people differentiate between the two species.

### HOST SPECIFICTY

No confirmed host known (Table 1). Reported to grow amongst *Acacia*, *Commiphora*, *Delonix*, *Terminalia*, *Gyrocarpus*, *Balanites, Boscia, Ziziphus, Zygophyllum & Salsola* (*Gillett* & *Watson* 2358 [K!]; *Gillett* 12781 [K!]), so the host is likely one of these. (*Burger* 2695 [K!]) describes it as ‘under *Acacia*’, although this single report is not sufficient to be confident that this is the true host. Further collections are required. Given its apparent relatedness to *H. abyssinica* based on morphology, it is perhaps likely that it parasitises *Acacia* (*Vachellia* or *Senegalia*), though this is not possible to confirm with the data available.

### NOTES

Three specimens were united in showing the same set of striking morphological differences in the tepals from any other *Hydnora* species. They are distinguished by the large glabrous outer cucullus, in combination with a reduced setose tepal margin that does not touch the dorsal edge of the tepal, a reduced apical osmophore to a small upside-down v-shape above the tepal margin, and an enlarged recessed osmophore. It is described here as a new species, *H. bolinii*, and placed within subgenus *Dorhyna* due its terete rhizome with randomly distributed tubercles, presence of an apical and recessed osmophore, and its probable Acacia host. Due to the very small number of known specimens, molecular data and further collections/observations in the field would provide considerable confidence to this taxonomic decision.

Certain characters, such as the glabrous outer cucullus, are shared with *H. sinandevu*, while others, such as the setose tepal margin, are shared with *H. abyssinica* and *H. solmsiana*. No other species has the apical osmophore reduced to a narrow strip above the tepal margin, and no other species has a setose tepal margin that does not touch the dorsal edge of the tepal. Both osmophores appear as distinctly paler regions in herbarium specimens, and are thus immediately recognisable. Unfortunately, there are currently no known photographs of a live specimen.

**IV) Hydnora** subgenus **Sineseta** *Hatt*, **subg. nov.** Type: Hydnora sinandevu *Beentje & Q. Luke*.

*Rhizome* irregular, brown or caramel coloured, often both terete, subterete and compressed at different points, rhizome ornamented with numerous lateral tubercles of variable size and shape, randomly and densely distributed on the surface, may occasionally form what appear to be lines. *Flower* usually 4-merous. *Perianth* external tissues coriaceous, brown to distinctly caramel coloured; perianth tissues somewhat fleshy; tepal whitish to yellow-orange inside (in life and dried), tepals narrowly lanceolate, typically often strongly curved, not fused at apex; tepal margins or outer cucullus entirely glabrous. *Osmophore* recessed in tepal cavity, fresh flower not seen by authors, reported to generate faint musty smell or no smell at all.

### RECOGNITION

Hydnora subgenus *Sineseta* is distinguished by having a combination of terete rhizomes with randomly distributed tubercles, entirely glabrous tepals, a recessed osmophore and a lack of an apical osmophore. Only known to parasitise *Commiphora*.

**10. Hydnora sinandevu** *Beentje & Q. Luke* (2002: 5); Bolin et al. (2018: 107). Type: Kenya, Kilifi District, Arabuko-Sokoke, Mida, between mangroves and forest, root parasite on *Commiphora africana* [39°59’E, 03°19’S], 2 m, 1 Jan. 1992, *Luke* 3033a (holotype EA!, isotypes K!, MO!, US!).

Herbaceous subterranean perennial root holoparasite, without leaves or stem. *Rhizome* terete, 0.7–2 cm diam., may be larger on mature specimens (not seen); rhizome surface coriaceous, brown, caramel coloured; rhizome ornamented with numerous lateral tubercles usually distributed in roughly parallel lines on the surface, although sometimes tubercles only loosely follow these lines or not at all, tubercles remain same or develop into flower or develop into haustorium; numerous flowers and flower buds on single rhizomes; rhizome flesh (live) not seen. *Flower* usually 4-merous; flower emerges only partially from the soil; two floral chambers, androecial chamber subtended by gynoecial chamber, inner surfaces of chambers glabrous. *Perianth* external tissues coriaceous, brown to distinctly caramel coloured, total length 8–16 cm; perianth tissues somewhat fleshy; perianth tube reddish to pale pink inside, 5.3–9 cm long × 1.9–2.8 cm wide; tepals whitish to yellow-orange inside (in life and dried), tepal (measured from apex to point of connation with adjacent tepal) 2.7–6.5 cm long × 0.9-2.9 cm wide (measured at midpoint); tepals narrowly lanceolate, typically often strongly curved, not fused at apex; tepal margins or outer cucullus glabrous and not setose. *Osmophore* recessed in tepal cavity, fresh flower not seen by author, reported to generate faint musty smell or no smell at all. *Androecium* antheral ring formed by connate w-shaped anther lobes, forming a central orifice 15–20 mm high, with the top 46-73 mm above the flower base, transversely striate and divided into numerous horizontal pollen sacs. *Gynoecium* ovary inferior, unilocular, subglobose, 15 -30 mm (when dry) in diameter, lobed and cushion-like stigma on floor of gynoecial chamber. *Fruit* only one observed, subterranean, globose, 9 cm in diameter. *Seed* spherical, black-brown, 0.5-2 mm, thousands of seeds within fruit. Fig. 15.

**Fig. 15.**
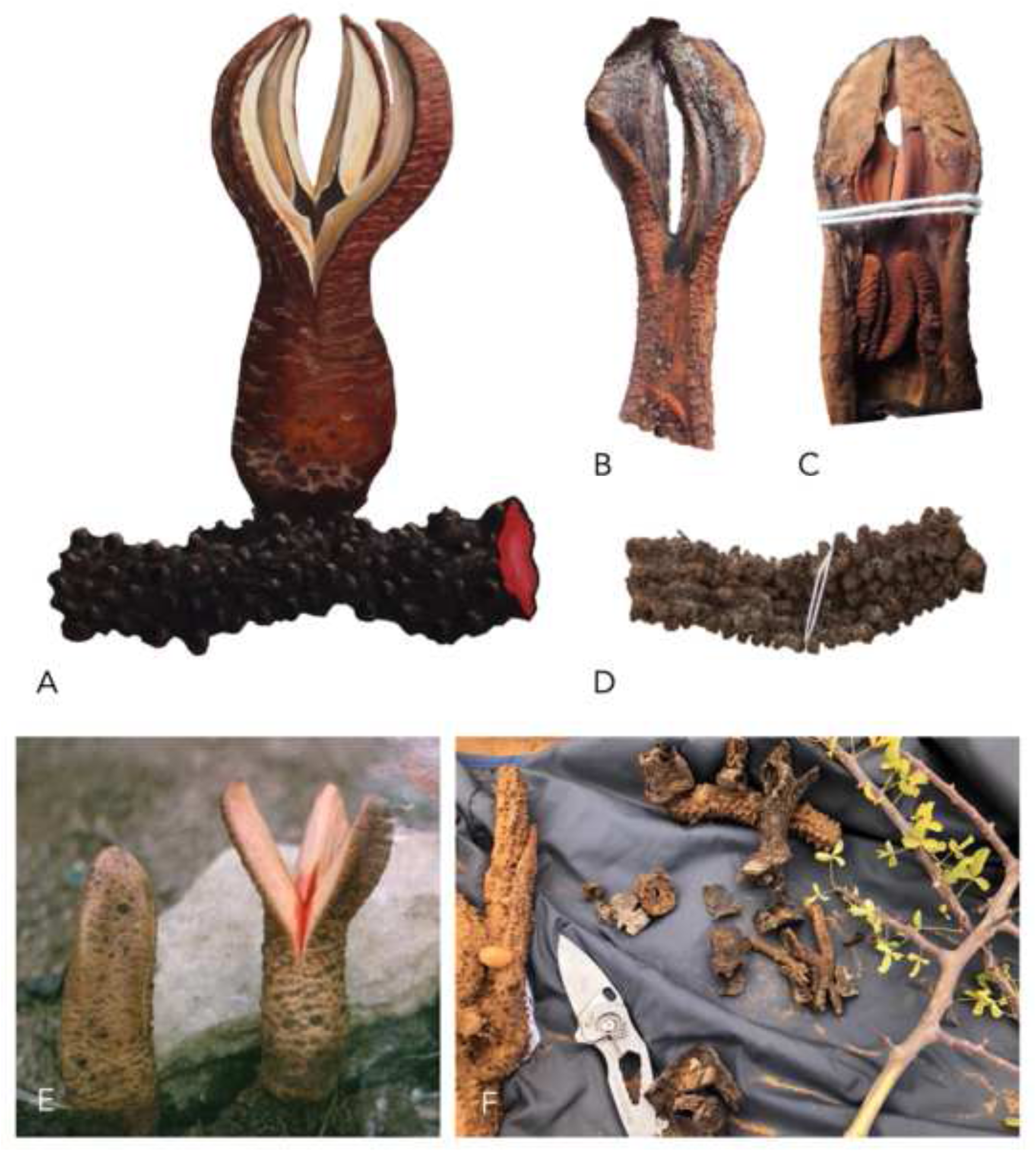
*Hydnora sinandevu*. **A** illustration of flower and rhizome; **B–D** examples of herbarium specimens: **B** flower cross-section (*Luke* 3033A [EA]); **C** young flower cross-section (*Luke* 4304 [EA]); **D** rhizome (*Luke* 3033A [EA]); **E** live bud (left) and open flower (right); **F** cut rhizomes and associated *Commiphora* sp. host (right). **A** ILLUSTRATED BY SEBASTIAN HATT. PHOTOS: **E** WILLIAM BURGER, slide no. 3490 [K].

### DISTRIBUTION

Known primarily from Kenya and Tanzania. A small number of specimens have been collected from Ethiopia, Somalia and Saudi Arabia. This suggests that the distribution of *H. sinandevu* is likely greater than it seems. Collected from an elevation of 0– 1500 m. Map 11.

### REPRESENTATIVE SPECIMENS

**SAUDI ARABIA.** Jizan Prov.: between Sabiya and Idabi, 7 March 1985, *Collenette* 5119 (K!, E!). **SOMALIA.** Northeast Region: Hobyo Distr., 20 km N of Hobyo [5°55.2’N, 48°53.8’E], 30 May 1987, *Wieland* 4422 (MO!); South Region: Lower Shabeelle, Cara Cadde, 1 Aug. 1988, *Kilian & Lobin* 2124 6976 (B!). **ETHIOPIA.** Oromiya Region: Filtù, 18 Oct. 1937, *Vàtova* 814 (FT!); Slopes W of Midaga above Gobelli River Valley, 9 May 1964, *Burger* 3490 (K!, US!); SNNPR: Turkana, a road going to the Ileret Pump [4°29.71’N, 36°22.40’E], 10 June 2012, *Kimeu & Brown* 52 (EA!). **KENYA.** K1: Tana River, S of Garissa, Jan. 1943, *Bally* 2051 (K!, EA!, MO); South Turkana, Kailongoi Mts [1°50’N, 35°50’E], 15 June 1970, *Matthew* 6850 (K!); K7: Kwale, Taru Desert, Oct. 1956, *Bally* 10434 (EA!, MO); Taita in Ost-Afrika, July 1877, *Hildebrandt* s.n. (G!); Kilifi, Arabuko Sokoke, 1 Jan. 1992, *Luke* 3033A (K!, EA!, MO, US!); Tana River Distr., Tana River National Primate Reserve [01°54’S, 040°04’E], 19 March 1990, *Luke et al.* TRP677 (K!, EA!, MO); Tsavo National Park, Voi Hill, 14 May 1963, *Sheldrick 177/63* (EA!). **TANZANIA.** T2: Oldurai Gorge, April 1964, *Leakey* 97/64 (EA!); T3: Lushoto Distr., Tanga, Muheza, 22 Aug. 1935, *Davies* 1004 (K!, EA!, MO); Pangani Distr., Msubugwe, 17 March 1950, *Verdcourt* AH9924 (EA!, MO); T4: Katavi National Park [0706S, 3103E], 18 Dec. 2006, *Luke* 11673 (K [not seen, may be missing], EA!); T6: Pwani, Bagamoyo, 1884, *Kirk* 150 (K [not seen, may be missing], MO!); Selous Game Reserve, Sand Rivers Lodge [07°46’S, 38°08’E], 8 Feb. 1995, *Luke & Luke* 4303 (K!, MO, EA!).

### HABITAT

In Kenya/Tanzania, found in scattered tree grassland, *Acacia-Commiphora* bushland, thicket or the forest margin between mangrove and forest. In Ethiopia it has been found on limestone slopes ∼1500 m under *Terminalia* shrubs/woodland, although most records are from lower altitudes, < 500 m. In Saudi Arabia it has been found in a shallow sandy wadi over basalt slabs, in shade under *Commiphora, Grewia, Maytenus* and *Cissus* shrubs.

### CONSERVATION STATUS

An IUCN red list assessment from 2008 listed *H. sinandevu* as Least Concern (Gereau et al., 2020). It is difficult to speculate on the conservation status of the species, particularly as recent discoveries of specimens in Ethiopia and Saudi Arabia indicate that the range is significantly greater than previously thought.

### PHENOLOGY

Collected in flower in January and August in Kenya/Tanzania, in March in Saudi Arabia and in May in Ethiopia and Somalia. These seemingly random flowering times may indicate that flowering is determined by recent rainfall rather than seasonal changes.

### ETYMOLOGY

The specific epithet translates to ‘without beard’ in Kiswahili. This is a reference to its glabrous tepal margins (Quentin Luke, pers. comm.).

### VERNACULAR NAMES

In northern Kenya it is known by Rendille herdsman as *lekiyomoy* (pers. obs.). Vernacular names for this plant may be the same as for *H. abyssinica*, and it is unknown whether locals differentiate between these two species.

### ETHNOBOTANY

The relatively recent description of this plant means that ethnobotanical information for this species is scant. It is possible that much of the ethnobotanical information for *H. abyssinica* also applies to this species, as the two were conflated in the past. Beentje and Luke (2002) note in the protologue that it has minor medicinal use against throat complaints. Discussions with Rendille herdsman in northern Kenya revealed that they make a drink from the rhizomes that has medicinal value related to birthing (pers. obs.).

### HOSTS

Restricted to *Commiphora* (Burseraceae) hosts, making it the only species of *Hydnora* known to do so (Table 1). The only confirmed hosts are *Commiphora campestris* Engl. and *C. africana* (A.Rich.) Engl. in Kenya and Tanzania. However, *H. sinandevu* in Saudi Arabia was recorded growing amongst several shrubs including *Commiphora quadricincta* Schweinf (Collenette, 1985). Although no direct host-parasite contact was recorded, it seems likely that this was the host plant for this specimen. It may well be the case that populations of *H. sinandevu* discovered in the future are found to parasitise other *Commiphora* species not mentioned here.

### NOTES

Species description adapted from (Beentje & Luke, 2002). *H. sinandevu* was described in 2002 by Beentje & Luke as part of the Flora of Tropical East Africa project. It is easily distinguished by being the only *Hydnora* species to be entirely glabrous and the only species to parasitise *Commiphora* (Burseraceae). It was previously thought to be restricted to Kenya and Tanzania, although collections from Somalia, Ethiopia and Saudi Arabia have been found that were previously misidentified as *H. abyssinica*. This considerably expands the potential distribution of this species. Due to its unique set of characters (see Infrageneric Classification), it has been assigned to a new subgenus, *Sineseta*.

## CONCLUSIONS AND RECOMMENDATIONS FOR FUTURE WORK

This monograph is the first comprehensive taxonomic revision of *Hydnora* for nearly a century, and should serve as a bedrock for future evolutionary and ecological studies into the genus. Following an exhaustive review of herbarium specimens and literature, the number of accepted species is raised from eight to ten, following the reinstatement of *H. solmsiana*, and the description of *H. bolinii* Hatt sp. nov.

While much can be gleaned from existing collections, further field work is critical to better understand all aspects of the biology of *Hydnora*, particularly distribution and host ecology. An important next step will be the production of a widely sampled molecular phylogeny for the genus. This will not only illuminate evolutionary relationships between the species, but also serve as a foundation for future studies. The distribution and host data recorded here has revealed interesting patterns in host specificity. Research into the mechanisms in operation at the host-parasite interface across a range of species will greatly assist our understanding of host specificity and evolution in the genus, and in parasitic plants more broadly.

## ACKNOWLEDGEMENTS

We thank the curators and staff of the following herbaria for hosting the authors, sending loan material or sharing photographs of herbarium specimens: AAU, B, BM, BOL, BR, E, EA, FT, G, GHPG, GRA, HARG, HBG, HUH, JUHN, K, L, LD, LISU, M, MO, NBG, OBG, ODU, P, PRE, PSUB, RO, S, SAM, SRGH, TAN, UPS, US, WIND. We would like to thank the authorities of Namibia (Namibian MET Permit No. RPIV001262022) for permission to collect material. Many thanks to the numerous friends and colleagues who have contributed valuable advice and discussion, including David Goyder, Nina Davies, Erika Maass, Martin Cheek, Gwil Lewis, Oscar Alejandro Pérez Escobar, Iain Darbyshire, Eve Lucas, Mike Gilbert, Alex Zuntini, Quentin Luke, Bat Vorontsova, Gemma Bramley, John Wood, Martin Xanthos, Norbert Holstein, Siro Masinde, Kennedy Matheka, Peter Phillipson, Frances Murray-Hudson, Leevi Nanyeni, Mejo, Quanita Daniels, Rosalia Tshikensho, Hendrina Mutota.

## MAP LEGENDS

**Map. 1.**
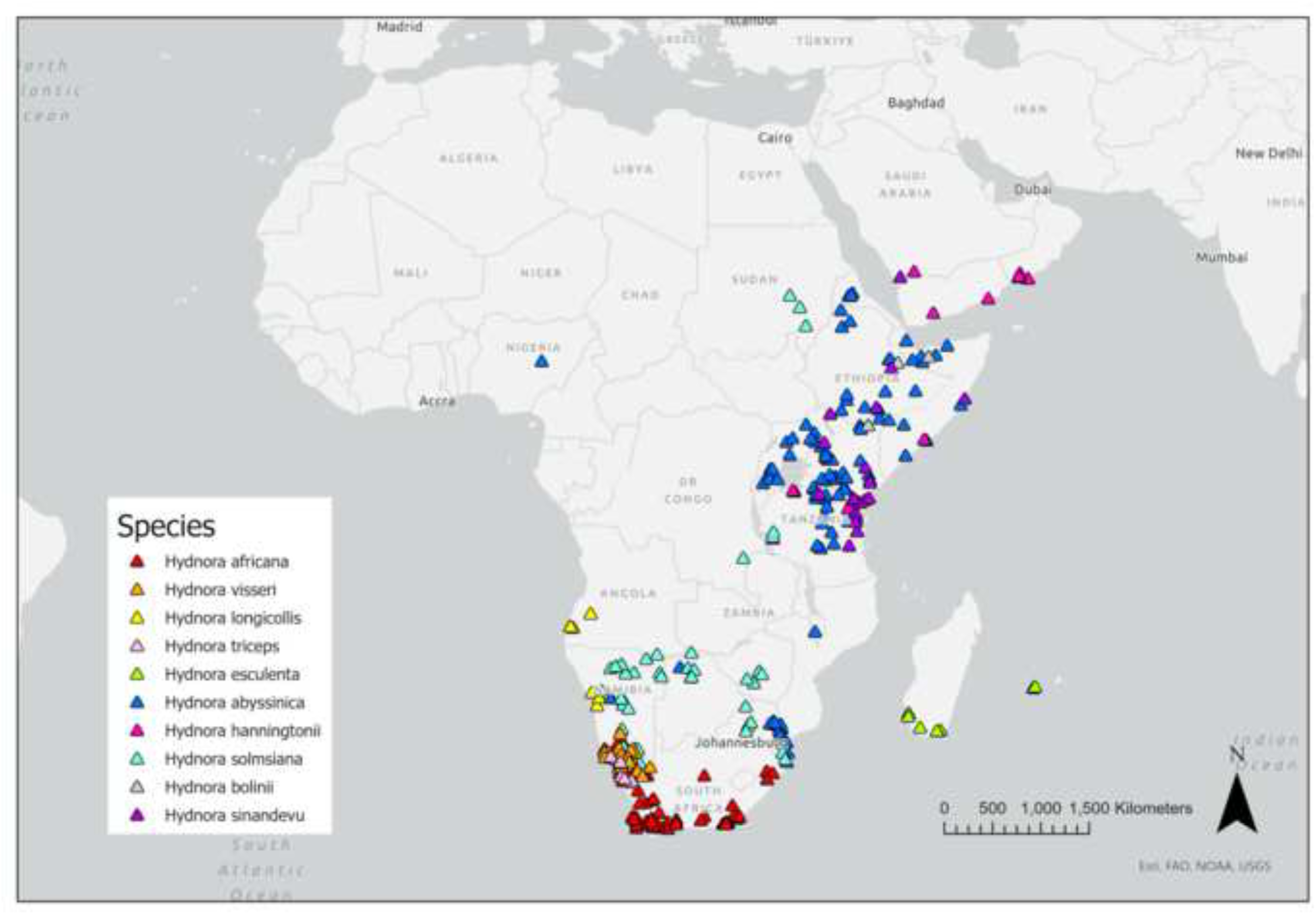
Distribution of all known species of *Hydnora*, compiled from herbarium specimens, iNaturalist records, literature references and personal observations.

**Map. 2.**
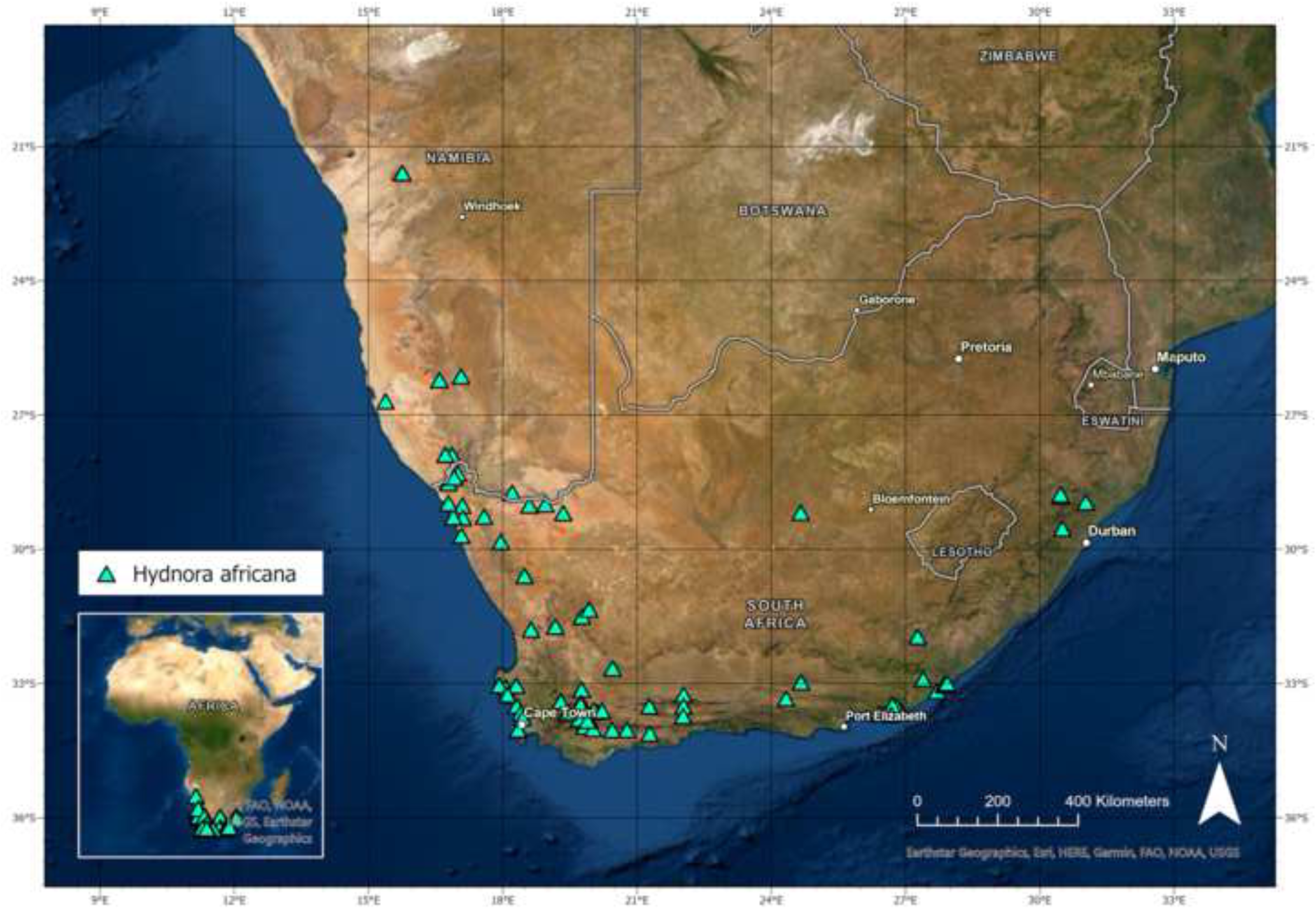
Distribution range of *Hydnora africana*.

**Map. 3.**
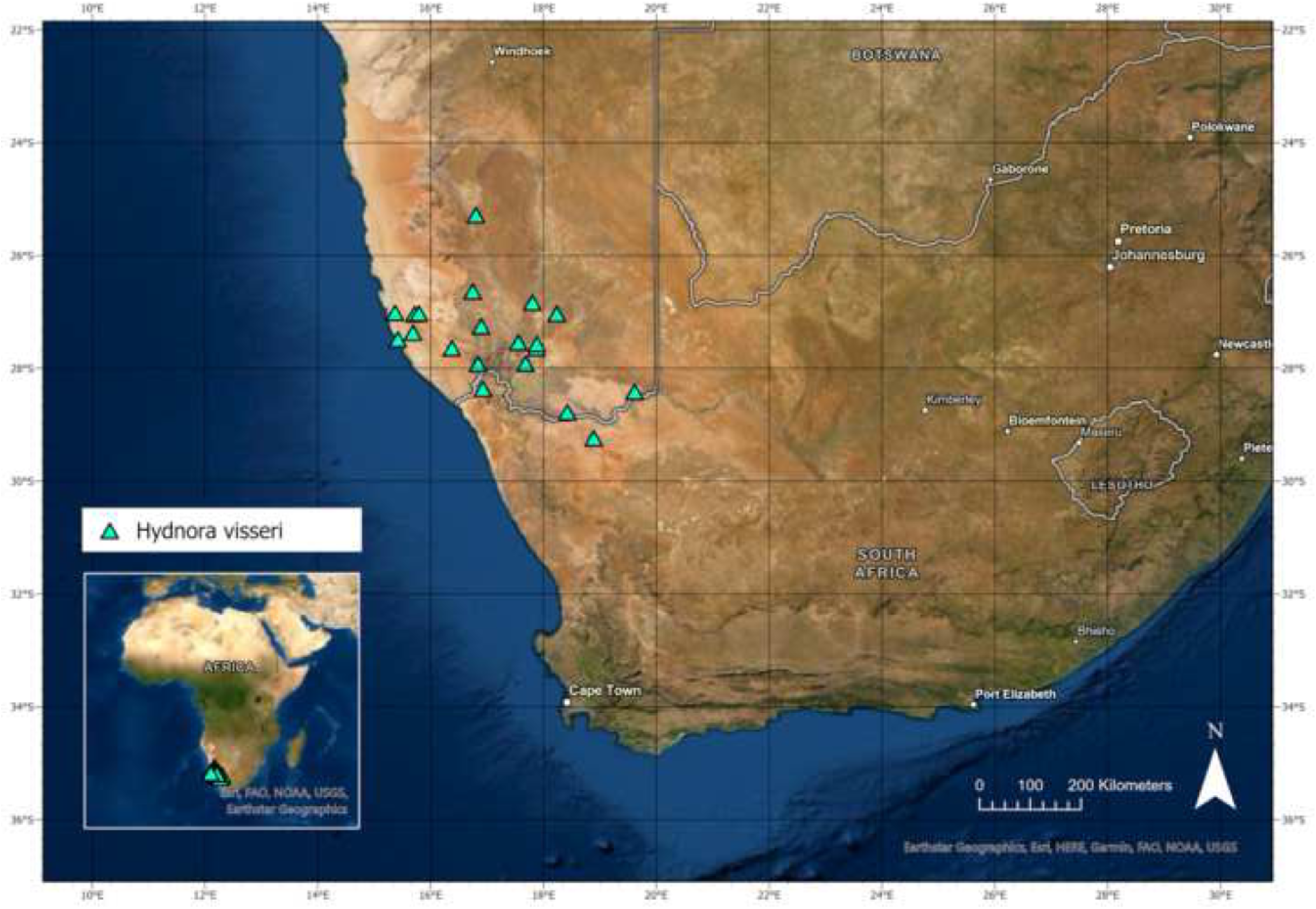
Distribution range of *Hydnora visseri*.

**Map. 4.**
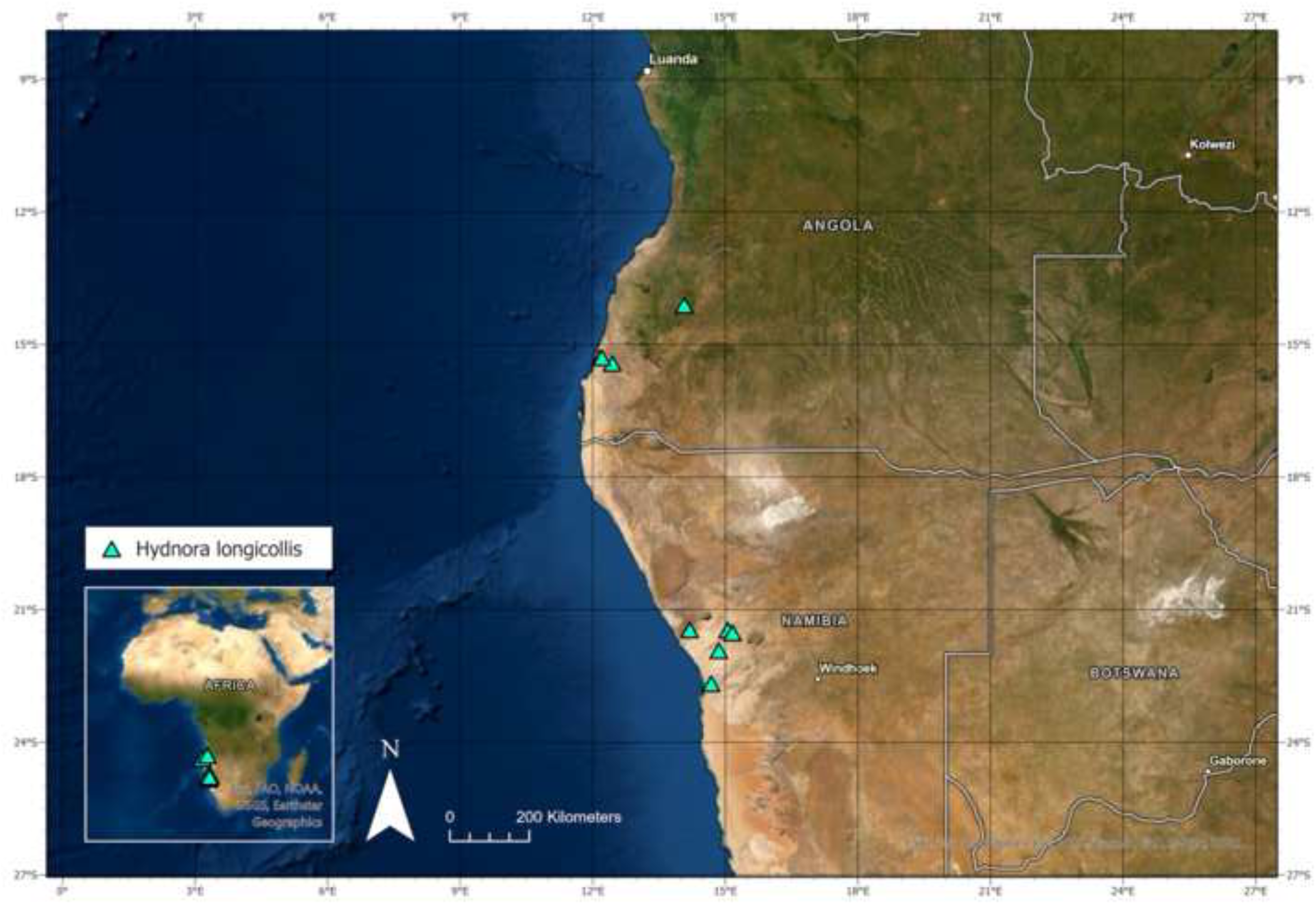
Distribution range of *Hydnora longicollis*.

**Map. 5.**
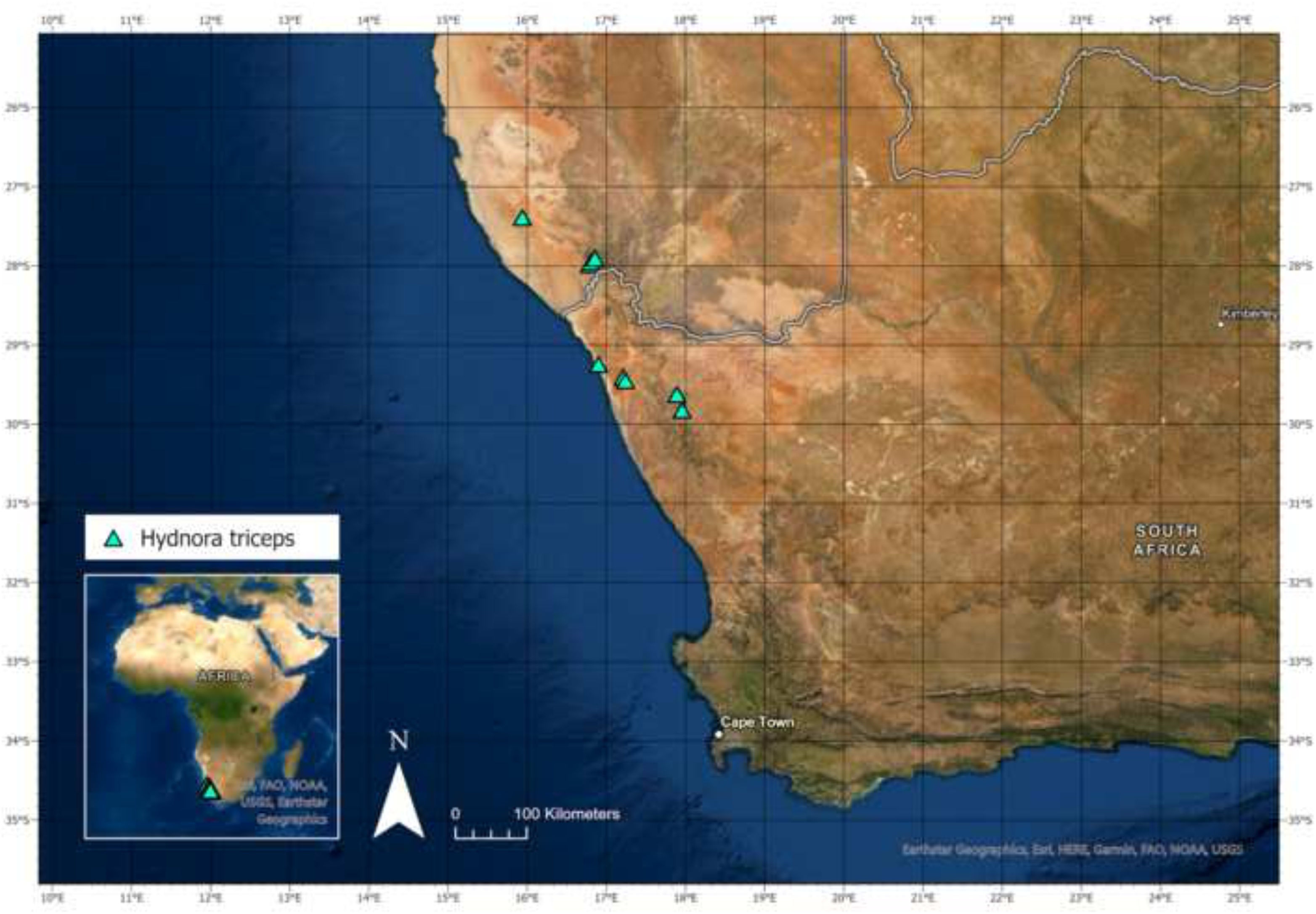
Distribution range of *Hydnora triceps*.

**Map. 6.**
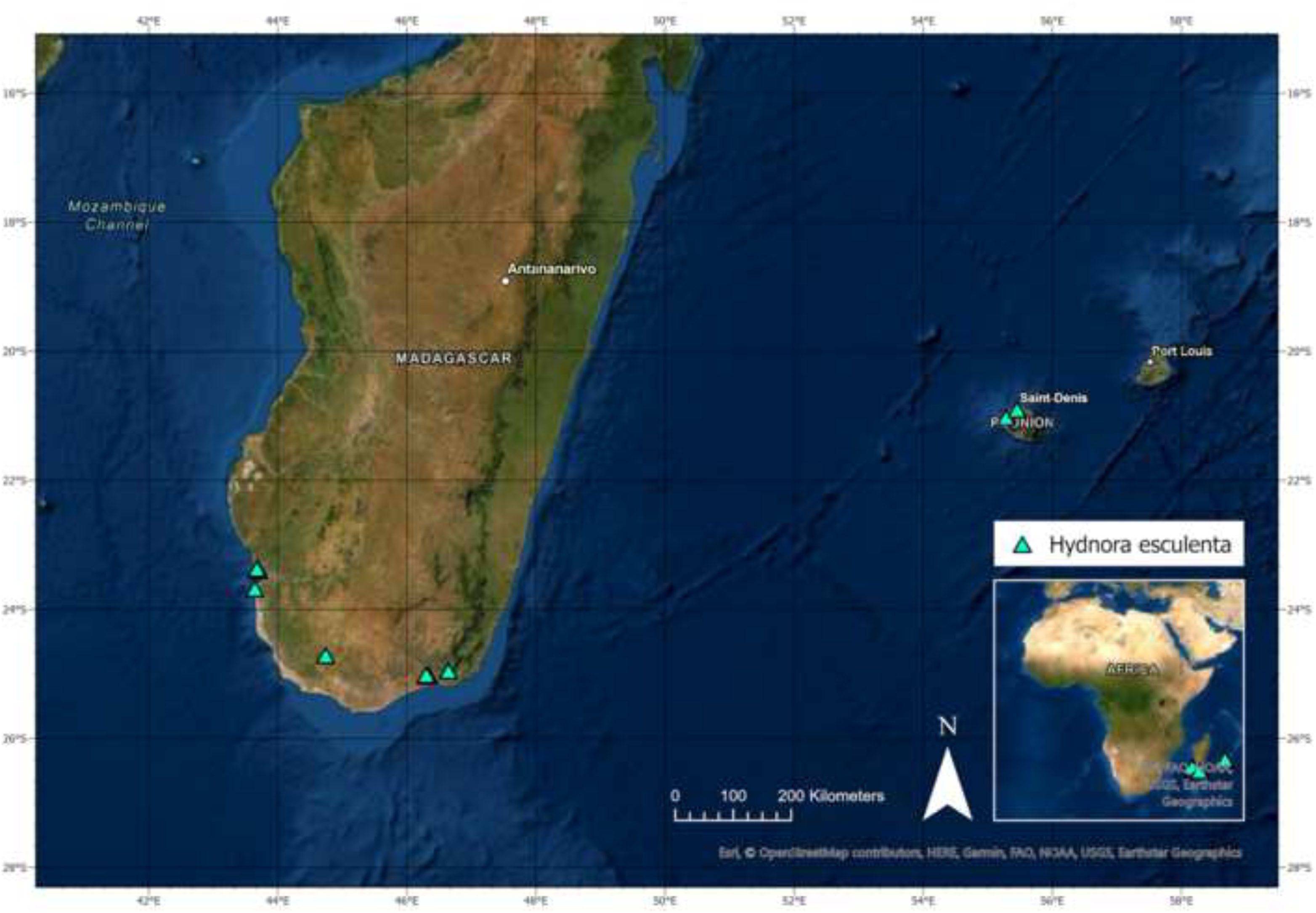
Distribution range of *Hydnora esculenta*.

**Map. 7.**
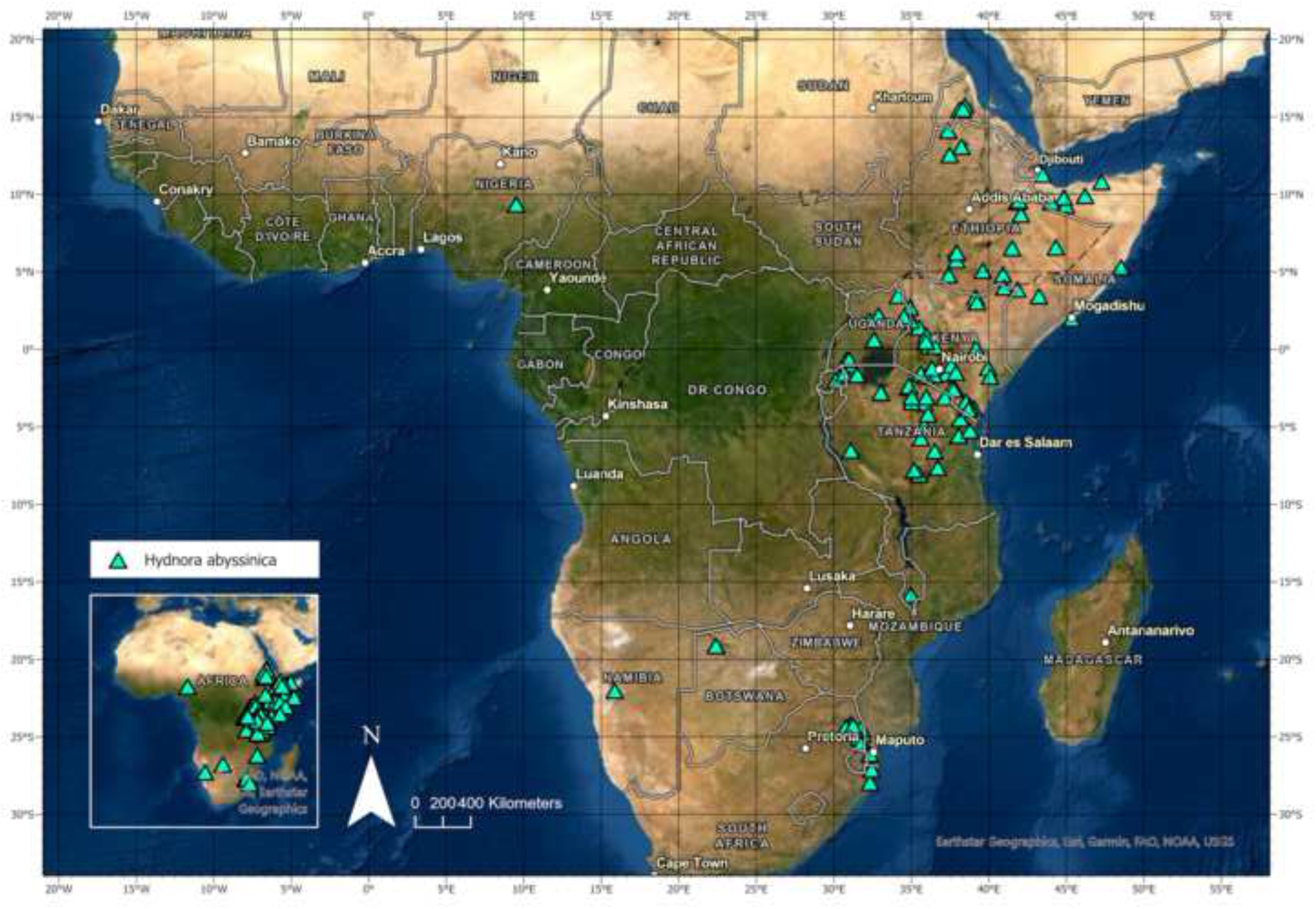
Distribution range of *Hydnora abyssinica*.

**Map. 8.**
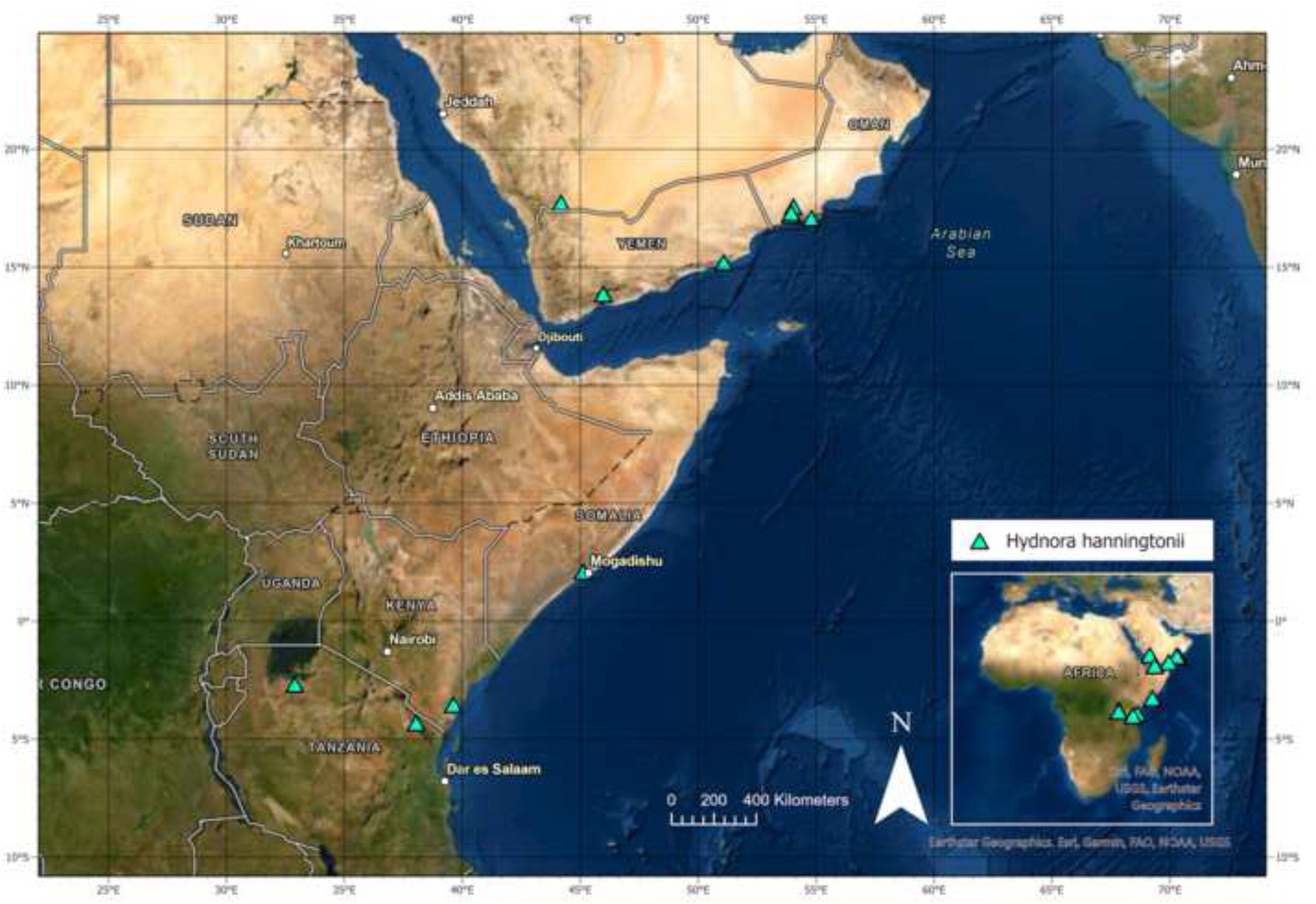
Distribution range of *Hydnora hanningtonii*.

**Map. 9.**
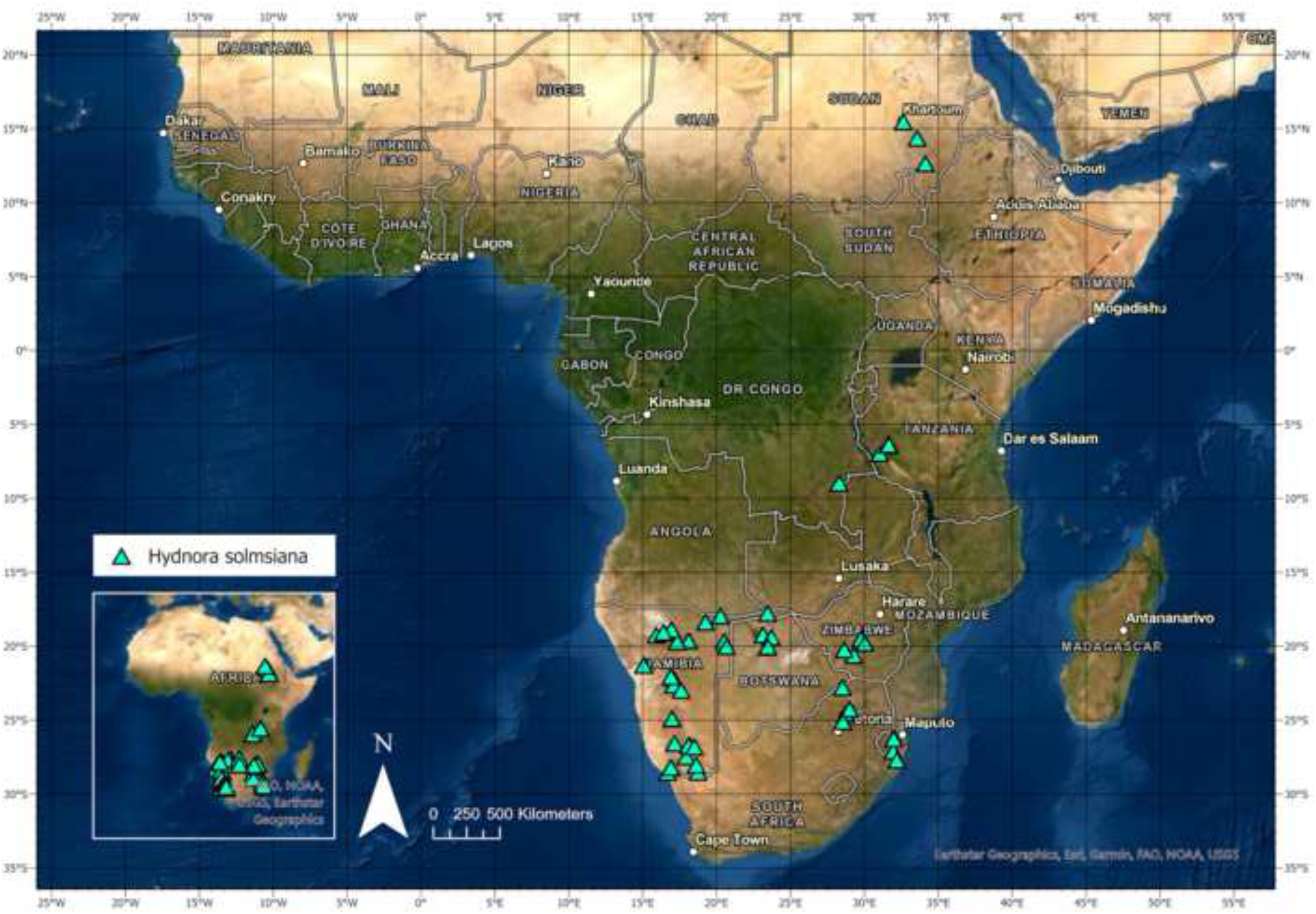
Distribution range of *Hydnora solmsiana*.

**Map. 10.**
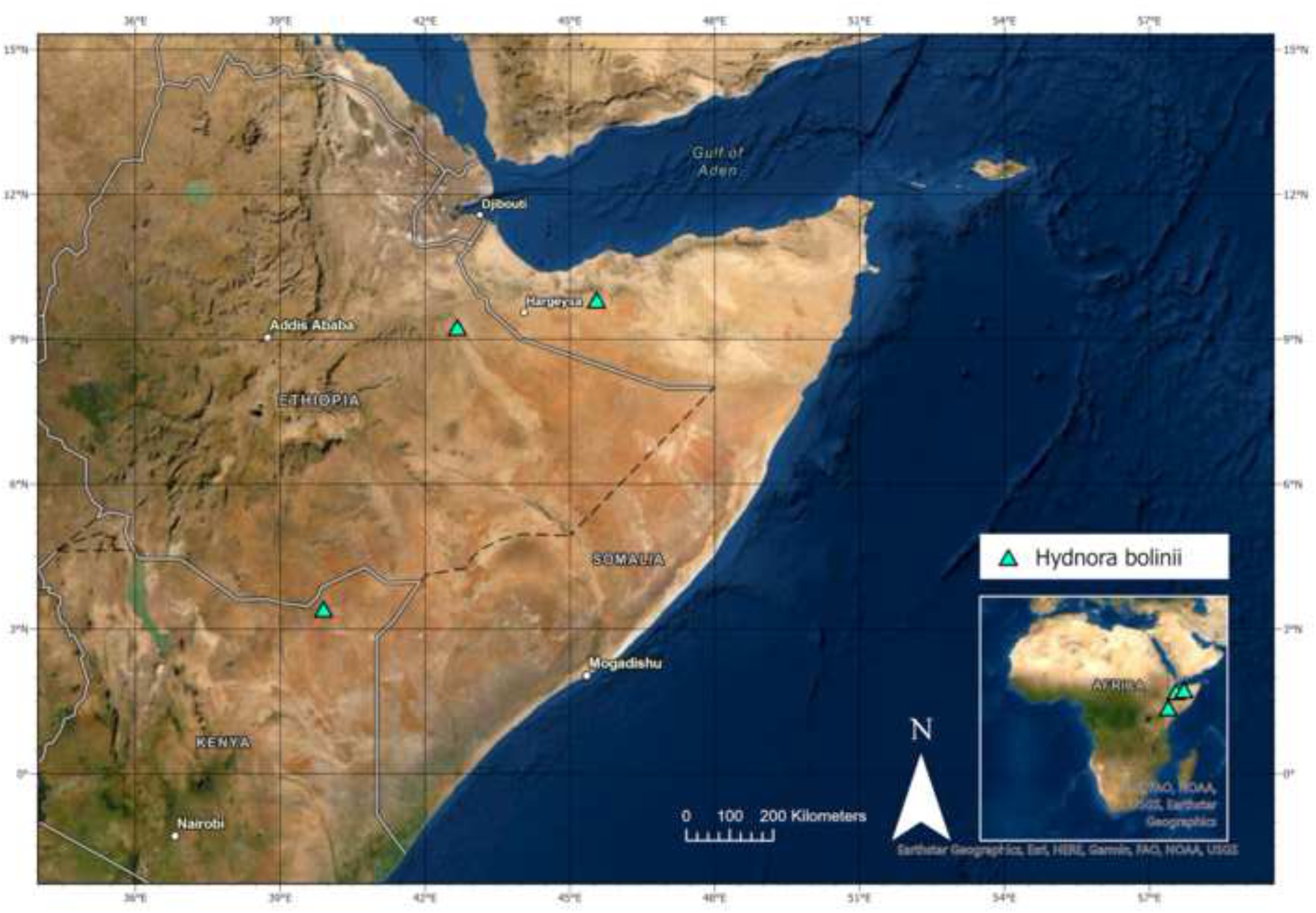
Distribution range of *Hydnora bolinii*.

**Map. 11.**
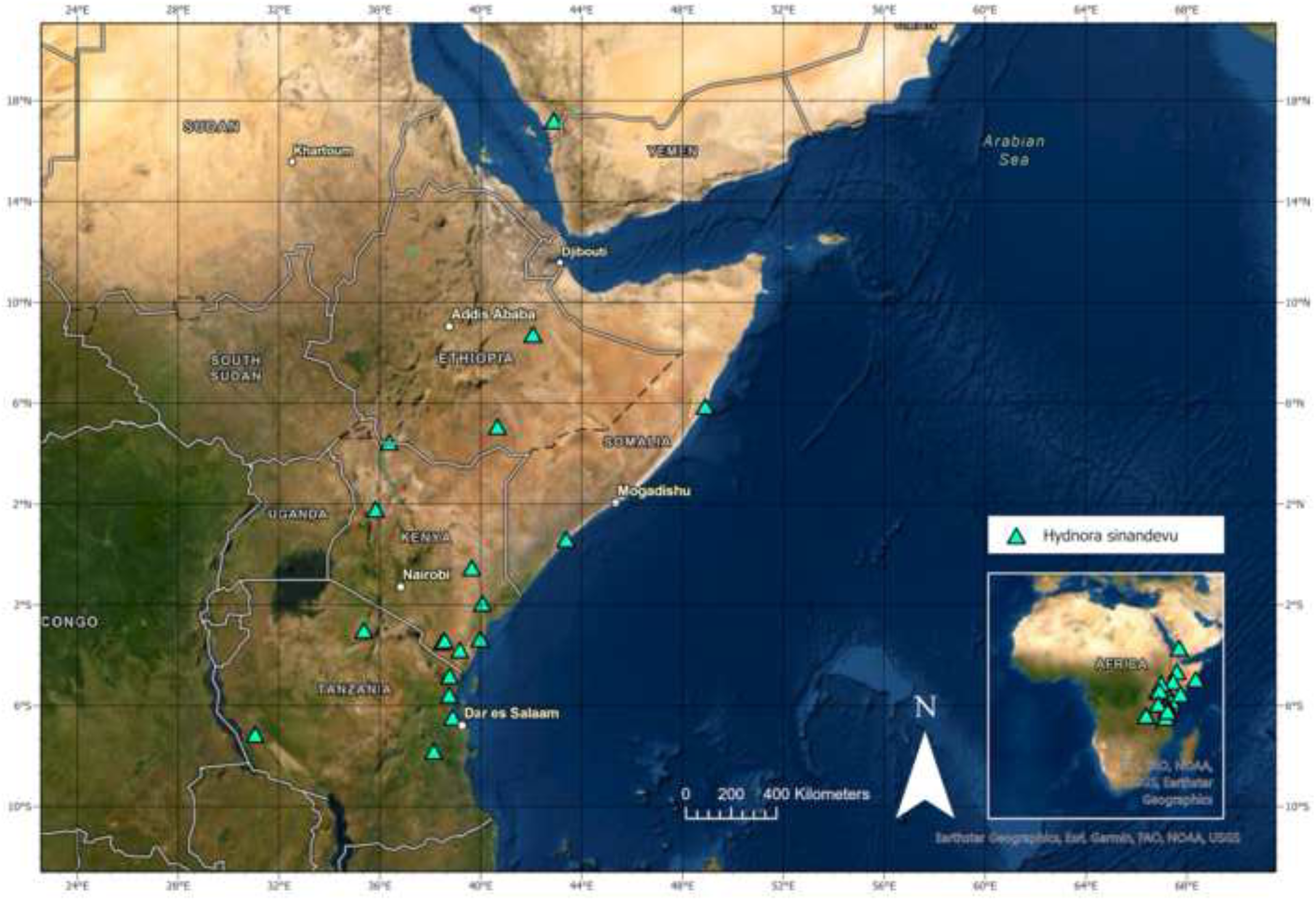
Distribution range of *Hydnora sinandevu*.

